# Synthesis and Structure Elucidation of the Human tRNA Nucleoside Mannosyl-Queuosine

**DOI:** 10.1101/2021.06.24.449707

**Authors:** Markus Hillmeier, Mirko Wagner, Timm Ensfelder, Eva Korytiakova, Peter Thumbs, Markus Müller, Thomas Carell

## Abstract

Queuosine (Q) is a structurally complex, non-canonical RNA nucleoside. It is present in many eukaryotic and bacterial species, where it is part of the anticodon loop of certain tRNAs. In higher vertebrates, including humans, two further modified queuosine-derivatives exist - galactosyl-(galQ) and mannosyl-queuosine (manQ). The function of these low abundant hypermodified RNA nucleosides remains unknown. While the structure of galQ was elucidated and confirmed by total synthesis, the reported structure of manQ still awaits confirmation. By combining total synthesis and LC-MS-co-injection, together with a metabolic feeding study of labelled hexoses, we show here that the natural compound manQ isolated from mouse liver deviates from the literature-reported structure. The chemical structure of the natural product manQ features a novel α-*allyl* connectivity. The data reported here shows that the yet unidentified glycosylases that attach galactose and mannose to the Q-base have different constitutional connectivity preferences. Knowing the correct structure of manQ will now pave the way towards further elucidation of its biological function.

## Introduction

Ribonucleic acid (RNA) is a central molecule of life, linking the genotype to the phenotype by integrating both catalytic and coding properties in the synthesis of proteins.^1,2^ To fulfil the plethora of functions known for RNA today, a huge chemical diversity has developed regarding the nucleobases through the evolution of life.^3,4^ The most densely modified RNA molecules are the tRNAs.^5^ Those small adapter molecules feed the required amino acids to the growing peptide chain in the ribosome. Of particular interest are the non-canonical nucleosides that are found in the anticodon loop of tRNAs, in particular at position 34, known as the Wobble position of the anticodon.^6^ Here, any chemical modification has a direct impact on the coding potential of the tRNA.^6-9^ Queuosine (**1**, Q, **Fig. 1a**) is a particularly complex non-canonical nucleoside.^10-12^ It is found in many bacterial and eukaryotic species, where it is located at position 34 of the anticodon loop of tRNA^Tyr^, tRNA^Asp^, tRNA^His^, and tRNA^Asn^.^13,14^

**Figure 1.**
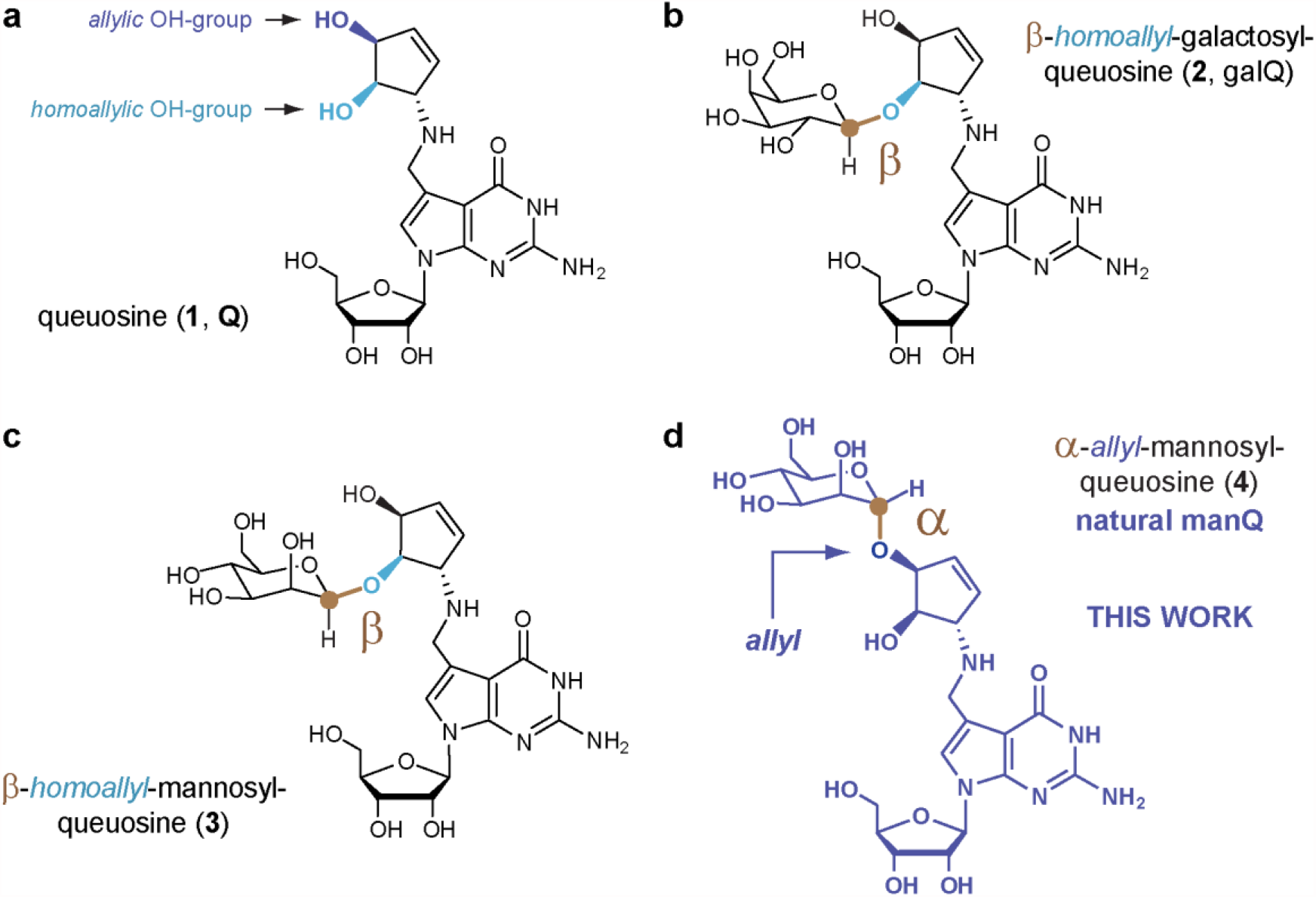
Nucleosides of the queuosine family present in higher vertebrates. **a**, Queuosine **1** is derived from a 7-deazaguanosine base with a 1(*S*)-amino-2(*R*),3(*S*)-dihydroxycyclopent-4-ene unit attached *via* a methylene linker to *C*7. Possible glycosylation sites are depicted in dark blue (*allylic* hydroxyl group) and light blue (*homoallylic* hydroxyl group); **b**, Structure of the naturally occurring nucleoside β-*homoallyl*-galQ **2**; **c**, Structure **3** shows the originally proposed structure of manQ, which we revise in this work; **d**, Corrected structure of natural manQ **4** featuring an α-*allylic* connectivity.

In eukaryotes, queuosine **1** is found in both cytosolic and mitochondrial tRNA.^15^ It is biochemically derived from guanosine (G) and characterized by the exchange of the *N7*-nitrogen for a *C7*-carbon atom and the addition of a 1(*S*)-amino-2(*R*),3(*S*)-dihydroxycyclopent-4-ene unit to the *C7*-position via a methylene linker.^12,16^ The presence of the Q-base in position 34 of the anticodon equips the corresponding tRNAs with the possibility to decode synonymous codons by wobble base pairing. ^17,18^ The Q-base is furthermore affecting translational speed,^19^ decoding fidelity,^20^ and it is essential for survival particularly in the absence of sufficient tyrosine.^21,22^ Eukaryotes are unable to biosynthesize Q, which forces them to acquire it from prokaryotic sources.^21,23-25^ This establishes a link between the central process of translation and the gut microbiome.^26^ Defective RNA modification processes are increasingly recognized as drivers for severe diseases. ^27,28^

In higher vertebrates, including humans, two glycosylated queuosine derivatives exist, which have yet unidentified functions. The first of these hypermodified Q derivatives is found in cytosolic tRNA^Tyr^, where the Q-base is modified with a galactose residue (galQ, **2, Fig. 1b**).^29,10^ The cytosolic tRNA^Asp^, in contrast, contains a Q-derivative, which is suggested to be mannosylated (manQ, **Fig. 1c, d**).^29^ Here, we report the total synthesis of the “human natural product” manQ and show that the originally proposed structure **3** (**Fig. 1c**) needs to be revised in two important aspects: In contrast to the original structure suggestion, we show that the mannose glycosidic bond is not β- but α-configured. Furthermore, and in contrast to β-galQ, the mannose is attached not to the *homoallylic*, but to the *allylic* hydroxyl group at *C3*. manQ has consequently the structure **4** shown in **Fig. 1d**. This result reveals that the yet unknown galactosyl and mannosyl transferases that attach the respective hexose to the Q-nucleobase in tRNA^Tyr^ and tRNA^Asp^, are able to differentiate the two hydroxyl groups (*allyl* vs. *homoallyl*) at the cyclopentene substructure for yet unknown reasons.

## Main Text

### Synthesis of literature-reported manQ

Because the molecules galQ and manQ are found only in the anticodon loop of one cytosolic tRNA each,^30^ the amounts that can be isolated from nature (e.g. porcine liver) are so small that a structure elucidation based solely on isolated material is practically impossible. Therefore, synthesis of the proposed manQ compound and comparison of the synthetic standard with natural material by LC-MS was the method of choice for us to investigate the manQ structure. The synthesis of compound **3**, proposed to be natural manQ, was performed as depicted in **Fig. 2a**.

**Figure 2.**
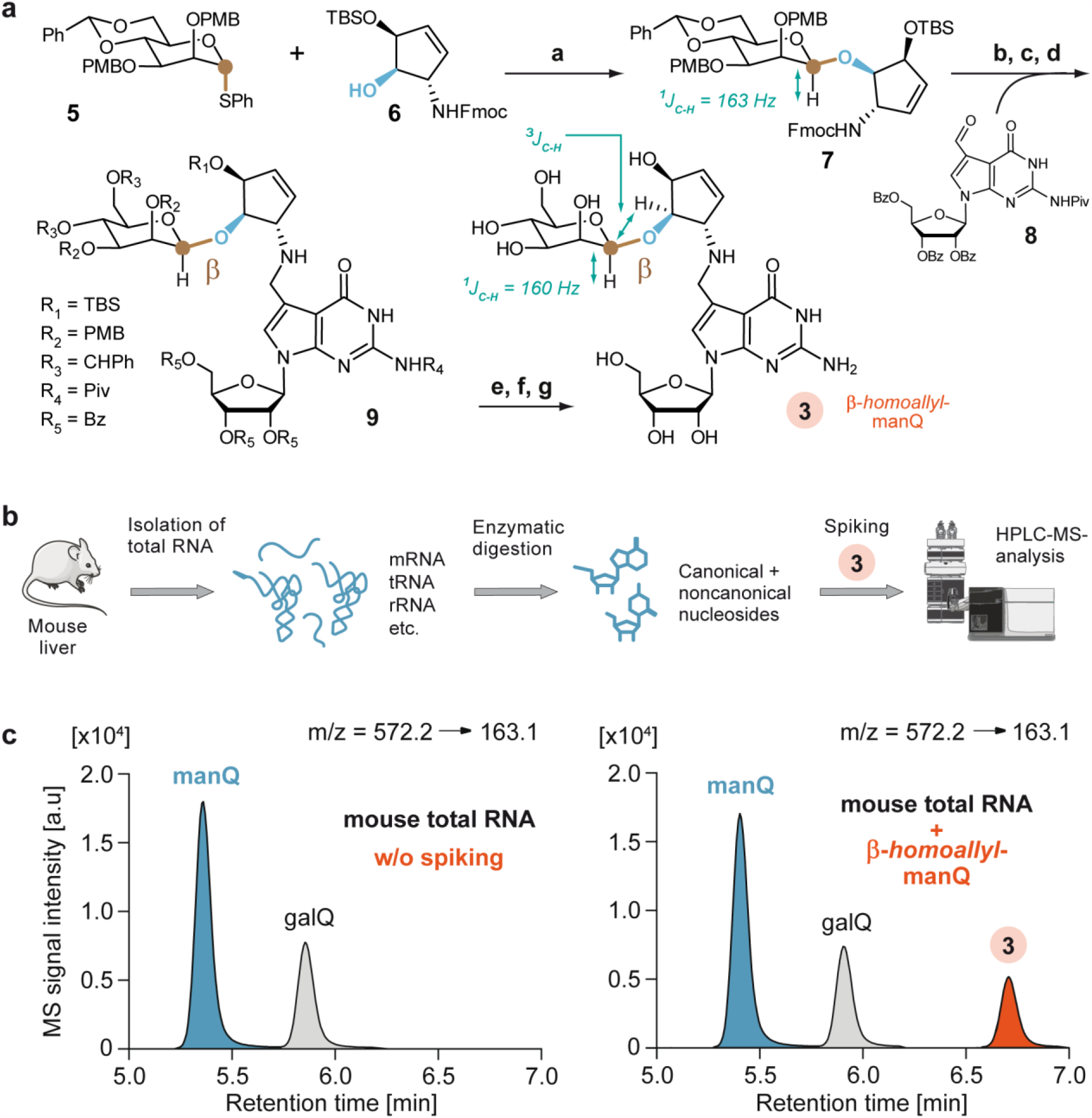
Attempt to confirm the originally proposed structure of manQ. **a**, Synthetic route towards β-*homoallyl*-manQ featuring a β-selective mannosylation key step; a: AgOTf, PhSCl, DTBMP, DCM, -78 °C, 2 h, 62 %; b: HNMe2, THF, rt, 1 h; c: **8**, MeOH, rt, 5 h; d: NaBH4, MeOH, 0 °C, 1 h; e: NaOMe, MeOH, rt, 3 h; f: TFA, DCM, 0 °C, 20 min; g: HF · pyridine, pyridine, rt, 18 h, 22 % (5 steps); **b**, Workflow for the comparison of the synthetic β-*homoallyl*-manQ standard with natural manQ *via* isolation of total RNA from mouse liver, digestion to the nucleoside level, and LC-MS-co-injection experiments; **c**, Chromatograms resulting from the co-injection experiments analysed by UHPLC-MS/MS; Left: Digested total RNA from mouse liver shows two peaks, natural manQ and galQ.; Right: Digested total RNA from mouse liver spiked with the synthetic standard β-*homoallyl*-manQ **3** shows the appearance of an additional peak, thereby disproving the originally proposed structure for manQ.

For the synthesis of the difficult-to-access β-mannosyl connectivity, we employed the *Crich*-method^31^, which required the preparation of the mannosyl donor **5**. This was achieved along the literature-reported synthesis.^31,32^ The mannosylation reaction of the Fmoc- and TBS-protected 1(*S*)-amino-2(*R*),3(*S*)-dihydroxycyclopent-4-ene unit **6** with the mannosyl donor **5** required an activation with Ag-triflate in the presence of phenylthiochloride and 2,6-di-*tert*-butyl-4-methylpyridine (DTBMP) at -78 °C in dry dichloromethane. The reaction gave the mannoside **7** in 62 % yield, and the β-configuration of its anomeric centre was confirmed by NMR-spectroscopy based on the coupling constant of ^1^J_C1-H1_ = 163 Hz, which is typical for β-mannosides.^33^ We next cleaved the Fmoc group of the mannoside **7** with dimethylamine (10 % DMA in THF, rt, 1 h) and performed the reductive amination with the benzoyl- and pivaloyl-protected 7-formyl-7-deazaguanosine compound **8** (MeOH, rt, 5 h, then NaBH_4_, 0 °C, 1 h) that was prepared as recently described by us.^10^ This afforded the fully protected manQ compound **9**. Cleavage of the benzoyl-protecting groups at the ribose was performed under *Zemplén*-conditions (0.5 M NaOMe, MeOH, rt, 5 h). The PMB-ethers were deprotected with trifluoro acetic acid in dichloromethane (10 %, 0 °C, 20 min). Finally, we removed the TBS-protecting groups with HF in pyridine (rt, 18 h). This furnished the final manQ compound **3** with mannose being β-configured and attached to the *homoallylic* hydroxyl group. As expected, compound **3** has a ^1^J_C-H_-coupling constant of 160 Hz at the anomeric centre, thereby proving its β-configuration. Furthermore, the observed ^3^J_C-H_-HMBC coupling in compound **3** between the mannosyl *C1*-carbon and the *homoallylic* hydrogen *C4*H of the cyclopentene ring (see Supplementary Information 4) unambiguously proved that the mannosyl residue is indeed connected to the *homoallylic* hydroxyl group.

In order to compare the synthetic manQ compound **3** with the natural material, we performed an LC-MS co-injection study. To this end, we isolated the total RNA from mouse liver and enzymatically digested this RNA down to the individual nucleosides as recently described by us.^10^ This procedure afforded a mixture of all nucleosides present in the total RNA pool. Subsequent analysis of the resulting nucleoside mixture was performed *via* UHPLC-MS/MS with a triple quadrupole mass spectrometer (QQQ) set to monitor a specific molecular fragmentation reaction of the glycosylated Q-derivatives. Using collision-induced dissociation, these nucleosides undergo two heterolytic bond dissociations, leading to a loss of both the ribose sugar and the glycosylated cyclopentene unit. Monitoring the corresponding mass transition of m/z = 572.2 → 163.1 allows a sensitive detection of these compounds (see **Fig. 2b** and Supplementary Figure 1). With an UHPLC separation gradient of 0→2 % (v/v) MeCN/H_2_O in 0→8 min on a *Poroshell 120 SB-C8* column, all hexose-modified queuosine derivatives can be separated chromatographically (see **Fig. 2c**). Indeed, when analysing the mouse liver sample, we detected two clearly separated signals in the RNA nucleoside pool with galQ **2** eluting at 5.9 min, and the natural manQ compound eluting at 5.4 min, respectively. The compound eluting at 5.9 min was unambiguously identified as galQ **2** with the help of a synthetic reference compound (Supporting Fig. 2). Upon co-injection of the synthetic material **3**, we were expecting to see again two signals with the manQ signal having gained in intensity. To our surprise, however, we noted that this is not the case. Instead, the co-injection experiment provided a third signal with a retention time of 6.8 min, clearly separated from both natural manQ and galQ **2**. This result unequivocally shows that manQ **3** is NOT identical to the natural manQ compound. Therefore, the structure proposal for manQ reported in literature must be incorrect.

Puzzled by this result, we initially reasoned that the compound originally identified as manQ may contain a different sugar than mannose, even though a previous study showed that a mannosyl moiety can enzymatically be transferred to the Q-base from GDP-mannose.^34^ To verify that manQ contains indeed a mannose sugar, we performed a metabolic labelling study (**Fig. 3a**): We added different isotope-labelled sugars to a HEK293T cell culture and subsequently isolated the total RNA of the cells to see if the natural manQ had incorporated the isotope labels. During these experiments, the cell culture medium was additionally supplemented with high concentrations of unlabelled glucose or mannose as carbon source in order to suppress the metabolic conversion of the labelled sugar into other carbohydrates. Without these metabolic suppressors, such conversion would initiate an isotope scrambling process that would jeopardize the experiments. The results of our study are depicted in **Figure 3b**. It is clearly evident that feeding of ^13^C_6_-galactose and ^13^C_6_-glucose in the presence of the metabolic suppressors glucose or mannose, respectively, gave little or no incorporation of ^13^C into the isolated natural manQ compound. In contrast, feeding of ^13^C_6_-mannose in combination with the metabolic suppressor glucose quickly led to the time-dependent formation of ^13^C_6_-manQ, thereby confirming that the sugar connected to the Q-base in manQ is indeed mannose.

**Figure 3.**
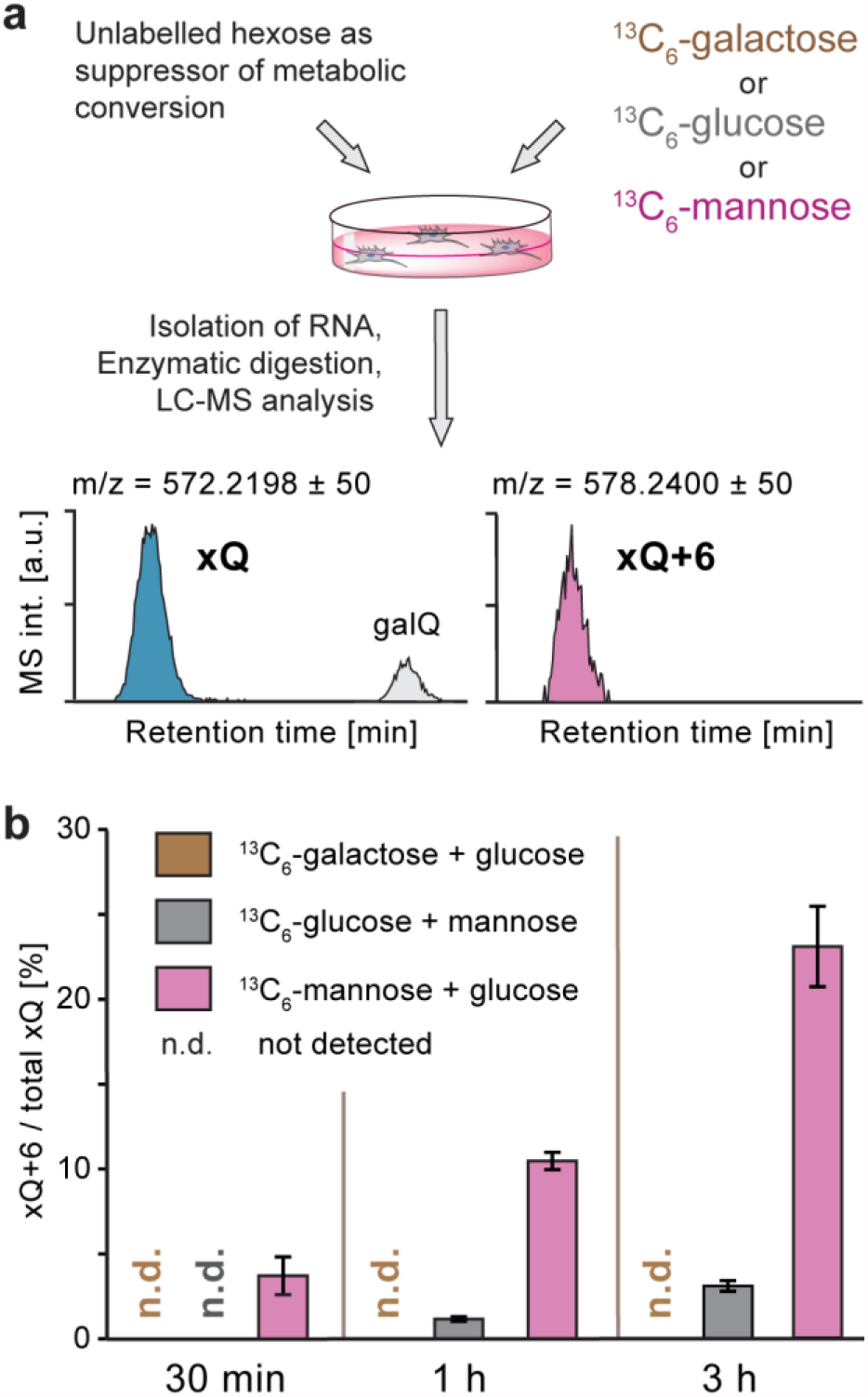
Confirmation of the incorporated hexose sugar as being mannose. **a**, Workflow of the metabolic labelling study providing ^13^C_6_-galactose, ^13^C_6_-glucose, or ^13^C_6_-mannose in combination with an unlabelled carbon source (glucose or mannose) that suppresses metabolic interconversion of the carbohydrates and hence isotope scrambling; **b**, Results of the LC-MS experiments showing quick incorporation of heavy sugar when feeding ^13^C_6_-mannose in presence of glucose, thereby confirming the identity of xQ as manQ; Error bars represent the standard deviation of three independent experimental replicates

We therefore reasoned that natural manQ differs from the reported structure **3** likely regarding the configuration of the anomeric centre and/or the connectivity at the cyclopentene moiety (*allyl* versus *homoallyl* mannoside). With **3** proven not to be the natural compound, we were left with the three remaining structures **4** (α-*allyl*-connectivity), **10** (β-*allyl*-connectivity), and **11** (α-*homoallyl*-connectivity, see **Fig. 4a**). For their synthesis, it was necessary to first develop syntheses for the properly protected 1-amino-2,3-dihydroxycyclopent-4-enes **13**-**15** (**Fig. 4b**). To access these compounds, we started with the Fmoc-protected 1-amino-2,3-dihydroxypent-4-ene **12**. For the synthesis of the 1-Fmoc-3-PMB-protected cyclopent-4-ene **13**, needed for the synthesis of **11**, we first protected in **12** both OH groups as an anisaldehyde acetal (anisaldehyde dimethylacetal, CSA, rt, 2 h, 92 %), followed by a selective reductive opening of the *homoallylic* OH group with DIBAL-H. This gave the 1-Fmoc-3-PMB-protected-dihydroxycyclopent-4-ene **13** (DCM, -78 °C, 2 h, 85 %). For **14**, we used **13** as the starting material. Protection of the *homoallylic* hydroxyl group in **13** with SEM-Cl (NBu_4_I, pyridine, DMF, 70 °C, 18 h, 83 %), followed by cleavage of the *p*-methoxybenzylether (10 % TFA in DCM, 0 °C, 5 min) furnished compound **14**. Reaction of **12** with TBSOTf at -55 °C provided after 15 min predominantly the 1-Fmoc-3-TBS-protected compound **6** as the kinetic product, which we had used before for the synthesis of the literature-reported structure of manQ **3**. If this reaction was, however, performed at -10 °C for a longer period of time (2 h), we observed formation of the thermodynamically more stable 1-Fmoc-2-TBS-protected compound **15**.

**Figure 4.**
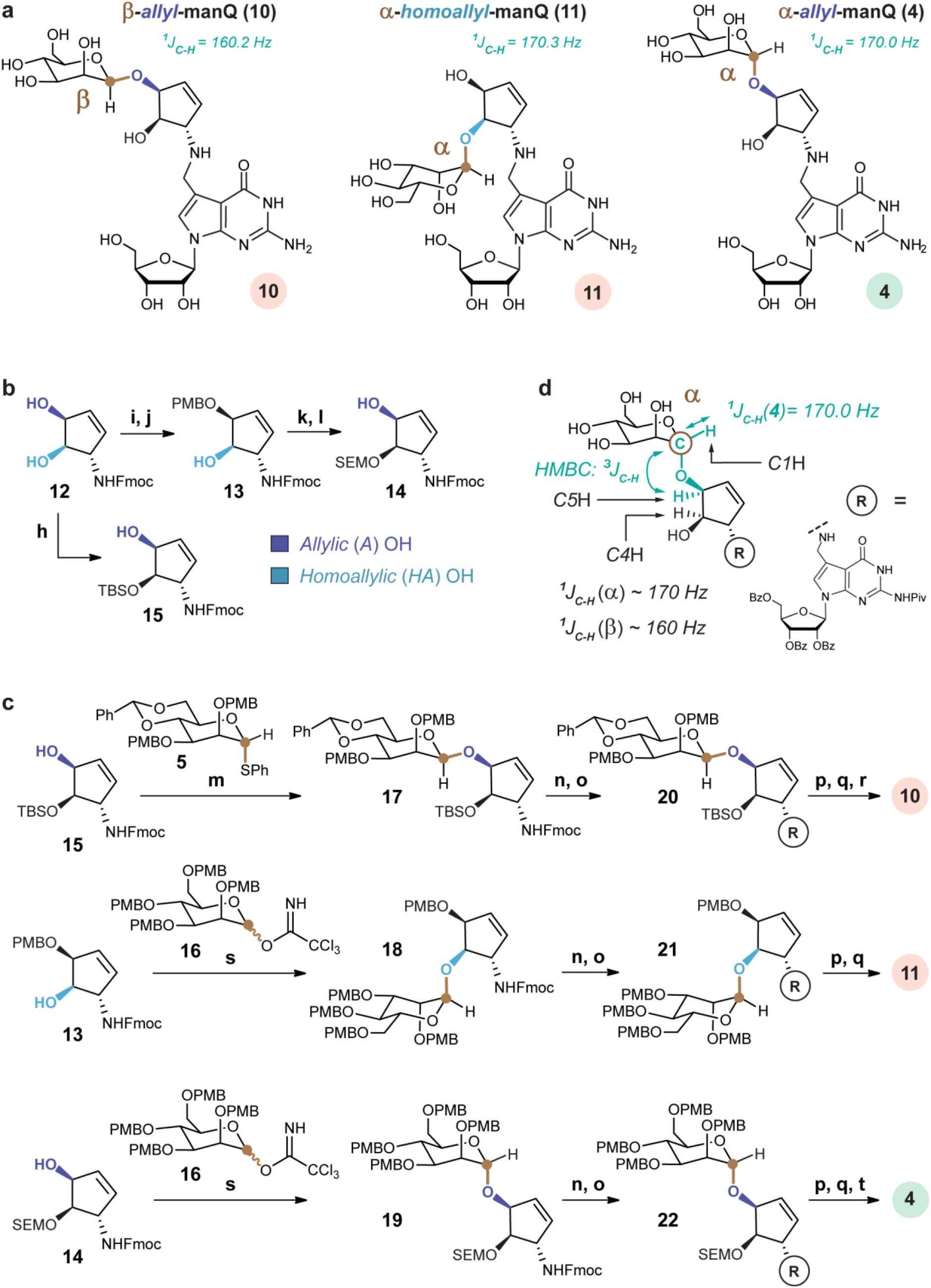
Alternative manQ structures and their syntheses. **a**, manQ target compounds: β-*allyl*-manQ **10**, α-*homoallyl*-manQ **11**, and the correct structure **4** of natural manQ featuring an α-*allyl*-connectivity; **b**, Regioselective protection of cyclopentene **12** for the synthesis of precursors **13**-**15**; h: TBSOTf, DMF, -10 °C, 2 h, 41 %; i: anisaldehyde dimethyl acetal, CSA, DMF, rt, 2 h, 98 %; j: DIBAL-H, DCM, -78 °C, 3 h, 85 %; k: SEM-Cl, NBu4I, pyridine, DMF, 70 °C, 18 h, 75 %; l: TFA, DCM, 0 °C, 5 min, 83 %; **c**, Synthesis of the manQ compounds **10, 11**, and **4** from **13, 14** and **15** *via* an appropriate stereoselective glycosylation step and subsequent reductive amination; m: AgOTf, PhSCl, DTBMP, DCM, -78 °C, 2 h, 52 %; n: DBU, MeCN, rt, 1 h; o: **8**, MeOH, 2-5 h, then NaBH4, 0 °C, 15-30 min; p: NaOMe, MeOH, rt; q: HF · pyridine, EtOAc, rt, 18 h; r: TFA, DCM, 0 °C; s: TMSOTf, DCM, 0 °C; t: HF · pyridine, MeCN, rt, 1 week; **d**, Depiction of the ^*1*^*J*_*C-H*_-coupling at the anomeric center of α-*allyl*-manQ **4**, and of the ^*3*^*J*_*C-H*_-HMBC-coupling between its anomeric mannose carbon *C1* and the *C5*-hydrogen atom of its cyclopentene moiety. Both couplings were used to confirm the structure of this compound via NMR, and similar couplings were used to confirm the structures of **3, 10**, and **11**.

With the differently protected cyclopentenes **13, 14** and **15** in hand, the manQ compounds **10, 11**, and **4** were synthesized *via* the glycosylation and reductive amination approach (**Fig. 4c**). The first target compound, β-*allyl*-manQ **10**, was prepared from cyclopentene precursor **13** starting with a stereoselective β-mannosylation reaction (AgOTf, PhSCl, 2,6-di-*tert*-butyl-4-methylpyridine, DCM, - 78 °C, 2 h, 52 %) to give the glycosylated product **17**. Fmoc-deprotection (10 % HNMe_2_ in THF, rt, 1 h) and subsequent reductive amination with deazaguanosine precursor **8** gave the fully protected β-*allyl*-manQ **20**. A three-step final deprotection procedure (1: NaOMe, MeOH, 18 h; 2: 10 % TFA in DCM, 0 °C, 15 min; 3: HF pyridine, MeCN, rt, 18 h) gave β-*allyl*-manQ **10**. For the second target compound, α-*homoallyl*-manQ **11**, we started from cyclopentene precursor **14** and employed *Schmitt-Sinay-* glycosylation conditions with the literature-known glycosyl donor **16** (TMSOTf, THF, 0 °C) to obtain glycosylated product **18**. The synthesis of α-*homoallyl*-manQ **11** was completed from there by Fmoc-deprotection, reductive amination with **8**, and a two-step final deprotection (1: 10 % TFA, DCM, 15 min; 2: NaOMe, MeOH, rt, 2 d).

The third target compound, α-*allyl*-manQ **4**, was synthesized again via *Schmitt-Sinay-*glycosylation conditions with donor **16**. In this case, the SEM-protected cyclopentene precursor **15** was used, because the TBS-protected precursor **13** proved to be sterically too demanding for the glycosylation reaction. Reaction of **15** with **16** however was not fully stereoselective. But after full completion of the synthesis by Fmoc-deprotection, reductive amination, and subsequent three-step deprotection (1: NaOMe, MeOH, rt, 18 h; 2: 10 % TFA, DCM, 0 °C, 15 min; 3: HF · pyridine, MeCN, rt, 1 week), we obtained a mixture of **10** and **4**, which was separated by reversed-phase HPLC-chromatography (see Supplementary Information 3.4). This allowed us to obtain α-*allyl*-manQ **4** in excellent purity.

We next confirmed the structure of the manQ compounds **4, 10**, and **11** using NMR spectroscopy (**Fig. 4d**). For β-*allyl*-manQ **10**, we observed a ^*1*^*J*_*C-H*_-coupling at the mannose anomeric centre of 160.2 Hz, indicative of the β-configuration. The *allylic* connectivity of **10** was confirmed by the ^*3*^*J*_*C-H*_-HMBC-coupling between the anomeric mannose *C1* carbon and the *allylic* hydrogen *C5*H of the cyclopentene ring. For α-*homoallyl*-manQ **11**, a ^*1*^*J*_*C-H*_-coupling of 170.3 Hz was observed at the anomeric centre, proving its α-configuration. In addition, with *δ* = 5.04 ppm, the chemical shift of the anomeric proton *C1*H of **11** was as expected higher than the shift observed for the corresponding β-anomer **3** (*δ* = 4.72 ppm). *Homoallylic* connectivity of **11** was confirmed by observation of an ^*3*^*J*_*C-H*_-HMBC-coupling of the anomeric carbon *C1* and the *homoallylic* hydrogen *C4*H of the cyclopentene moiety. The structure of α-*allyl*-manQ **4** was proven based on an anomeric ^*1*^*J*_*C-H*_ -coupling of 170.0 Hz, indicating its α-configuration. Here, the chemical shift of the anomeric proton *C1*H (*δ* = 4.97 ppm) was again higher than that of the corresponding β-anomer **10** (*δ* = 4.70 ppm), thereby also confirming the α-configuration of **4**. The *allylic* connectivity of **4** was confirmed by an ^*3*^*J*_*C-H*_-HMBC signal resulting from a coupling of the anomeric mannose carbon *C1* with the *allylic* hydrogen *C5*H of the cyclopentene ring.

With all four possible manQ isomers now available (α/β-*allyl-*manQ **4** and **10**, and α/β-*homoallyl*-manQ **3** and **11**), we next performed individual co-injection studies using again a digest of total RNA from mouse liver as reference material, and the UHPLC-MS/MS-method described above for analysis (**Fig. 5**). While for β-*allyl-*manQ **10** and α-*homoallyl*-manQ **11** LC-MS signals were obtained that were well-separated both from galQ **2** and natural manQ (**Fig. 5a**), we discovered to our delight a full signal overlap of our synthetic α-*allyl-*manQ **4** with the natural manQ compound (**Fig. 5b**) We finally confirmed the overlap of the two compound signals with a second HPLC-MS-method using a different HPLC separation column and gradient (see Supplementary Figure 2). Here, too, the synthetic material co-eluted with the natural manQ compound. These co-injection experiments therefore show that the natural manQ compound present in the anticodon loop of tRNA^Asp^ has the chemical structure **4** (see **Fig. 1d** and **4a**). The configuration of its anomeric centre is in fact α, not β. In addition, the mannosyl moiety of natural manQ is connected to the *allylic* hydroxyl group of the cyclopentene unit, and not to the *homoallylic* hydroxyl group as in galQ.

**Figure 5.**
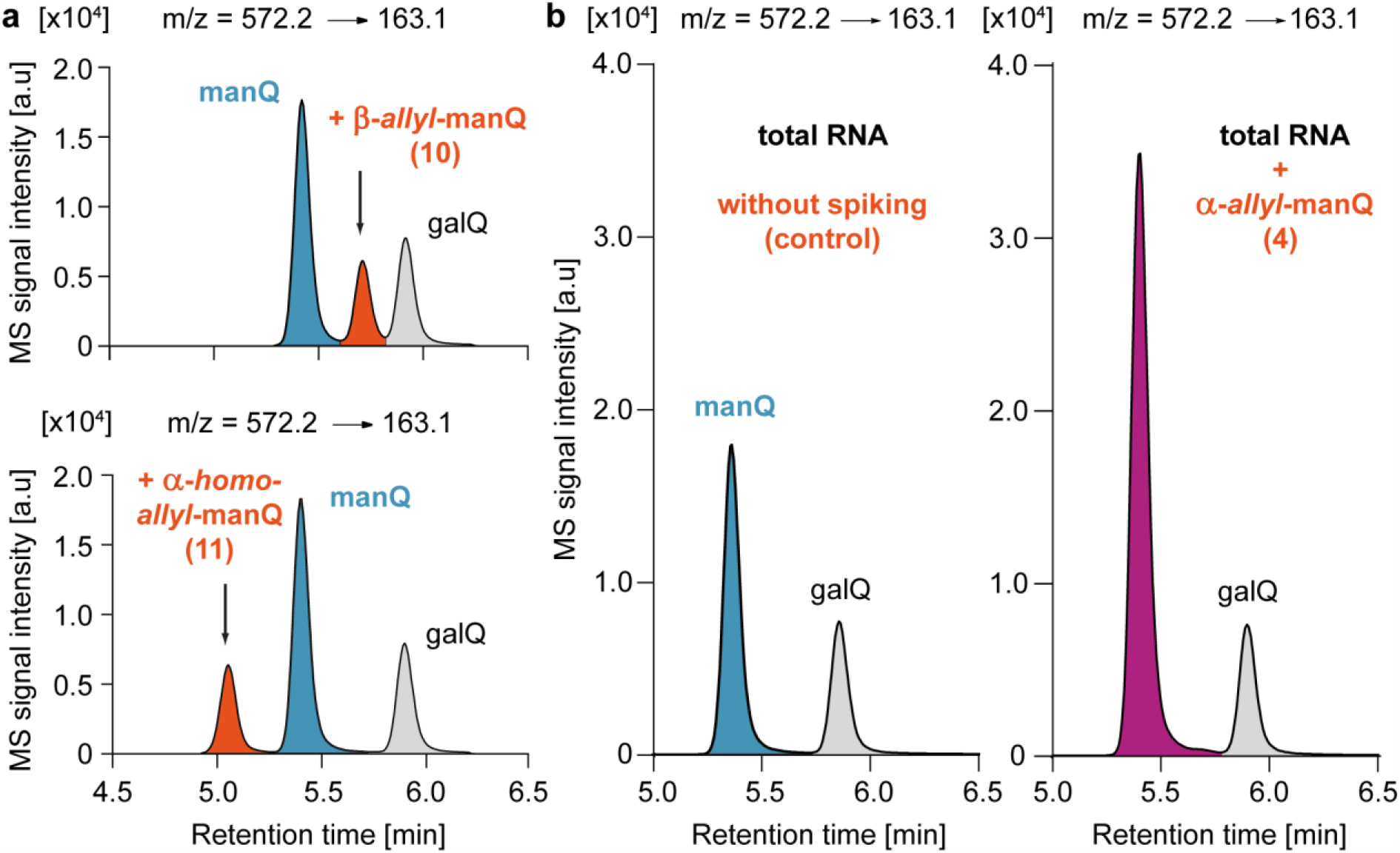
Chromatograms resulting from the co-injection of the synthetic manQ compounds **10, 11**, and **4** with enzymatically digested total RNA from mouse liver analysed by UHPLC-MS/MS. **a**, Co-injection of β-*allyl*-manQ **10** and α-*homoallyl*-manQ **11** show the appearance of a third peak, these compounds are therefore different from natural manQ; **b**, Co-injection of α-*allyl*-manQ **4** with digested total RNA from mouse liver shows a complete signal overlap of **4** and natural manQ, thereby leading to an increased signal intensity of manQ (right) in comparison to the control sample without synthetic standard (left). This result shows that our synthetic compound **4** is identical to the naturally occurring manQ nucleoside and establishes its α-*allyl*-connectivity.

### Conclusion

Based on these experiments, we showed that the widely accepted β-*homoallyl* structure reported in literature for the non-canonical RNA nucleoside manQ is not correct. Instead, natural manQ is an α-*allyl*-mannoside (**4, Fig. 1d**). Its connectivity is therefore maximally different from the β-*homoallyl*-compound galQ (**2, Fig. 1b**). The elucidation of the correct manQ structure and the availability of synthetic material will be the basis for illuminating the biological functions of this hypermodified RNA nucleoside.

## Methods

In order to obtain RNA samples for LC-MS-analysis, mice livers were removed and immediately snap-frozen in liquid nitrogen. The subsequent isolation of total RNA, as well as the isolation of total RNA from HEK cells, were performed using commercially available *TriReagent* from *Sigma Aldrich* as recently described by us. Enzymatic digestion of the total RNA down to the nucleoside level was achieved with the commercially available *Nucleoside Digestion Mix* from *New England Biolabs*. Protocols and additional details for sample preparation can be found in the Supplementary Information. For co-injection experiments, manQ-standards synthesized by us were spiked to digested total RNA samples. Details of this procedure are listed in the Supplementary Information. LC-MS-measurements were performed on a QQQ-UHPLC-MS/MS system and repeated on an Orbitrap LC-MS system. For additional details on the devices and methods used see Supplementary Information. Synthetic procedures for the preparation of the manQ-compounds **3, 4, 10** and **11** as well as characterization data can be found in the Supplementary Information, together with synthetic procedures and characterization data for all compounds newly synthesized in the course of this study.

## Data availability

All data generated and analysed during this study are included in this article and its Supplementary Information. Chromatograms of co-injection experiments are provided in the article figures and Supplementary Information. Synthesis protocols and NMR-spectra for all relevant compounds, proving their chemical identity and bulk purity, are given in the Supplementary Information.

## Acknowledgements

This project has received funding from the European Research Council (ERC) under the European Union’s Horizon 2020 research and innovation program under grant agreement EpiR, No 741912. Further financial support was obtained from the Deutsche Forschungsgemeinschaft (DFG, German Research Foundation) via SFB1309 (Project-ID 325871075) and SPP 1784 (Project-ID 277203618). We thank the Volkswagen Foundation for generous support through the EvoRib VW-project.

## Author contributions

M.H. and P.T. synthesized the compounds. E.K. and M.W. developed UHPLC-MS/MS methods for detection of the manQ derivatives. M.W., E.K., and M.H. performed the LC-MS co-injection-experiments. T.E. performed the metabolic feeding experiments and did the isolation of total RNA from mouse liver and HEK cells. M.W. developed HPLC-MS methods and performed the LC-MS analysis of the feeding experiments. T.C. and M.M. supervised the experiments and the project. T.C. designed the project. T.C. wrote the paper with contributions from M.H., M.W., T.E., E.K., and M.M..

## References

1 Crick, F. Central Dogma of Molecular Biology. Nature 227, 561–563, (1970).

2 Cobb, M. 60 years ago, Francis Crick changed the logic of biology. PLoS Biol. 15, e2003243, (2017).

3 Boccaletto, P. et al. MODOMICS: a database of RNA modification pathways. 2017 update. Nucleic Acids Res. 46, D303–D307, (2018).

4 Cantara, W. A. et al. The RNA modification database, RNAMDB: 2011 update. Nucleic Acids Res. 39, D195–D201, (2011).

5 Sarin, L. P. & Leidel, S. A. Modify or die?--RNA modification defects in metazoans. RNA Biol. 11, 1555–1567, (2014).

6 Rozov, A. et al. Novel base-pairing interactions at the tRNA wobble position crucial for accurate reading of the genetic code. Nat. Commun. 7, 10457, (2016).

7 Agris, P. Wobble position modified nucleosides evolved to select transfer RNA codon recognition: A modified-wobble hypothesis. Biochimie 73, 1345–1349, (1991).

8 Yarian, C. et al. Accurate Translation of the Genetic Code Depends on tRNA Modified Nucleosides. J. Biol. Chem. 277, 16391–16395, (2002).

9 Duechler, M., Leszczyńska, G., Sochacka, E. & Nawrot, B. Nucleoside modifications in the regulation of gene expression: focus on tRNA. Cell. Mol. Life Sci. 73, 3075–3095, (2016).

10 Thumbs, P. et al. Synthesis of galactosyl-queuosine and distribution of hypermodified Q-nucleosides in mouse tissues. Angew. Chem. Int. Ed., (2020).

11 Klepper, F., Jahn, E. M., Hickmann, V. & Carell, T. Synthesis of the transfer-RNA nucleoside queuosine by using a chiral allyl azide intermediate. Angew. Chem. Int. Ed. 46, 2325–2327, (2007).

12 Nishimura, S. Structure, Biosynthesis, and Function of Queuosine in Transfer RNA. Prog. Nucleic Acid Res. Mol. Biol. 28, 49–73, (1983).

13 Harada, F. & Nishimura, S. Possible anticodon sequences of tRNAHis, tRNAAsn, and tRNAAsp from Escherichia coli. Universal presence of nucleoside O in the first position of the anticodons of these transfer ribonucleic acid. Biochemistry 11, 301–308, (1972).

14 Katze, J. R., Basile, B. & McCloskey, J. A. Queuine, a modified base incorporated posttranscriptionally into eukaryotic transfer RNA: wide distribution in nature. Science 216, 55–56, (1982).

15 Salinas-Giegé, T., Giegé, R. & Giegé, P. tRNA biology in mitochondria. Int. J. Mol. Sci. 16, 4518–4559, (2015).

16 Kasai, H. et al. Structure of the modified nucleoside Q isolated from Escherichia coli transfer ribonucleic acid. 7-(4,5-cis-Dihydroxy-1-cyclopenten-3-ylaminomethyl)-7-deazaguanosine. Biochemistry 14, 4198–4208, (1975).

17 Morris, R. C., Brown, K. G. & Elliott, M. S. The effect of queuosine on tRNA structure and function. J. Biomol. Struct. Dyn. 16, 757–774, (1999).

18 Meier, F., Suter, B., Grosjean, H., Keith, G. & Kubli, E. Queuosine modification of the wobble base in tRNAHis influences ‘in vivo’ decoding properties. EMBO J. 4, 823–827, (1985).

19 Tuorto, F. et al. Queuosine-modified tRNAs confer nutritional control of protein translation. EMBO J. 37, e99777, (2018).

20 Zaborske, J. M. et al. A Nutrient-Driven tRNA Modification Alters Translational Fidelity and Genome-wide Protein Coding across an Animal Genus. PLoS Biol. 12, e1002015, (2014).

21 Marks, T. & Farkas, W. R. Effects of a diet deficient in tyrosine and queuine on germfree mice. Biochem. Biophys. Res. Commun. 230, 233–237, (1997).

22 Rakovich, T. et al. Queuosine deficiency in eukaryotes compromises tyrosine production through increased tetrahydrobiopterin oxidation. J. Biol. Chem. 286, 19354–19363, (2011).

23 Farkas, W. R. Effect of diet on the queuosine family of tRNAs of germ-free mice. J. Biol. Chem. 255, 6832–6835, (1980).

24 Fergus, C., Barnes, D., Alqasem, M. & Kelly, V. The Queuine Micronutrient: Charting a Course from Microbe to Man. Nutrients 7, 2897–2929, (2015).

25 Reyniers, J. P., Pleasants, J. R., Wostmann, B. S., Katze, J. R. & Farkas, W. R. Administration of exogenous queuine is essential for the biosynthesis of the queuosine-containing transfer RNAs in the mouse. J. Biol. Chem. 256, 11591–11594, (1981).

26 Tuorto, F. & Lyko, F. Genome recoding by tRNA modifications. Open Biol. 6, (2016).

27 Barbieri, I. & Kouzarides, T. Role of RNA modifications in cancer. Nat. Rev. Cancer 20, 303–322, (2020).

28 Suzuki, T. The expanding world of tRNA modifications and their disease relevance. Nat. Rev. Mol. Cell Biol. 22, 375–392, (2021).

29 Kasai, H. et al. Letter: The structure of Q* nucleoside isolated from rabbit liver transfer ribonucleic acid. J. Am. Chem. Soc. 98, 5044–5046, (1976).

30 Okada, N., Shindo-Okada, N. & Nishimura, S. Isolation of mammalian tRNAAsp and tRNATyr by lectin-Sepharose affinity column chromatography. Nucleic Acids Res. 4, 415–423, (1977).

31 Crich, D. & Sun, S. Direct Formation of β-Mannopyranosides and Other Hindered Glycosides from Thioglycosides. J. Am. Chem. Soc. 120, 435–436, (1998).

32 D.Crich. M. de la Mora, R.C. Synthesis of the mannosyl erythritol lipid MEL A; confirmation of the configuration of the meso-erythritol moiety. Tetrahedron 58, 35–44, (2002).

33 Tvaroska, I. & Taravel, F. R. Carbon-Proton Coupling Constants In The Conformational Analysis of Sugar Molecules. Adv. Carbohydr. Chem. Biochem. 51, 15–61, (1995).

34 Okada, N. & Nishimura, S. Enzymatic synthesis of Q nucleoside containing mannose in the anticodon of tRNA: isolation of a novel mannosyltransferase from a cell-free extract of rat liver. Nucleic Acids Res. 4, 2931–2938, (1977).

