## Supplementary Information for "Synthesis and Structure Elucidation of the Human tRNA Nucleoside Mannosyl-Queuosine"

#### Supporting Information

Markus Hillmeier[a], Mirko Wagner[a], Timm Ensfelder[a], Eva Korytiakova[a], Peter Thumbs[a], Markus Müller[a], and Thomas Carell[a,#]

a) Department Chemie, Ludwig-Maximilians-Universität München, Butenandtstraße 5-13, 81377 München

Correspondence should be addressed to:

Thomas Carell

Telefone: +49 (0)89 218077755

Webpage: www.carellgroup.de

#### Supporting Information - Table of Contents

#### 1. LC-MS-based comparison of natural manQ from mouse liver with our synthetic manQ-compounds for structure elucidation

##### 1.1 Isolation of total RNA from mice livers

The organs of mice were removed and immediately snap-frozen in liquid nitrogen. For subsequent isolation of total RNA, 1 mL *TriReagent (Sigma Aldrich)* was used per 50 mg of organ sample. The tissue was homogenized with a Tissue Lyzer (*Schwingmühle MM400* from *Retsch*), first at 20 Hz for 4 min, then at 30 Hz for 2 min. The homogenized tissue was transferred into a fresh 2 mL-tube and mixed with 200 µl chloroform per 50 mg of organ sample. The phases were separated by centrifugation (12000 g, 15 min, 4 °C). The upper clear phase was transferred to a new 2 mL-tube and mixed with 500 µl isopropanol per 50 mg of organ sample. The RNA was precipitated at -20 °C overnight. The precipitated RNA was pelleted by centrifugation (21130 g, 30 min, 4 °C). The supernatant was carefully removed and 1 mL of ice-cold 75 % ethanol was added, followed by another centrifugation step (21130 g, 20 min, 4 °C). The ethanol washing steps were repeated two more times. The supernatant was removed, and the RNA pellet was first dried at room temperature and then dissolved in nuclease-free water for subsequent enzymatic digestion.

##### 1.2 Enzymatic digestion of total RNA to the nucleoside level

For each LC-MS-measurement, 3 µg of mouse (or HEK 293T) total RNA were digested to the nucleoside level using the *Nucleoside Digestion Mix* (*New England BioLabs*). To this reason, a solution of 3 µg total RNA in 42.5 µL of nuclease-free water was prepared. 5 µL of the *Nucleoside Digestion Mix Reaction Buffer (10x)*, and 2.5 µL of the *Nucleoside Digestion Mix* were added, and the mixture was incubated for 2 h at 37 °C. If needed, the digested total RNA samples could be stored at -20 °C before further treatment.

For the co-injection experiments, samples were subsequently supplemented (spiked) with an appropriate amount of a synthetic manQ compound dissolved in nuclease-free water, or an equal volume of nuclease-free water (control samples). All samples were filtrated before the measurement using an *AcroPrep Advance 96 filter plate 0.2 µm Supor* from *Pall Life Sciences* and subsequently analyzed by LC-MS.

#### 1.3 UHPLC-MS/MS (QQQ)-based co-injection experiments of for manQ structure elucidation

Co-injection experiments were performed by spiking equimolar amounts of standard compared to the natural manQ nucleoside present in the digested total RNA to the sample. Standards were spiked after digestion. UHPLC-MS/MS analysis of digested RNA samples was performed using an *Agilent 1290 UHPL*C system equipped with a UV detector and an *Agilent 6490* triple quadrupole mass spectrometer. The autosampler was cooled to 4 °C. The source-dependent parameters were as follows: gas temperature 230 °C; gas flow 12 L/min (N2); nebulizer 40 psi; sheath gas heater 300 °C; sheath gas flow 6 L/min (N2); capillary voltage 2.500 V in the positive ion mode; capillary voltage −2.250 V in the negative ion mode; nozzle voltage 0 V. The fragmentor voltage was 380 V. Fragmentation was performed with a collision energy of 35 eV and a cell accelerator voltage of 5 V. A specific fragmentation pattern of m/z = 572.2→163.1, as depicted in **Supplementary Fig. 1**, was observed. Delta EMV was set to 500 (positive mode) and 800 (negative mode). Chromatography was performed using a *Poroshell 120 SB-C8 column* (*Agilent*, 2.7 µm, 2.1 mm × 150 mm) at 35 °C with a gradient of water and MeCN, each containing 0.0085 % (v/v) formic acid, at a flow rate of 0.35 mL/min. The gradient was as follows: 0→8min, 0→2 % MeCN (v/v); 8→10.9 min, 2→3.85 % MeCN; 10.9→11.3 min, 3.85 →80 % MeCN; 11.3→12.0 min, 80 % MeCN; 12.0→12.3 min, 80→0 % MeCN; 12.3→14.0 min, 0% MeCN. No other peaks besides the expected ones of manQ, galQ and the spiked standard were detected when using the parameters given above. The identity of galQ was identified by spiking with synthetic galQ-standard (see **Supplementary Fig. 2**).

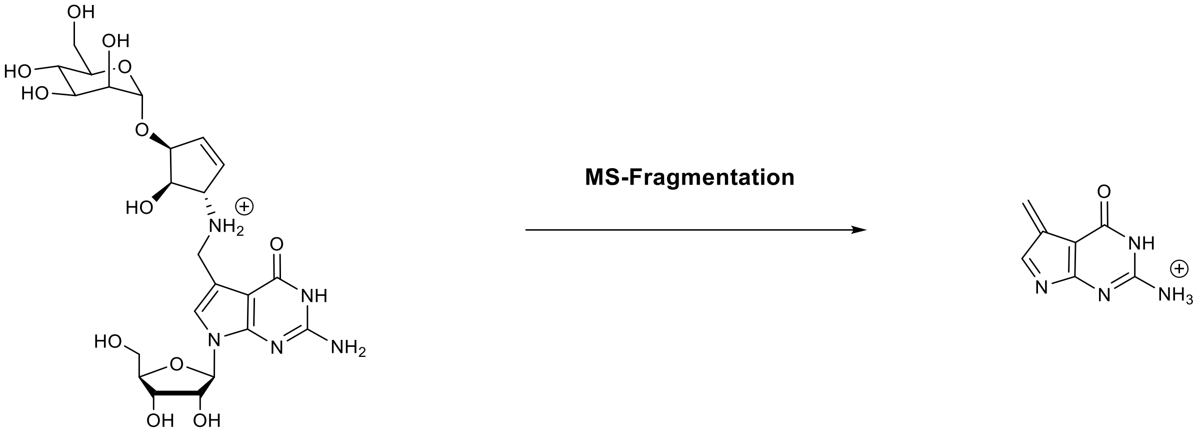

**4**, m/z = 572.2

**23**, m/z = 163.1

**Supplementary Figure 1.** Fragmentation pattern used for specific detection of manQ and galQ on the QQQ mass spectrometer; the fragmentation is depicted for natural manQ **4**, but occurs in a similar way for the precursor ions **2**, **3**, **10,** and **11**, giving the same daughter ion **23** with m/z = 163.1 and the likely structure depicted here.

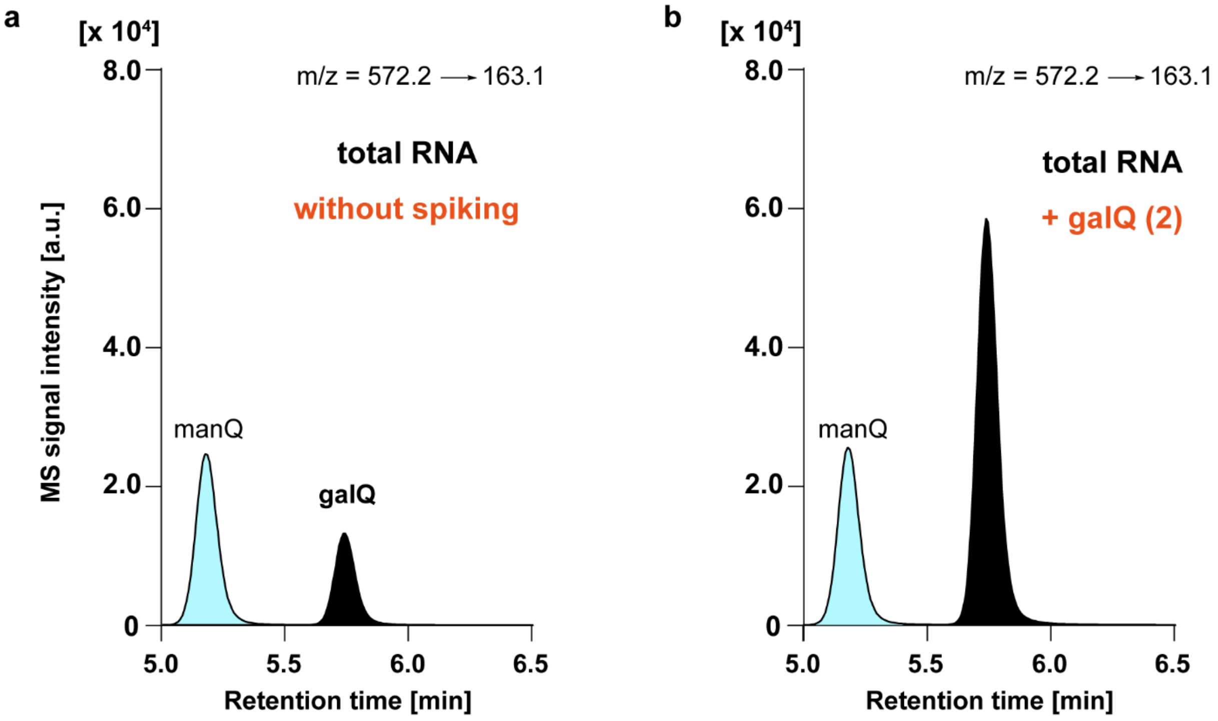

**Supplementary Figure 2.** UHPLC-MS/MS-based identification of the galQ-peak by co-injection; **a:** Digested total RNA from mouse tissue only (control); **b:** Digested total RNA from mouse tissue spiked with synthetic galQ as recently prepared by us[1] shows a complete signal overlap with the peak appearing at higher retention time, therefore identifying this peak as natural galQ.

##### 1.4 HPLC-MS (Orbitrap)-based repetition of co-injection experiments for manQ structure confirmation

To double-check our results from the UHPLC-MS/MS experiments, we repeated the co-injection experiments on a second LC-MS system using a different HPLC separation column. To take into account the lower sensitivity of the Orbitrap system in comparison to the QQQ-system, the samples, containing the digested RNA (see Supplementary Information **1.2**) and the spiked synthetic material, were scaled accordingly. The ratios of digested RNA vs. amount of spiked material, however, were again about equimolar and therefore the same as for the UHPLC-MS/MS analyses. All samples were filtrated before measurement using an *AcroPrep Advance 96 filter plate 0.2 µm Supor* from *Pall Life Sciences*. The HPLC-HESI-MS analyses were performed on a *Dionex Ultimate 3000* HPLC system coupled to a *Thermo Fisher LTQ Orbitrap XL* mass spectrometer. Nucleosides were separated with an *Interchim Uptisphere120-3HDO C18* column whose temperature was maintained at 30 °C. Elution buffers were buffer X (2 mM $NH_4HCOO$ in $H_2O$; pH 5.5) and buffer Y (2 mM $NH_4HCOO$ in $H_2O$/MeCN 20/80 v/v; pH 5.5) and the gradient was as follows: 0→10 min, 0 % Y, 0.15 mL/min; 10→50 min, 0→5 % Y,

0.15 mL/min; 50→52 min, 5→75 % Y, 0.15 mL/min→0.20 mL/min; 52→55 min, 75 % Y, 0.20 mL/min. The chromatogram was recorded at 260 nm with a *Dionex Ultimate 3000 Diode Array Detector,* and the chromatographic eluent was directly injected into the ion source of the mass spectrometer without prior splitting. Ions were scanned in the positive polarity mode over a full-scan range of m/z = 225-2000 with a resolution of 100,000. Parameters of the mass spectrometer were tuned with a freshly mixed solution of inosine (5 µM) in buffer X and set as follows: Capillary temperature 275 °C; APCI vaporizer temperature 100 °C; sheath gas flow 5.00; aux gas flow 21.0; sweep gas flow 1.00; source voltage 4.80 kV; capillary voltage 0 V; tube lens voltage 45.0 V; skimmer offset 0 V. The ion chromatograms of the compounds of interest were extracted from the total ion current (TIC) chromatogram with a mass range set to +/- 0.0020 u around the exact mass $[M+H]^+$ of the compounds of interest.

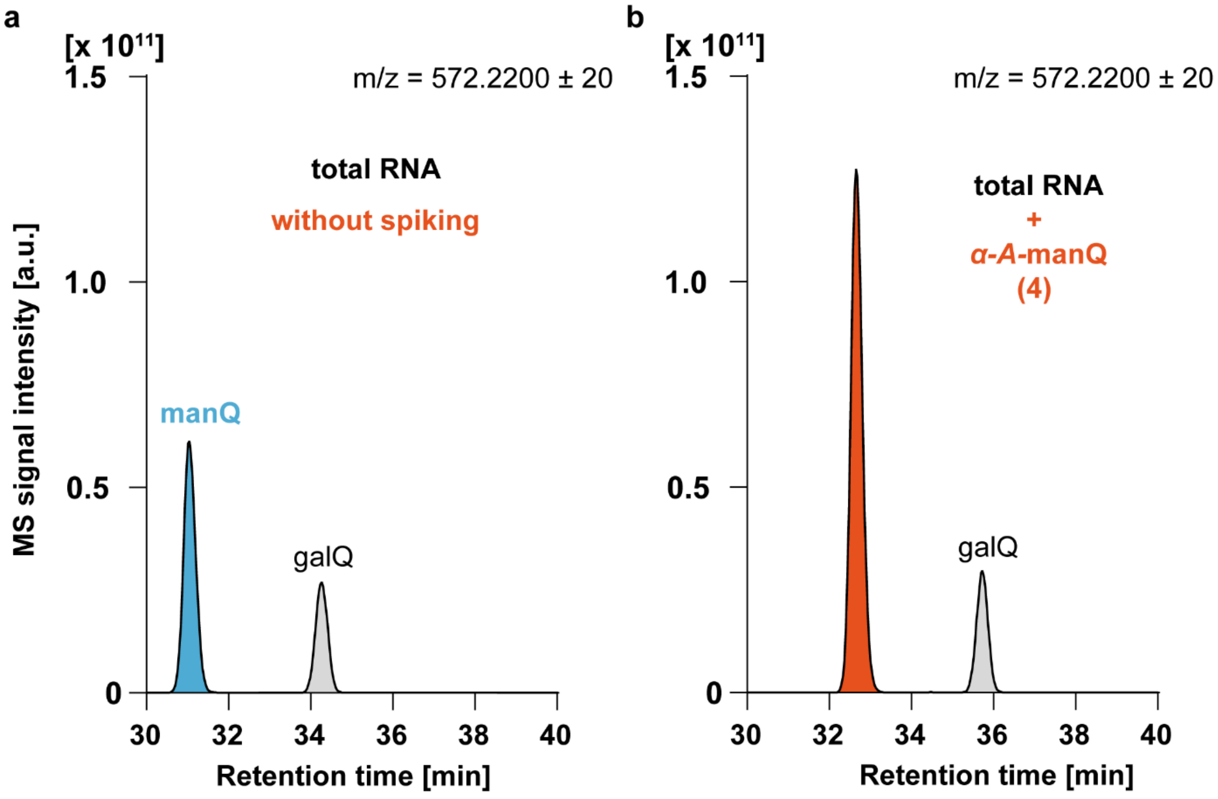

**Supplementary Figure 3.** Extracted ion chromatograms resulting from the co-injection of the synthetic manQ compound **4** with enzymatically digested total RNA from mouse liver analyzed by HPLC-MS: Co-injection of α-*allyl*-manQ **4** with digested total RNA from mouse liver shows a complete signal overlap of **4** and natural manQ, thereby leading to an increased signal intensity of manQ (**b**) in comparison to the control sample (**a**) without synthetic standard. This result confirms the findings from the UHPLC-MS/MS-experiments and shows again that our synthetic α-*allyl*-manQ **4** is identical to the naturally occurring manQ compound.

As shown in **Supplementary Fig. 3**, these experiments again did NOT result in the appearance of a new peak, but resulted in a full signal overlap of the natural manQ compound from mouse liver and our synthetic $\alpha$-*allyl*-manQ **4**. As expected from our UHPLC-MS/MS co-injection experiments (see Supplementary Information **1.3**), these results thereby confirm the identity of the natural manQ compound as $\alpha$-*allyl*-manQ **4**.

Of note, the proper HPLC-separation of the manQ-compounds on this system was more sensitive towards changes of gradient and column age than in case of the UHPLC-based separation with a *Poroshell 120 SB-C8* column. In rare cases, when the column had been used for a prolonged period of time, the peaks of synthetic $\alpha$-*homoallyl*-manQ **11** and of the natural manQ compound $\alpha$-*allyl*-manQ **4** were not properly separable any more.

#### 2. Metabolic feeding study to confirm mannose as the hexose-part of natural manQ

##### 2.1 Cell culture metabolic feeding

To determine which hexose is incorporated into the mannosyl-queuosine of $tRNA^{Asp}$, HEK 293T cells were cultured in different heavy RPMI media (see **Supplementary Table *1***). The heavy RPMI media contained two monosaccharides each: One of the two hexoses was fully $^{13}C$-isotope-labelled, while the other hexose was non-labelled and used in these experiments as a suppressor of metabolic interconversion. Heavy RPMI media with the following hexose combinations were used:

- D-mannose-$^{13}C_6$, D-glucose
- D-mannose, D-glucose-$^{13}C_6$
- D-galactose-$^{13}C_6$, D-Glucose
- D-galactose, D-glucose-$^{13}C_6$

1.3 million HEK 293T cells were seeded on a p60 cell culture dish. Cultivation was performed for 18 - 21 h in RPMI medium at 37 °C and 5 % $CO_2$. Subsequently, the medium was removed and replaced with heavy RPMI medium. Incubation continued under the under the same conditions for 30, 60 and 180 min.

**Supplementary Table 1:** Composition of the used media. Light RPMI Medium corresponds to regular RPMI medium. For Heavy RPMI medium, a glucose free RPMI mixture was used (w/o g.: without glucose). A combination of either mannose and glucose or galactose and glucose were added to 11 mM final concentration each. One of them was fully $^{13}C$-labelled in each mixture.

| Medium | Composition |
|---|---|
| Light RPMI medium | 88 % (v/v) RPMI; 10 % (v/v) FBS; 1 % (v/v) Pen-Strep; 1 % Ala-Gln |
| Heavy RPMI medium gal/glc | 88 % (v/v) RPMI (w/o g.); 10 % (v/v) FBS; 1 % (v/v) Pen-Strep; 1 % Ala-Gln; 11 mM D-galactose(-$^{13}C_6$); 11 mM D-glucose(-$^{13}C_6$) |
| Heavy RPMI medium man/glc | 88 % (v/v) RPMI (w/o g.); 10 % (v/v) FBS; 1 % (v/v) Pen-Strep; 1 % Ala-Gln; 11 mM D-mannose(-$^{13}C_6$); 11 mM D-glucose(-$^{13}C_6$) |

##### 2.2. Isolation of total RNA from HEK 293T cells

After completion of the respective time, the medium was removed and the cells were carefully detached from the cell culture dish using PBS and transferred to a 2 ml reaction tube. This was followed by pelleting for 1 min at 500 g and 4 °C. The PBS was removed and the cell pellet immediately resuspended in 1 ml *TriReagent (Sigma Aldrich)*. Samples taken after 30 min in labelled medium were incubated for 40 min, the 60 min- and 180 min-samples were incubated for 5 min. Following incubation, 200 µL of chloroform were added to the sample, which then was heavily vortexed and subsequently centrifuged for 15 min at 12,000 g and 4 °C. The upper clear phase was transferred to a new 2 ml reaction tube. Isopropanol was added to the transferred phase and mixed. Precipitation of the RNA was performed at -20 °C overnight. Following this, samples were directly pelleted for 30 min at 21,130 g and 4 °C. The supernatant was carefully removed. Thereafter, 1 ml of 75 % (v/v) cold ethanol (-20 °C) was added and again the sample was centrifuged for 20 min at 21,130 g and 4 °C. The ethanol washing step including the centrifugation was repeated two more times. The supernatant was removed and the pellet was dried at room temperature. Thereafter, the pellet was dissolved in nuclease-free water.

##### 2.2 LC-MS-analysis of total RNA from metabolically labeled HEK cells

HPLC-HESI-MS analysis of the enzymatically digested total RNA of HEK cells (see **Supplementary Information 1.2**) were performed on a *Dionex Ultimate 3000* HPLC system coupled to a *Thermo Fisher LTQ Orbitrap XL* mass spectrometer. Samples were filtrated before the measurement using an *AcroPrep Advance 96 filter plate 0.2 µm Supor* from *Pall Life Sciences*. Nucleosides were separated with an *Interchim Uptisphere120-3HDO C18* column whose temperature was maintained at 30 °C. Elution buffers were buffer X (2 mM $NH_4HCOO$ in $H_2O$; pH 5.5) and buffer Y (2 mM $NH_4HCOO$ in $H_2O$/MeCN 20/80 v/v; pH 5.5) with a flow rate of 0.15 mL/min. The gradient was as follows: 0→3.5 min, 0 % Y; 3.5→4 min, 0→0.2 % Y; 4→10 min, 0.2 % Y; 10→50 min, 0.2→4.7% Y; 50→55 min, 4.7→60 % Y; 55→57 min, 60→100 % Y; 57→62 min, 100 % Y. The chromatogram was recorded at 260 nm with a *Dionex Ultimate 3000 Diode Array Detector,* and the chromatographic eluent was directly injected into the ion source of the mass spectrometer without prior splitting. Ions were scanned in the positive polarity mode over a full-scan range of m/z = 210-800 with a resolution of 60,000. Parameters of the mass spectrometer were tuned with a freshly mixed solution of inosine (5 µM) in buffer X and set as follows: Capillary temperature 275 °C; APCI vaporizer temperature 100 °C; sheath gas flow 5.00; aux gas flow 21.0; sweep gas flow 1.00; source voltage 4.80 kV; capillary voltage 0 V; tube lens voltage 45.0 V; skimmer offset 0 V. The ion chromatograms of the compounds of interest were extracted from the total ion current (TIC) chromatogram with a mass range set to +/- 0.0050 u around the exact mass $[M+H]^+$ of a compound. The peak areas in the extracted ion chromatograms of the heavy and corresponding light compound were integrated and the percentual $^{13}C_6$-labelling of a compound within a sample was calculated.

### 3. Chemical Synthesis

#### 3.1 Synthesis of β-homoallyl-manQ 3

##### 3.1.1 Synthesis of glycosyl donor 5

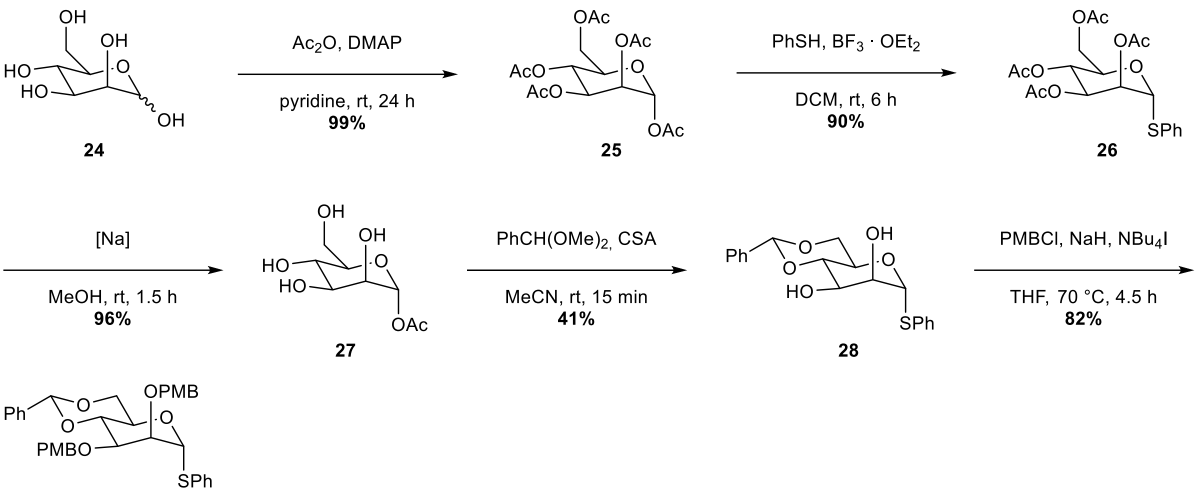

5

**Supplemenary Figure 4.**Synthesis of glycosyl donor **5** in 5 steps starting from D-mannose (29% overall yield)

**1,2,3,4,6-Penta-*O*-acetyl-α-D-mannopyranose 25**[2]

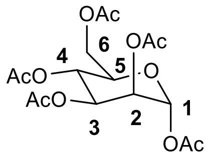

D-Mannose (5.00 g, 27.8 mmol, 1.00 eq) was dissolved in pyridine (104 mL). $Ac_2O$ (78.0 mL, 416 mmol, 30.0 eq) and DMAP (339 mg, 2.78 mmol, 0.10 eq) were added at 0 °C. The reaction was stirred at room temperature for 24 h, which resulted in an orange solution. After evaporation of the solvent *in vacuo* the residue was taken up in EtOAc (100 mL) and washed with aqueous HCl (1 M, 100 mL), $H_2O$ (2 x 100 mL) and brine (100 mL). After evaporation of the solvent *in vacuo* the product (10.8 g, 27.7 mmol, 99%) was obtained as yellow viscous syrup. (only α-anomer).

**$R_f$** (EtOAc/*iso*-hexane 1:1) = 0.50

**$^1$H-NMR** ($CDCl_3$, 400 MHz): δ = 6.08 (d, *J* = 1.9 Hz, 1H, C1H), 5.35 (m, 2H, C3H, C4H), 5.26 (td, $J_1$ = 2.2 Hz, $J_2$ = 0.7 Hz, 1H, C2H ), 4.28 (dd, $J_1$ = 12.4, $J_2$ = 4.9 Hz, 1H, C6Ha), 4.13-4.02 (m, 2H, C5H, C6Hb 2.18 (s, 3 H, $CO_2CH_3$), 2.17 (s, 3 H, $CO_2CH_3$), 2.09 (s, 3 H, $CO_2CH_3$), 2.05 (s, 3 H, $CO_2CH_3$), 2.01 (s, 3 H, $CO_2CH_3$) ppm.

**$^{13}$C-NMR** ($CDCl_3$, 101 MHz): δ = 170.8 ($CO_2CH_3$), 170.2 ($CO_2CH_3$), 169.9 ($CO_2CH_3$), 169.7 ($CO_2CH_3$), 168.2 ($CO_2CH_3$), 90.7 (C1), 70.7 (C5), 68.8 (C3), 68.4 (C2), 65.6 (C4), 62.2 (C6), 21.0 ($CO_2CH_3$), 20.9 ($CO_2CH_3$), 20.9 ($CO_2CH_3$), 20.8 ($CO_2CH_3$), 20.8 ($CO_2CH_3$) ppm.

**HRMS** (ESI$^+$) m/z: calc. for $C_{16}H_{22}O_{11}Na$ $[M + Na]^+$: 413.1060; found: 413.1056.

NMR-spectra in accordance with literature.

##### 2,3,4,6-Penta-*O*-acetyl-1-thiophenyl-α-D-mannopyranoside 26[3]

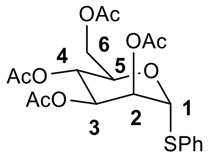

**25** (5.00 g, 12.8 mmol, 1.00 eq) was dissolved in DCM (25 mL). Thiophenol (2.65 mL, 2.82 g, 25.6 mmol, 2.00 eq) and $BF_3 \cdot OEt_2$ (4.80 mL, 5.45 g 34.8 mmol, 3.00 eq) were added at 0 °C. The reaction was stirred at room temperature for 6 h. Saturated aqueous $NaHCO_3$-solution (20 mL) was added and the reaction mixture was stirred for 15 min (until no more gas evolution was observable). The phases were separated and the aqueous phase was re-extracted with DCM (2 x 30 mL). The combined organic phases were washed with brine (20 mL), dried over $Na_2SO_4$, filtered and the solvent was removed *in vacuo*. Column chromatographic purification (EtOAc/*iso*-hexane 1:3) afforded the product (5.09 g, 11.6 mmol, 90%) as yellow syrup.

**$R_f$** (EtOAc/*iso*-hexane 1:1) = 0.76

**$^1$H-NMR** ($CDCl_3$, 600 MHz): δ = 7.44 (m, 2H, arom. CH), 7.26 (m with solvent peak, 3H, arom. CH), 5.46 (m, 2H, C1H, C2H), 5.28 (m, 2H, C3H, C4H), 4.50 (m, 1H, C5H), 4.26 (dd, $J_1$ = 12.2, $J_2$ = 5.9 Hz, 1H, C6Ha), 4.07 (m, 1H, C6Hb), 2.11 (s, 3H, $CO_2CH_3$), 2.03 (s, 3H, $CO_2CH_3$), 2.01 (s, 3H, $CO_2CH_3$), 1.97 (s, 3H, $CO_2CH_3$) ppm.

**$^{13}$C-NMR** ($CDCl_3$, 125 MHz): δ = 170.7 ($CO_2CH_3$), 170.0 ($CO_2CH_3$), 169.9 ($CO_2CH_3$), 169.9 ($CO_2CH_3$), 132.7 (arom. C), 132.2 (arom. C), 129.3 (arom. C), 128.2 (arom. C), 85.5 (C1), 71.0 (C2), 69.6 (C5), 69.5 (C3/C4), 66.5 (C3/C4), 62.6 (C6), 21.0 ($CO_2CH_3$), 20.8 ($CO_2CH_3$), 20.8 ($CO_2CH_3$), 20.8 ($CO_2CH_3$) ppm.

**HRMS** ($ESI^+$) calc. for $C_{20}H_{24}NaO_9S$: 463.1039 $[M + Na]^+$; found: 463.1037;

Analytical data in accordance with literature.

**1-Thiophenyl-α-D-mannopyranoside 27**[3]

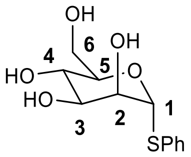

**26** (5.09 g, 11.6 mmol) was dissolved in MeOH (50 mL). A catalytic amount of sodium was added and the reaction mixture was stirred for 1.5 h at room temperature resulting in a suspension. The reaction was neutralized with AcOH, upon which the precipitate redissolved. Evaporation of the solvent gave the product (2.69 g, 11.1 mmol, 96%) as colorless foam.

**$R_f$** (EtOAc/*iso*-hexane 1:1) = 0.03

**$^1$H-NMR** (MeOD, 600 MHz): δ = 7.55-7.52 (m, 2H, aromatic CH), 7.34-7.26 (m, 3H, aromatic CH), 5.43 (d, $J$ = 1.6 Hz, 1H, C1H), 4.09 (dd, $J_1$ = 3.2 Hz, $J_2$ = 1.6 Hz, 1H, C2H), 4.04 (ddd, $J$ = 9.4, 5.4, 2.5 Hz, 1H, C5H), 3.83 (dd, $J_1$ = 12.0, $J_2$ = 2.5 Hz, 1H, C6Ha), 3.77 (dd, $J$ = 12.0, 5.5 Hz, 1H, C6Hb), 3.73 (t, $J_1$ = $J_2$ = 9.4 Hz, 1H, C4H), 3.69 (dd, $J_1$ = 9.4 Hz, $J_2$ = 3.1 Hz, 1H, C3H)

**$^{13}$C-NMR** (MeOD, 125 MHz): δ = 135.9 (aromatic C), 133.0 (aromatic C), 130.0 (aromatic C), 128.5 (aromatic C), 90.5 (C1), 75.7 (C5), 73.8 (C2), 73.2 (C3), 68.7 (C4), 62.6 (C6) ppm.

**HRMS** (ESI$^-$) calc. for $C_{12}H_{15}O_5S$: 271.0646 [M-H]$^-$; found: 271.0644

Analytical data in accordance with literature.

**4,6-O-Benzylidene-1-thiophenyl-α-D-mannopyranoside 28**[4]

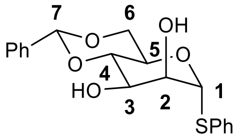

**27** (1.00 g, 3.67 mmol, 1.00 eq) was dissolved in MeCN (15 mL). CSA (213 mg, 0.918 mmol, 0.25 eq) and $PhCH(OMe)_2$ (0.61 mL, 615 mg, 4.04 mmol, 1.10 eq) were added. The reaction was stirred at room temperature for 15 min, after which a solid precipitated. DCM (50 mL) and $H_2O$ (50 mL) were added and the phases were separated. The aqueous phase was reextracted with DCM (2 x 30 mL) and the combined organic phases were dried over $Na_2SO_4$, filtered and the solvent was evaporated. Column chromatographic purification (MeOH/DCM 1:200) gave the product (540 mg, 1.50 mm0l, 41%) as colorless solid.

**$R_f$** (MeOH/DCM 1:100) = 0.24

**$^1$H-NMR** (MeOD, 600 MHz): δ = 7.53 (m, 4H, arom. CH), 7.34 (m, 6H, arom. CH), 5.64 (s, 1H, C7H), 5.50 (d, $J$ = 1.2 Hz, 1H, C1H), 4.25 (td, , $J_1$ = 9.7 Hz, $J_2$ = 4.9 Hz, 1H, C5H), 4.19 (dd, $J_1$ = 3.3 Hz, $J_2$ = 1.4 Hz, 1H, C2H), 4.14 (dd, , $J_1$ = 10.2 Hz, $J_2$ = 4.9 Hz, 1H, C6Ha), 4.04 (t, $J$ = 9.5 Hz, 1H, C4H), 3.96 (dd, $J_1$ = 9.9 Hz, $J_2$ = 3.3 Hz, 1H, C3Hw), 3.85 (t, $J$ = 10.4 Hz, 1H, C6Hb) ppm.

**$^{13}$C-NMR** (MeOD, 600 MHz): δ = 137.4, 132.9, 130.2, 130.1, 129.1, 128.7, 127.5, 125.8 (all aromatic C), 103.4 (C7), 89.4 (C1), 72.9 (C2), 72.8 (C4), 68.6 (C3), 68.2 (C6), 65.1 (C5) ppm.

**HRMS** (ESI$^+$) calc. for $C_{19}H_{21}O_5S$: 361.1104 $[M + H]^+$ ; found: 361.1106.

Analytical data in accordance with literature.

**4,6-O-Benzylidene-2,3-di-*O*-(*para*-methoxybenzyl)-1-thiophenyl-α-D-mannopyranoside 5**[5]

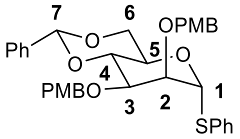

**28** (540 mg, 1.50 mmol, 1.00 eq) was dissolved in THF (11 mL). NaH (240 mg, 6.00 mmol, 4.00 eq) was added at 0 °C. Next, PMB-Cl (0.81 mL, 940 mg, 6.00 mmol, 4.00 eq) and $NBu_4I$ (28.0 mg, 0.075 mmol, 0.05 eq) were added. The reaction was stirred at 70 °C for 4.5 h before being cooled to room temperature. Saturated aqueous $NaHCO_3$-solution (20 mL) was added and the phases were separated. The aqueous phase was re-extracted with EtOAc (3 x 50 mL). The combined organic phases were dried over $Na_2SO_4$, filtered and the solvent was removed *in vacuo*. Column chromatographic purification (EtOAc/*iso*-hexane 1:2) afforded the product (739 mg, 1.23 mmol, 82%) as colorless syrup.

$\mathbf{R_f}$ (MeOH/DCM 1:100) = 0.24

**$^1$H-NMR** ($CDCl_3$, 400 MHz): δ = 7.33 (m, 14H, arom. CH), 6.85 (m, 4H, arom. CH), 5.62 (s, 1H, C7H), 5.43 (d, *J* = 1.4 Hz, 1H, C1H), 5.62 (s, 1H, C1H), 5.43 (d, *J* = 1.4 Hz, 1H), 4.72 (d, *J* = 11.8 Hz, 1H, $CH_2$), 4.63 (s, 2H, $CH_2$), 4.56 (d, *J* = 11.8 Hz, 1H, $CH_2$), 4.23 (m, 3H, C3H, C5H, C6Ha), 3.97 (dd, $J_1$ = 3.3 Hz, $J_2$ = 1.4 Hz, 1H, C2H), 3.89 (m, 2H, C4H, C6Hb), 3.80 ($OCH_3$), 3.79 ($OCH_3$) ppm.

**$^{13}$C-NMR** ($CDCl_3$, 100 MHz): δ = 159.4, 159.3, 137.7, 134.0, 131.7, 130.6, 130.0, 129.9, 129.5, 129.2, 129.0, 128.3, 127.7, 126.2, 114.0, 113.9 (all aromatic C), 101.6 (C7), 87.3 (C1), 79.1 (C3), 77.6 (C2), 75.9 (C4), 72.8 ($CH_2$), 72.8 ($CH_2$), 68.6 (C6), 65.6 (C5), 55.4 (2C, $OCH_3$) ppm.

**HRMS** ($ESI^+$) calc. for $C_{35}H_{36}O_7SNa$: 623.2079 $[M + Na]^+$; found: 623.2076.

Analytical data in accordance with literature.

**3.1.2 Synthesis of ß-HA-ManQ**

**(9H-fluoren-9-yl)methyl((1S,4S,5R)-5-(((2R,4aR,6R,7S,8S,8aR)-7,8-bis((4-methoxy-benzyl)oxy)-2-phenylhexahydro-pyrano[3,2-d][1,3]dioxin-6-yl)oxy)-4-((tert-butyl-dimethylsilyl)oxy)cyclopent-2-en-1-yl)carbamate 7**

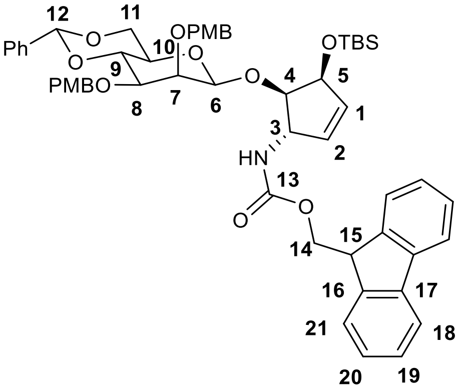

Cyclopentene **6** (100 mg, 219 µmol, 1.00 eq) was dissolved in DCM (2.5 mL) and stirred with 3 Å molsieves for 15 min (solution **A**). Glycosyl donor **5** (131 mg, 219 µmol, 1.00 eq) and DTBMP (135 mg, 656 µmol, 3.00 eq) were dissolved in DCM (2.5 mL) and stirred with 3 Å molsieves for 15 min (solution **B**). AgOTf (168 mg, 656 µmol, 3.00 eq) was dissolved in DCM (2.5 mL) and 3 Å molsieves were added. The reaction was cooled to -78 °C and stirred at that temperature for 15 min. A solution of PhSCl (546 µmol, 2.50 eq) in DCM (2.7 mL) was added and the reaction mixture was stirred at -78 °C for 5 min. Next, solution **B** was added and the reaction was stirred at -78 °C for 10 min. Then solution **A** was added and the reaction was stirred at -78 °C for 2 h. Saturated aqueous $NaHCO_3$-solution (20 mL) was added and the mixture was warmed to room temperature. After filtration, the phases were separated and the aqueous phase was re-extracted with DCM (3 x 20 mL). The combined organic phases were dried over $Na_2SO_4$, filtered and the solvent was removed *in vacuo*. Column chromatographic purification (EtOAc/*iso*-hexane 1:5 → 1:3) gave the product (136 mg, 142 µmol, 62%) as colorless solid.

**$R_f$** (EtOAc/*iso*-hexane 1:3) = 0.12.

**$^1$H-NMR** (600 MHz, $CDCl_3$): δ = 7.75 (d, $J$ = 7.7 Hz, 2H, C21H), 7.56 (d, $J$ = 7.4 Hz, 1H, C18H), 7.49 (m, 2H, C120H), 7.36 (m, 7H, arom. CH), 7.29 (m, 2H, C19H), 7.14 (d, $J$ = 8.2 Hz, 2H, arom. CH), 6.83 (m, 2H, arom. CH), 6.79 (d, $J$ = 8.4 Hz, 2H, arom. CH), 5.93 (m, 1H, C1H), 5.82 (d, $J$ = 6.2 Hz, 1H, C2H), 5.59 (s, 1H, C12H), 4.83 (m, 4H, C3H, C6H (**$^1J_{C\text{-}H}$ = 158 Hz**), $CH_2$), 4.51 (m, 6H, C13H, C5H, 2 x $CH_2$), 4.25 (dd, , $J_1$ = 10.3 Hz, $J_2$ = 4.8 Hz, 1H, C11Ha), 4.20 (t, , $J$ = 6.7 Hz, 1H, C14H), 4.11 (m, 1H, C9H), 3.96 (m, 2H, C4H, C7H), 3.86 (t, $J$ = 10.3 Hz, 1H, C11Hb), 3.78 (s, 3H, $OCH_3$), 3.77 (s, 3H, $OCH_3$), 3.53 (d, , $J$ = 9.8 Hz, 1H, C8H), 3.24 (m, 1H, C10H), 0.87 (s, 9H, $SiC(CH_3)_3$), 0.07 (s, 6H, $SiCH_3$) ppm.

**$^{13}$C-NMR** (150 MHz, $CDCl_3$): δ = 159.2 (arom. C), 155.8 (C13), 143.9 (C17), 141.5 (C16), 134.6 (C2), 134.3 (C1), 130.41 (arom. C), 129.2 (arom. C), 128.9 (arom. C), 128.3 (arom. C), 127.9 (arom. C), 127.2 (C19), 126.2 (C20), 125.0 (C18), 120.1 (C21), 113.8 (arom. C), 113.6 (arom. C), 101.5 (C12), 101.1 (C6), 82.8 (C4), 78.7 (C9), 77.6 (C8), 75.3 (C7), 74.4 ($OCH_2$), 74.3 (C5), 71.9 ($OCH_2$), 68.8 (C11), 67.8 (C10), 66.7 (C14), 59.0 (C3), 55.4 ($OCH_3$), 55.4 ($OCH_3$), 47.4 (C15), 26.1 ($SiC(CH_3)_3$), -4.2 ($SiCH_3$), -4.4 ($SiCH_3$) ppm.

**HRMS** ($ESI^+$) calc. for $C_{55}H_{67}N_2O_{11}Si$: 959.4509 $[M + NH_4]^+$; found: 959.4524.

**β-homoallyl-Mannosyl-Queuosine 3**

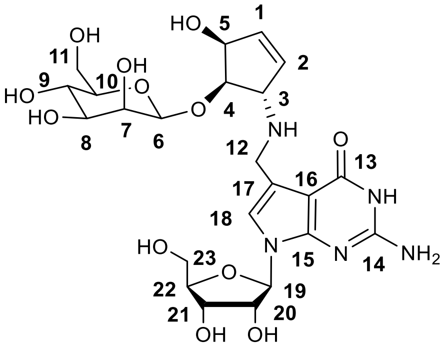

**12** (23 mg, 24.0 µmol was dissolved in 10% $HNMe_2$ in THF (1 mL). The reaction was stirred for 1 h at room temperature before the solvent was removed *in vacuo*. The resulting yellow solid was washed with hexane to afford the product as colorless solid. The resulting Fmoc-deprotected derivative (14.9 mg, 20.6 µmol, 1.00 eq) was dissolved in MeOH (0.5 mL) and **13** (14.6 mg, 20.6 µmol, 1.00 eq) and AcOH (1 µL) were added. The reaction was stirred at rt for 5 h before it was cooled to 0 °C and $NaBH_4$ (2.02 mg, 53.6 µmol, 2.60 eq) was added. The reaction was stirred at 0 °C for 1 h. Then $H_2O$ was added and the solvent was evaporated. The residue was dissolved in MeOH (1 mL) and NaOMe (27.0 mg, 0.5 mmol) was added. The reaction mixture was stirred at room temperature for 5 h, after which LC-MS indicated deprotection of all ester type protecting groups. The solution was neutralized with DOWEX-$H^+$-resin, then filtered and the solvent was evaporated. The residue was taken up in DCM (0.9 mL) and TFA (0.1 mL) was added at 0 °C. The reaction was stirred at 0 °C for 20 min, then saturated aqueous $NaHCO_3$-solution was added (1 mL). The mixture was evaporated and the residue was taken up in pyridine (1.5 mL) and HF · pyridine (7.5 eq) was added in a plastic falcon. The reaction was stirred for 18 h at room temperature before TMSOMe (15.0 eq) was added at 0 °C. After stirring the reaction for 1 h at 0 °C, the solvent was evaporated *in vacuo*. The residue was dissolved in $H_2O$ (5 mL), filtered and subjected to preparative HPLC-purification (0-8% buffer B, 45 min, $R_t$ = 29.2 min) to afford β-homoallyl-ManQ as a colorless solid (2.99 mg, 5.24 µmol, 22% over 5 steps).

**RP-HPLC:** $R_t$ (C18-column, 0-8% buffer B, 45 min) = 29.2 min.

**$^1$H-NMR** (MeOD, 600 MHz): δ = 7.04 (s, 1H, C18H), 6.11 (ddd, $J_1$ = 6.3 Hz, $J_2$ = 2.6 Hz, , $J_3$ = 1.9 Hz, 1H, C1H), 6.07 (dd, , $J_1$ = 6.3 Hz, , $J_2$ = 1.9 Hz, 1H, C2H), 5.93 (d, $J$ = 5.9 Hz, 1H, C19H), 4.72 (d, $J$ = 0.7 Hz, $^{\mathbf{1}}\boldsymbol{J}_{\mathbf{C\text{-}H}}$ **= 160 Hz**, 1H, C6H), 4.68 (ddd, , $J_1$ = 5.1 Hz, , $J_2$ = 2.6 Hz, , $J_3$ = 1.5 Hz, 1H, C5H), 4.43 (t, $J$ = 5.5 Hz, 1H, C20H), 4.23 (dd, , $J_1$ = 5.4 Hz, $J_2$ = 3.6 Hz, 1H, C21H), 4.16 (m, 4H, C3H, C4H, C12H), 4.02 (q, , $J$ = 3.4 Hz, 1H, C22H), 3.98 (dd, , $J_1$ = 3.2 Hz, $J_2$ = 0.4 Hz, 1H, C7H), 3.87 (dd, , $J_1$ = 11.8 Hz, $J_2$ = 2.3 Hz, 1H, C11Ha), 3.81 (dd, , $J_1$ = 12.2 Hz, $J_2$ = 3.1 Hz, 1H, C23Ha), 3.75 (dd, $J_1$ = 12.0 Hz, , $J_2$ = 5.6 Hz, C11Hb), 3.72 (dd, $J_1$ = 12.2 Hz, , $J_2$ = 3.6 Hz, 1H, C23Hb), 3.62 (t, $J$ = 9.6 Hz, 1H, C9H), 3.49 (dd, $J_1$ = 9.4 Hz, $J_2$ = 3.2 Hz, 1H, C8H), 3.28 (ddd, $J_1$ = 9.8 Hz, $J_2$ = 5.5 Hz, $J_3$ = 2.3 Hz, C10H), 1.93 (s, 3H, $OAc^-$) ppm.

**$^{13}$C-NMR** (MeOD, 150 MHz): δ = 162.3 (C12), 154.4 (C14), 153.6 (C15), 136.3 (C1), 132.9 (C2), 130.9 (arom. C), 130.6 (arom C), 119.4 (C18), 113.9 (C17), 102.5 (C6), 100.8 (C16), 89.7 (C19), 86.5 (C22), 84.9 (C4), 78.5 (C10), 75.7 (C20), 75.1 (C8), 75.0 (C5), 72.3 (C7), 72.2 (C21), 68.2 (C9), 65.9 (C3), 63.3 (C23), 62.6 (C11), 43.9 (C12), 22.0 ($CH_3CO_2^-$) ppm.

**HRMS** (ESI$^+$) calc. for $C_{23}H_{33}N_5O_{12}$: 572.2199 [M + H]$^+$; found: 572.2200.

#### 3.2 Regioselective protection of 12 towards cyclopentene precursors 13-15

##### (9H-fluoren-9-yl)methyl((1S,4S,5R)-4,5-*para*-methoxybenzylidene-2-en-1-yl)carbamate 29

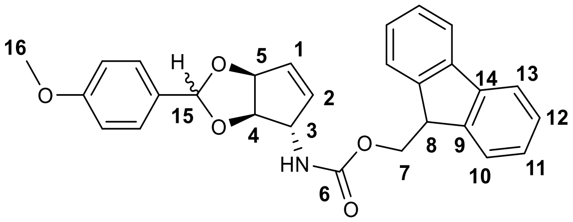

**12** (1.50 g, 4.50 mmol, 1.00 eq) was dissolved in DMF (75 mL) and anisaldehyde dimethyl acetal (0.92 mL, 984 mg, 5.40 mmol, 1.20 eq) and racemic camphor-10-sulfonic acid (105 mg, 0.45 mmol, 0.10 eq) were added subsequently. The reaction was stirred at room temperature for 2 h. Saturated aqueous $NaHCO_3$-solution was added and the phases were separated. After extracting the aqueous phase with DCM (3 x 200 mL), the organic phases were dried over $Na_2SO_4$, filtered and the solvent was evaporated *in vacuo*. Column chromatographic purification (EtOAc/*iso*-hexane 1:5 → EtOAc) gave the product (2.00 g, 4.39 mmol, 98%, 2 Isomers, A:B = 1:0.35) as colorless solid.

**$R_f$** (EtOAc/*iso*-hexane 1:3) = 0.30 (A+B, copolar)

**$^1$H-NMR** (600 MHz, $CDCl_3$): δ = 7.77 (d, *J* = 7.6 Hz, 2H, C13H), 7.58 (d, *J* = 7.4 Hz, 2H, C10H), 7.39 (m, 4H, C12H, arom. CH), 7.32 (m, 2H, C11H), 6.88 (m, 2H, arom. CH), 6.07 (m, 1 H, C1H (Isomer A + Isomer B)), 5.99 (m, 1H, C2H (Isomer B), 5.81 (m, 1H, C2H (Isomer B)), 5.85 (s, 1H, C15H (Isomer A)), 5.65 (s, 1H, C15H (Isomer B)), 5.49 (m, 1H, C5H), 4.76 (m, 2H, C3H), 4.59 (m, 1H, C4H), 4.46 (d, J = 6.8 Hz, 2H, C7H), 4.22 (m, 1H, C8H), 3.81 (s, 3H, $OCH_3$ (Isomer B)), 3.80 (s, 3H, $OCH_3$ (Isomer A) ppm.

**$^{13}$C-NMR** (150 MHz, $CDCl_3$): δ = 160.7 (arom. C), 155.8 (C6), 144.0 (C9), 141.5 (C14), 132.6 (C2 (Isomer A), 135.6 (C1, Isomer A + B), 134.8 (C1, Isomer A), 129.0 (arom. C), 128.4 (arom. C), 127.9 (C12), 127.2 (C11), 125.1 (C10), 120.2 (C13), 113.9 (arom. C), 105.6 (C15, Isomer A), 101.8 (C15, Isomer B), 85.4 (C4 Isomer A), 84.8 (C5), 84.0 (C4 Isomer B), 66.8 (C7), 62.4 (C3), 55.5 ($OCH_3$), 47.4 (C8) ppm.

**HRMS** (ESI$^+$) calc. for $C_{28}H_{25}NO_5Na$ [M + Na]$^+$: 478.1630; found: 478.1629.

**(9H-fluoren-9-yl)methyl((1S,4S,5R)-5-hydroxy-4-(*para*-methoxybenzyl)oxycyclopent-2-en-1-yl)carbamate 13**

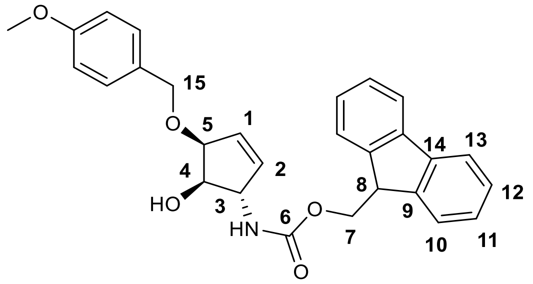

**29** (2.00 g, 4.39 mmol, 1.00 eq) was dissolved in DCM (60 mL) and cooled to -78 °C (dry ice/acetone). To ensure temperature equilibration, the mixture was stirred for 30 min. Then DIBAL-H (35.1 mL, 1 M solution in hexane, 35.1 mmol, 8.00 eq) was added slowly over 1.5 h ensuring that the temperature stays constant. After complete addition, the reaction mixture was stirred for another 1.5 h at -78 °C. Next, saturated aqueous Rochelle salt solution (30 mL) was added and the mixture was warmed to room temperature. Then, more DCM (150 mL) and Rochelle salt solution (500 mL) was added and the phases were separated. The aqueous phase was extracted with DCM (3 x 150 mL) and the organic phases were dried over $Na_2SO_4$, filtered and the solvent was evaporated. Column chromatographic purification (EtOAc/*iso*-hexane 3:1) gave the product (1.71 g, 3.73 mmol, 85%) as colorless solid.

**$R_f$** (EtOAc/*iso*-hexane 1:3) = 0.22

**$^1$H-NMR** (600 MHz, $CDCl_3$): δ = 7.76 (d, *J* = 7.7 Hz, 2H, C13H), 7.59 (m, 2H, C10H), 7.40 (t, *J* = 7.4 Hz, 2H, C12H), 7.31 (tt, $J_1$ = 7.4 Hz, $J_2$ = 1.1 Hz, 2H, C11H), 7.26 (m, 2H (overlapping with solvent peak), arom. CH), 6.89 (d, *J* = 8.3 Hz, arom. CH), 5.97 (m, 1H, C1H), 5.93 (m, 1H, C2H), 4.81 (d, *J* = 7.7 Hz, 1H, C4OH), 4.61 (m, 1H, C3H), 4.56 (m, 2H, C15H), 4.43 (m, 3H, C5H, C7H), 4.22 (m, 1H, C8H), 3.99 (m, 1H, C4H), 3.81 (s, 3H, $OCH_3$) ppm.

**$^{13}$C-NMR** (150 MHz, $CDCl_3$): δ = 159.5 (arom. C), 156.2 (C6), 144.0 (C9), 141.5 (C14), 136.0 (C2), 132.0 (C1), 125.2 (C10), 120.1 (C13), 127.8 (C12), 127.2 (C11), 129.6 (arom. C), 129.0 (arom. C), 114.1 (arom. C), 80.02 (C5), 77.3 (C4), 71.8 (C15), 66. 8 (C7), 62.8 (C3), 55.3 ($OCH_3$), 47.2 (C8) ppm.

**HRMS** ($ESI^+$) calc. for $C_{28}H_{27}NO_5Na$: 480.1787 $[M + Na]^+$; found: 480.1785.

**(9H-fluoren-9-yl)methyl((1S,4S,5R)-4-(*para*-methoxybenzyl)oxy-5-((2-trimethylsilylethoxy)-methyl)oxy-cyclopent-2-en-1-yl)carbamate 30**

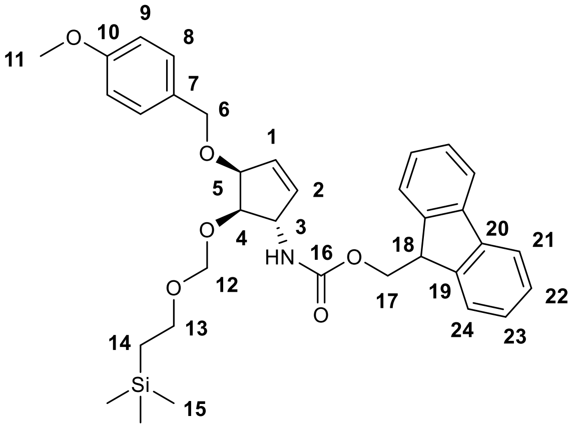

**13** (800 mg, 1.75 mmol, 1.00 eq) was dissolved in DMF (16 mL) and SEM-Cl (1.55 mL, 1.46 g, 8.74 mmol, 5.00 eq), pyridine (2.11 mL, 26.2 mmol, 15.0 eq) and $NBu_4I$ (64.6 mg, 0.175 mmol, 0.10 eq) were added subsequently. The reaction was stirred at 70 °C for 18 h before the solvent was evaporated *in vacuo*. Column chromatographic purification (EtOAc/*iso*-hexane 5:1 → 4:1) gave the product (767 mg, 1.30 mmol, 75%) as colorless solid.

**$R_f$** (EtOAc/*iso*-hexane 1:4) = 0.15

**$^1$H-NMR** (600 MHz, $CDCl_3$): δ = 7.76 (d, *J* = 7.5 Hz, 1H, C21H), 7.59 (t, *J* = 6.0 Hz, 1H, C24H ), 7.40 (t, *J* = 7.5 Hz, 1H, C22H), 7.31 (tt, $J_1$ = 7.5 Hz, $J_2$ = 1.3 Hz, 2H, C23H), 7.26 (m, 2H, arom. CH (PMB)), 6.87 (d, *J* = 8.4 Hz, 2H, arom. CH (PMB)), 6.01 (s, 1H, C2H), 5.95 (s, 1H, C1H), 4.88 (d, *J* = 4.8 Hz, 1H, C12Ha), 4.78 (m, 2H, C12Hb, C3H), 4.60 (d, *J* = 11.6 Hz, 1H, C6Ha), 4.51 (d, *J* = 11.6 Hz, 1H, C6Hb), 4.45 (m, 2H, C5H, C17Ha), 4.36 (m, 1H, C17Hb), 4.21 (m, 1H, C18H), 3,98 (m, 1H, C4H), 3.80 (s, 3H, C11H), 3.71 (m, 2H, C13H), 0.94 (t, *J* = 8.6 Hz, 2H, C14H), -0.02 (s, 9H, -$Si(CH_3)_3$) ppm.

**$^{13}$C-NMR** (125 MHz, $CDCl_3$): δ = 159.2 (C10), 155.8 (C16), 143.9 (C19), 141.3 (C20), 135.9 (C2), 131.7 (C1), 130.5 (C7), 129.3 (C8), 127.7 (C22), 127.0 (C23), 125.0 (C24), 120.0 (C21), 113.7 (C9), 94.3 (C12), 81.3 (C4), 78.6 (C5), 70.7 (C6), 66.7 (C17), 65.7 (C13), 60.6 (C3), 55.2 (C11), 47.2 (C18), 18.1 (C14), -1.50 (C15) ppm.

**HRMS** (ESI$^+$) calc. for $C_{34}H_{41}NO_6SiNa$ $[M + Na]^+$: 610.2601; found: 610.2599.

**(9H-fluoren-9-yl)methyl((1S,4S,5R)-4-hydroxy-5-((2-trimethylsilylethoxy)-methyl)oxy-cyclopent-2-en-1-yl)carbamate 14**

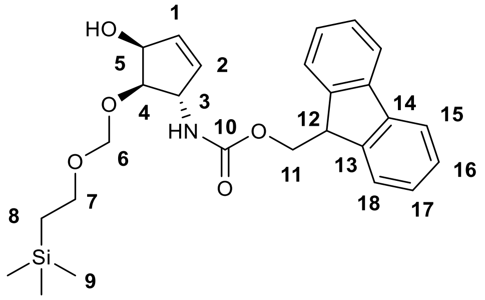

**30** (506 mg, 0.86 mmol) was dissolved in DCM (22.5 mL) and cooled to 0 °C. The mixture was stirred at 0 °C for 30 min to ensure temperature equilibration. Next, TFA (2.5 mL) was added and the reaction was stirred at 0 °C for 5 min. The reaction was stopped by first addition of $HNEt_3OAc$ (1 M in DCM, 20 mL) and then saturated aqueous $NaHCO_3$-solution in quick succession. The phases were separated and the aqueous phase was extracted with DCM (3 x 50 mL). The organic phases were dried over $MgSO_4$, filtered and the solvent was evaporated *in vacuo*. Column chromatographic purification (EtOAc/*iso*-hexane 2:1) gave the product (335 mg, 0.71 mmol, 83%) as colorless solid.

$\mathbf{R_f}$ (EtOAc/*iso*-hexane 1:2) = 0.20

**$^1$H-NMR** (600 MHz, $CDCl_3$): δ = 7.77 (dt, $J_1$ = 7.5 Hz, $J_2$ = 0.9 Hz, 2H, C15H), 7.59 (m, 2H, C18H), 7.40 (m, 2H, C16H), 7.32 (td, $J_1$ = 7.4 Hz, $J_2$ = 1.1 Hz, 2H, C17H), 6.03 (m, 1H, C1H), 5.92 (m, 1H, C2H), 4.85 (m, 2H, C5H, C6Ha), 4.71 (m, 2H, C3H, C6Hb), 4.43 (m, 2H, C11H), 4.22 (m, 1H, C12H), 3.95 (m, 1H, C4H), 3.68 (m, 2H, C7H), 0.96 (t, J = 8.6 Hz, 2H, C8H), 0.01 (s, 9H, $(SiCH_3)_3$) ppm.

**$^{13}$C-NMR** (125 MHz, $CDCl_3$): δ =144.2 (C13), 141.6 (C14), 135.2 (C2), 134.2 (C1), 128.0 (C16), 127.4 (C17), 125.3 (C18), 120.3 (C15), 95.1 (C6), 82.2 (C4), 73.6 (C5), 67.0 (C11), 66.4 (C7), 60.9 (C3), 47.5 (C12), 18.4 (C8), -1.15 $(SiCH_3)_3$) ppm.

**HRMS** (ESI$^+$) calc. for $C_{26}H_{33}NO_5SiNa$: 490.2026 [M + Na]$^+$; found: 490.2024.

**(9H-fluoren-9-yl)methyl((1S,4S,5R)-5-((tert-butyldimethylsilyl)oxy)-4-hydroxycyclopent-2-en-1-yl)carbamate 15**

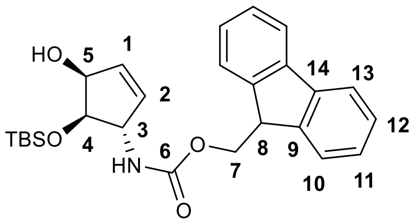

**12** (200 mg, 0.59 mmol, 1.00 eq) was dissolved in DMF (6 mL) and cooled to -10 °C (NaCl/ice bath). Next, TBSOTf (163 µL, 188 mg, 0.71 mmol, 1.20 eq) was added and the reaction was stirred for 2 h at -10 °C. Saturated aqueous $NaHCO_3$-solution was added and the phases were separated. After extracting the aqueous phase with DCM (3 x 50 mL), the organic phases were dried over $Na_2SO_4$, filtered and the solvent was evaporated *in vacuo*. Column chromatographic purification (EtOAc/*iso*-hexane 1:5) gave the product (110 mg, 0.24 mmol, 41%) as colorless solid.

**$R_f$** (EtOAc/*iso*-hexane 1:3) = 0.12.

**$^1$H-NMR** (600 MHz, $CDCl_3$): δ = 7.77 (d, *J* = 7.5 Hz, 2H, C13H), 7.58 (d, *J* = 7.5 Hz, 2H, C10H), 7.40 (t, *J* = 7.5 Hz, 2H, C12H), 7.31 (t, *J* = 7.5 Hz, 2H, C11H), 6.00 (d, *J* = 5.7 Hz, 1H, C1H), 5.83 (d, *J* = 6.0 Hz, 1H, C2H), 4.56 (m, 2H, C3H, C5H), 4.43 (m, 2H, C6H), 4.21 (t, *J* = 6.8 Hz, 1H, C7H), 4.04 (m, 1H, C4H), 0.92 (s, 9H, $SiC(CH_3)_3$), 0.17 (s, 3H, $SiCH_3$), 0.14 (s, 3H, $SiCH_3$) ppm.

**$^{13}$C-NMR** (150 MHz, $CDCl_3$): δ = 155.8 (C6), 144.0 (C9), 141.5 (C14), 135.8 (C1), 133.5 (C2), 127.8 (C12), 127.2 (C11), 125.1 (C10), 120.1 (C13), 77.5 (C4), 74.2(C5), 66.7 (C7), 62.6 (C3), 47.4 (C8), 25.9 ($SiC(CH_3)_3$)), 18.2 ($SiC(CH_3)_3$)), -4.5 ($SiCH_3$), -5.0 ($SiCH_3$) ppm.

**$^{29}$Si-NMR** (80 MHz, $CDCl_3$): δ = 23.2 ppm.

**HRMS** ($ESI^+$) calc. for $C_{26}H_{37}N_2O_4Si$: 469.2517 $[M + NH_4]^+$; found: 469.2520.

##### 3.3 Synthesis of glycosyl donor 17

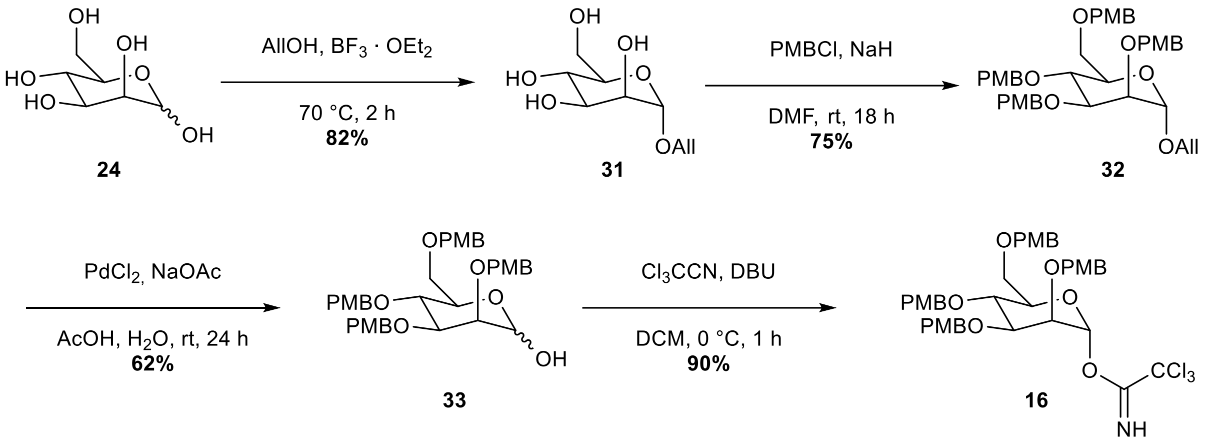

**Supplementary Figure 5:** Synthesis of glycosyl donor **16** in 4 steps starting from D-mannose (overall yield: 34%)

##### 1-*O*-Allyl-D-mannopyranose (31)[6]

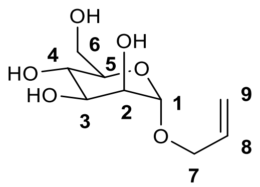

D-Mannose (5.00 g, 27.8 mmol, 1.00 eq) was suspended in AllOH (25 mL) and $BF_3 \cdot OEt_2$ (343 µL, 2.78 mmol, 0.10 eq) was added. The reaction was heated to 70 °C for 2 h. After cooling to room temperature, the mixture was neutralized with $NEt_3$ and the solvent was evaporated *in vacuo*. Column chromatographic purification (MeOH/DCM 1:10) gave the product (5.02 g, 22.8 mmol, 82%) as colorless gum.

**$R_f$** (MeOH/DCM 1:5) = 0.46

**$^1$H-NMR** (600 MHz, $d^4$-MeOH): δ = 5.98 (m, 1H, C8H), 5.34 (dq, $J_1$ = 17.2 Hz, $J_2$ = 1.7 Hz, 1H, C9Ha), 5.21 (m, 1H, C9Hb), 4.83 (d, $J$ = 1.7 Hz, 1H, C1H), 4.26 (ddt, $J_1$ = 13.1 Hz, $J_2$ = 5.1 Hz, $J_3$ = 1.5 Hz, 1H, C7Ha), 4.05 (ddt, $J_1$ = 13.1 Hz, $J_2$ = 6.0 Hz, $J_3$ = 1.4 Hz, 1H, C7Hb), 3.87 (dd, $J_1$ = 11.8 Hz, $J_2$ = 2.4 Hz, 1H, C6Ha), 3.85 (dd, $J_1$ = 3.4 Hz, $J_2$ = 1.7 Hz, 1H, C2H), 3.75 (m, 2H, C3H, C6Hb), 3.65 (t, $J$ = 9.5 Hz, 1H, C4H), 3.58 (m, 1H, C5H) ppm.

**$^{13}$C-NMR** (125 MHz, $d^4$-MeOH): δ = 135.5 (C8), 117.2 (C9), 100.8 (C1), 74.8 (C5), 72.7 (C3), 72.6 (C2), 68.8 (C7), 68.7 (C4), 63.4 (C6) ppm.

**HRMS** (ESI$^+$) calc. for $C_9H_{16}O_6Na$ $[M + Na]^+$: 243.0845; found: 243.0838.

**1-*O*-Allyl-2,3,4,6-*O*-(*para*-methoxybenzyl)-D-mannopyranose (32)**

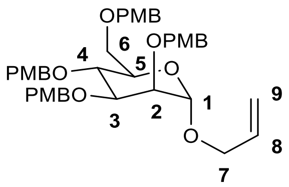

**31** (1.84 g, 8.36 mmol, 1.00 eq) was dissolved in DMF (19 mL). NaH (2.00 g, 50.1 mmol, 6.00 eq) was added at 0 °C and the reaction was stirred for 30 min before PMB-Cl (6.80 mL, 7.85 g, 50.1 mmol, 6.00 eq) was added dropwise. The mixture was stirred at room temperature for 18 h and the poured into saturated aqueous $NaHCO_3$-solution. The solution was extracted with DCM (3 x 200 mL), the combined organic phases were dried over $Na_2SO_4$, filtered and the solvent was evaporated *in vacuo*. Column chromatographic purification (EtOAc/*iso*-hexane 1:4 → 1:3) yielded the product (4.39 g, 6.27 mmol, 75%) as colorless syrup.

$\mathbf{R_f}$ (EtOAc/*iso*-hexane 1:3) = 0.15

$^1$**H-NMR** (400 MHz, $CDCl_3$): δ = 7.28 (m, 6H, arom. CH), 7.04 (m, 2H, arom. CH), 6.83 (m, 8H, arom. CH), 5.83 (m, 1H, C8H), 5.20 (dd, $J_1$ = 10.4 Hz, $J_2$ = 1.4 Hz, 1H, C9Ha), 5.13 (dd, $J_1$ = 10.4 Hz, $J_2$ = 1.4 Hz, 1H, C9Hb), 4.87 (d, J = 1.8 Hz, 1H, C1H), 4.77 (d, J = 10.3 Hz, 1H, $CH_2$), 4.63 (m, 3H, $CH_2$), 4.51 (m, 3H, $CH_2$), 4.38 (d, J = 10.2 Hz, 1H, $CH_2$), 4.15 (m, 1H, C7Ha), 3.90 (m, 3H, C4H, C5H, C7Hb), 3.81(s, 3H, $OCH_3$), 3.80(s, 3H, $OCH_3$), 3.79 (s, 3H, $OCH_3$), 3.79 (s, 3H, $OCH_3$), 3.75 (dd, $J_1$ = 2.7 Hz, $J_2$ = 1.9 Hz, 1H, C2H), 3.69 (m, 3H, C3H, C6H) ppm.

$^{13}$**C-NMR** (125 MHz, $CDCl_3$): δ = 159.3 (2C, arom. C), 159.1 (2C, arom. C), 134.0 (C8), 130.9 (arom. C), 130.8 (arom. C), 130.6 (arom. C), 130.6 (arom. C), 129.8 (arom. C), 129.6 (2C, arom. C), 129.3 (arom. C), 117.2 (C9), 113.8 (arom. C), 113.8 (arom. C), 113.8 (2C, arom. C), 97.2 (C1), 80.1 (C5), 74.9 ($CH_2$), 74.8 (2C, C4, $CH_2$), 74.2 (C2), 72.2 ($CH_2$), 72.0 ($CH_2$), 71.9 (C3), 68.9 (C6), 67.8 (C7), 55.4 ($OCH_3$), 55.4 ($OCH_3$), 55.4 ($OCH_3$), 55.4 ($OCH_3$) ppm.

**HRMS** ($ESI^+$) calc. for $C_{51}H_{42}NO_{10}$: 718.3586 $[M + NH_4]^+$ ; found: 718.3584;

**2,3,4,6-*O*-(*Para*-methoxybenzyl)-D-mannopyranose (33)[7]**

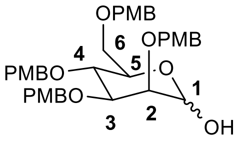

**32** (2.70 g, 3.85 mmol, 1.00 eq) was dissolved in AcOH (20 mL) and $H_2O$ (0.65 mL). NaOAc (886 mg, 10.8 mmol, 2.80 eq) and $PdCl_2$ (887 mg, 5.00 mmol, 1.30 eq) were added and the reaction was stirred at room temperature for 24 h. The resulting mixture was diluted with DCM and filtered over celite. The filtrate was washed with saturated aqueous $NaHCO_3$-solution (50 mL) and brine (50 mL), dried over $MgSO_4$, filtered and the solvent was evaporated *in vacuo*. Column chromatographic purification (EtOAc/*iso*-hexane 1:1.5) gave the product (1.57 g, 2.38 mmol, 62%, α:β = 1:0.26) as colorless syrup.

$R_f$ (EtOAc/*iso*-hexane 1:1) = 0.36

**$^1$H-NMR** (600 MHz, $CDCl_3$): δ = 7.26 (m, 6H, arom. CH), 7.05 (d, J = 8.3 Hz, 2H, arom. CH), 6.84 (m, 8H, arom. CH), 5.20 (m, 1H, C1Hα), 5.00 (d, 2H, J = 11.5 Hz, $CH_2$α), 4.88 (s, 1H, C1Hβ), 4.58 (m, 8H, $CH_2$), 3.97 (ddd, $J_1$ = 9.8 Hz, $J_2$ = 6.5 Hz, $J_3$ = 2.0 Hz, 1H, C5Hα), 3.90 (dd, $J_1$ = 9.3 Hz, $J_2$ = 3.0 Hz, 1H, C3Hα), 3.86 (m, 1H, C4Hα), 3.80 (s, 3H, $OCH_3$), 3.79 (m, 7H, C4Hα, 2 x $OCH_3$), 3.78 (s, 3H, $OCH_3$), 3.75 (t, J = 2.5 Hz, 1H, C2Hα), 3.66 (m, C6Hαa, $CH_2$), 3.61 (dd, $J_1$ = 10.4 Hz, $J_2$ = 6.5 Hz, 1H, C6Hαb) ppm.

**$^{13}$C-NMR** (600 MHz, $CDCl_3$): δ = 159.3, 159.3, 159.3, 159.3, 159.3, 113.9, 113.9, 113.9, 113.9, 113.8, 130.9, 130.8, 130.6, 130.4, 130.4, 130.3, 130.0, 129.8, 129.8, 129.8, 129.7, 129.6, 129.4, 129.4, 129.4 (arom. C), 97.9 (C1β), 93.0 (C1α), 79.5 (C3α), 75.8, 75.4, 75.0 (C4α), 74.8 ($CH_2$), 74.8, 74.4, 74.4 (C2α), 73.1, 73.1 ($CH_2$), 72.5, 72.4, 72.4 ($CH_2$), 72.0 ($CH_2$), 71.9, 71.9 (C5α), 71.6, 69.3 (C6α), 68.7, 55.4, 55.4, 55.4 ($OCH_3$) ppm.

**NOTE:** Signals for β-Anomer cannot be assigned unambiguously due to the low abundance in the mixture and overlapping signals.

**HRMS** (ESI$^+$) calc. for $C_{38}H_{44}O_{10}Na$: 683.3832 $[M + Na]^+$; found: 683.2831.

Analytical data in accordance with literature.

**2,3,4,6-*O*-(*Para*-methoxybenzyl)-α-D-mannopyranosyltrichloroacetimidate (16)**

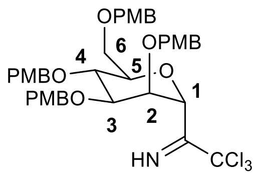

**33** (720 mg, 1.09 mmol, 1.00 eq) was dissolved in DCM (12 mL). $Cl_3CCN$ (1.09 mL,1.57 g, 10.9 mmol, 10.0 eq) and DBU (33.0 µL, 33.5 mg, 0.22 mmol, 0.20 eq) were added at 0 °C and the reaction was stirred at the same temperature for 1 h. The solvent was evaporated *in vacuo*. Column chromatographic purification (EtOAc/*iso*-hexane 1:3) gave the product (789 mg, 0.98 mmol, 90%) as colorless syrup. (NOTE: a slight yellow color indicates amine impurities, which hamper glycosylation)

**$R_f$** (EtOAc/*iso*-hexane 1:3) = 0.32

**$^1$H-NMR** (600 MHz, $CDCl_3$): δ = 8.51 (s, 1H, NH), 7.33 (d, J = 8.6 Hz, 2H, arom. CH), 7.25 (m with solvent peak, 2H, arom. CH), 7.22 (d, J = 8.5 Hz, 2H, arom. CH), 7.09 (d, J = 8.5 Hz, 2H, arom. CH), 6.83 (m, 8H, arom. CH), 6.33 (d, J = 2.0 Hz, 1H, C1H), 4.79 (d, J = 10.2 Hz, 1H, $CH_2$), 4.69 (m, 2H, $CH_2$), 4.61 (d, J = 11.7 Hz, 1H, $CH_2$), 4.52 (m, 2H, $CH_2$), 4.44 (m, 2H, $CH_2$), 4.06 (t, J = 9.7 Hz, 1H, C4H), 3.92 (ddd, $J_1$ = 9.8 Hz, $J_2$ = 4.5 Hz, $J_3$ = 1.7 Hz, 1H, C5H), 3.87 (dd, $J_1$ = 9.5 Hz, $J_2$ = 3.2 Hz, 1H, C3H), 3.81 (m, 1H, C2H), 3.80 (s, 3H, $OCH_3$), 3.79 (s, 3H, $OCH_3$), 3.79 (s, 3H, $OCH_3$), 3.78 (s, 3H, $OCH_3$), 3.76 (m, 1H, C6Ha), 3.67 (dd, $J_1$ = 11.2 Hz, $J_2$ = 1.9 Hz, 1H, C6Hb) ppm.

**$^{13}$C-NMR** (600 MHz, $CDCl_3$): δ = 160.6 (OCNH), 159.4, 159.4, 159.4, 159.3, 130.6, 130.6, 130.4, 130.2, 130.0 129.8, 129.7, 129.6, 113.9, 113.9, 113.9, 133.8 (all aromatic C), 96.3 (C1), 91.3 ($CCl_3$), 75.1 ($CH_2$), 75.0 (C5), 74.0 (C4), 73.1 ($CH_2$), 73.1 (C2), 78.6 (C3), 72.3 ($CH_2$), 72.0 ($CH_2$), 68.5 (C6), 55.4, 55.4, 55.4, 55.4 (all $OCH_3$) ppm.

**HRMS** ($ESI^+$) calc. for $C_{40}H_{44}NO_{10}Cl_3Na$: 826.1928 $[M + Na]^+$; found: 826.1922.

#### 3.4 Synthesis of ManQ-Isomers 10, 11 and 4

**(9H-fluoren-9-yl)methyl((1S,4S,5R)-4-(((2R,4aR,6R,7S,8S,8aR)-7,8-bis((4-methoxybenzyl)oxy)-2-phenylhexahydropyrano[3,2-d][1,3]dioxin-6-yl)oxy)-5-((tert-butyldimethylsilyl)oxy)cyclopent-2-en-1-yl)carbamate 17**

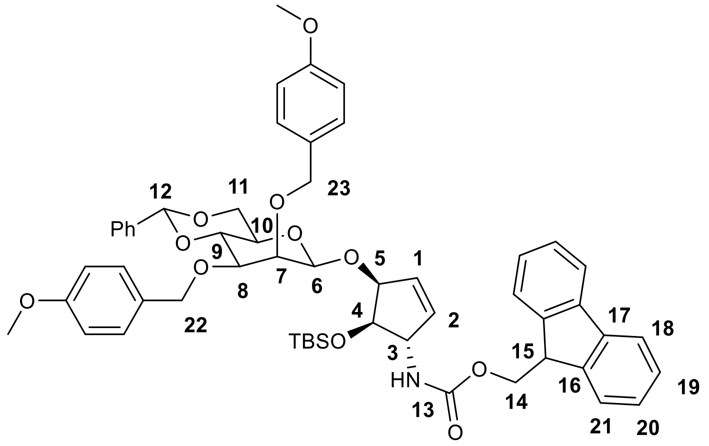

AgOTf (68.3 mg, 266 µmol, 3.00 eq) was dissolved in DCM (1 mL) and pulverized 3 Å molsieves (20 w% of the solvent volume) were added. The mixture was cooled to -78 °C before a solution of PhSCl (32.1 mg, 222 µmol, 2.50 eq) in DCM (1 mL) was added. The reaction was stirred for 5 min at – 78 °C. Next, a solution of sugar donor **5** (53.2 mg, 88.6 µmol, 1.00 eq) and 2,6-di-*tert*-butyl-4-methylpyridine (54.6 mg, 266 µmol, 3.00 eq) in DCM (1 mL) was added dropwise and the reaction was stirred for 15 min at -78 °C. Then, a solution of glycosyl acceptor **15** (40.0 mg, 88.6 µmol, 1.00 eq) in DCM (1 mL) was added and the reaction was kept stirring at – 78 °C for 2 h. A saturated aqueous $NaHCO_3$-solution (5 mL) and DCM (10 mL) were added and the reaction was warmed to room temperature before it was filtered over celite. The phases were separated and the organic phase was washed with brine and dried over $Na_2SO_4$. After filtration, the solvent was removed *in vacuo* and the residue was purified by column chromatography (EtOAc/*iso*-hexane 1:10 → 1:5 → 1:3) to yield the product (43.4 mg, 46.1 µmol, 52%) as colorless solid.

$\mathbf{R_f}$ (EtOAc/*iso*-hexane 1:3) = 0.12.

**$^1$H-NMR** (600 MHz, $CDCl_3$): δ = 7.78 (d, *J* = 7.6 Hz, 2H, C18H), 7.60 (t, *J* = 6.9 Hz, 2H, C21H), 7.51 (d, *J* = 7.0 Hz, 2H, arom. CH), 7.38 (m, xH, C19H, arom. CH), 7.32 (t, *J* = 7.4 Hz, 2H, C20H), 7.15 (d, *J* = 7.2 Hz, 2H, arom. CH), 6.88 (d, *J* = 8.2 Hz, 2H, arom. CH), 6.81 (d, *J* = 8.3 Hz, 2H, arom. CH), 6.00 (m, 1H, C1H), 5.88 (d, *J* = 6.2 Hz, 1H, C2H), 5.61 (s, 1H, C12H), 4.88 (d, *J* = 12.0 Hz, 1H, C23Ha), 4.79 (d, *J* = 12.0 Hz, 1H, C23Hb), 4.71 (m, 3H, C3H, C6H), 4.46 (m, 5H, C5H, C14H, C21H), 4.31 (dd, $J_1$ = 10.4 Hz $J_2$ = 4.8 Hz, C11Ha), 4.22 (t, *J* = 6.6 Hz, 1H, C15H), 4.17 (t, *J* = 9.6 Hz, C9H), 3.96 (m, 3H, C4H, C7H, C11Hb), 3.81 (s, 3H, $OCH_3$), 3.79 (s, 3H, $OCH_3$), 3.48 (dd, $J_1$ = 9.9 Hz, $J_2$ = 3.2 Hz, 1H, C8H), 3.27 (td, $J_1$ = 9.7 Hz, $J_2$ = 4.8 Hz, 1H, C10H), 0.81 (s, 9H, $SiC(CH_3)_3$), 0.04 (s, 3H, $SiCH_3$), 0.00 (s, 3H, $SiCH_3$) ppm.

**$^{13}$C-NMR** (150 MHz, $CDCl_3$): δ = 159.2 (2C, arom. C), 155.9 (C13), 144.0 (C15), 141.5 (C16), 137.0 (C1), 131.3 (C2), 130.7 (arom. C), 130.6 (arom. C), 130.5 (arom. C), 130.4 (arom. C), 129.3 (arom. C), 128.9 (arom. C), 128.3 (arom. C), 127.8 (C19), 127.2 (C20), 126.2 (arom C), 125.1 (C21), 120.1 (C18), 113.8 (arom. C), 113.7 (arom. C), 102.1 (C6), 101.4 (C12), 80.1 (C5), 78.7

(C7), 78.5 (C9), 77.5 (C8), 74.6 (C4), 74.4 (C23), 71.9 (C22), 67.8 (C10), 68.7 (C11), 66.8 (C14), 61.0 (C3), 55.4 ($OCH_3$), 55.3 ($OCH_3$),  47.4 (C15), 25.8 ($SiC(CH_3)_3$), 18.1 ($SiC(CH_3)_3$),  -4.5 ($SiCH_3$), -4.7 ($SiCH_3$) ppm.

**HRMS** ($ESI^+$) calc. for $C_{55}H_{67}N_2O_{11}Si$: 959.4509 $[M + NH_4]^+$; found: 959.4522.

**(9H-fluoren-9-yl)methyl((1S,4S,5R)-4-((4-methoxybenzyl)oxy)-5-(((2R,3R,4R,5S,6S)-3,4,5-tris((4-methoxybenzyl)oxy)-6-(((4-methoxybenzyl)oxy)methyl)tetrahydro-2H-pyran-2-yl)oxy)cyclopent-2-en-1-yl)carbamate 18**

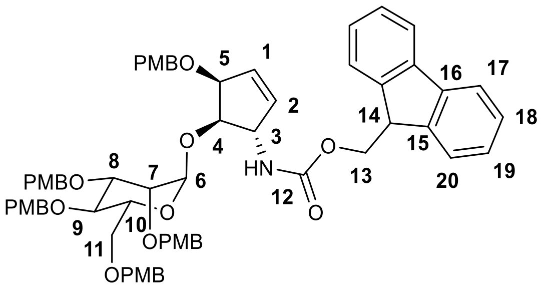

**13** (20.0 mg, 43.7 µmol, 1.00 eq) and **16** (42.2 mg, 62.5 µmol, 1.20 eq) were dissolved in THF (3 mL) and activated 3 Å molsieves (20 w%) were added. The solution was stirred at 0 °C for 30 min. TMSOTf (0.1 M in THF, 87.4 µL, 8.74 µmol, 0.20 eq, pre-dried over 3 Å molsieves for 30 min) was added dropwise and the reaction was stirred at 0 °C for 1 h. Saturated aqueous $NaHCO_3$-solution (3 mL) was added and the mixture was warmed to room temperature, diluted with DCM (20 mL) and $H_2O$ (20 mL), filtered and the phases were separated. The aqueous phase was re-extracted with DCM (3 x 20 mL), the combined organic phases were dried over $Na_2SO_4$, filtered and the solvent was evaporated *in vacuo*. Column Chromatographic purification (EtOAc/Hexane 1:4 → 1:2) gave the product (26.4 mg, 24.0 µmol, 55%) as colorless solid.

$\mathbf{R_f}$ (EtOAc/*iso*-hexane 1:3) = 0.12

**$^1$H-NMR** (600 MHz, $CDCl_3$): δ = 7.73 (d, *J* = 7.8 Hz, 2H, C17H), 7.57 (d, *J* = 7.6 Hz, 2H, C20H), 7.35 (t, *J* = 7.4 Hz, 2H, C18H), 7.22 (m, 10H, C19H, arom. CH), 7.04 (m, 2H, arom. CH),6.80 (m, 10H, arom. CH), 6.01 (m, 1H, C1H), 5.90 (m, 1H, C2H), 4.96 (m, 1H), 4.94 (m, $\boldsymbol{^1J_{C\text{-}H}}$ **= 170.3 Hz**, 1H, C6H), 4.77 (t, J = 9.9 Hz, 1H), 4.61 (d, J = 3.7 Hz, 1H), 4.47 (m,), 4.16 (m), 3.92 (m), 3.77 (m, $OCH_3$),

**NOTE:** Signals cannot be assigned unambiguously. $^{13}$C-NMR-data are not assigned, the spectrum can be found in the spectra-section.

**HRMS** (ESI$^+$) calc. for $C_{66}H_{77}N_3O_{14}$: 1117.5056 $[M + NH_4]^+$ ; found: 1117.5084.

**(9H-fluoren-9-yl)methyl((1S,4S,5R)-5-((2-(trimethylsilyl)ethoxy)methoxy)-4-(((2R,3R,4R,5S,6S)-3,4,5-tris((4-methoxybenzyl)oxy)-6-(((4-methoxybenzyl)oxy)methyl)-tetrahydro-2H-pyran-2-yl)oxy)cyclopent-2-en-1-yl)carbamate 19**

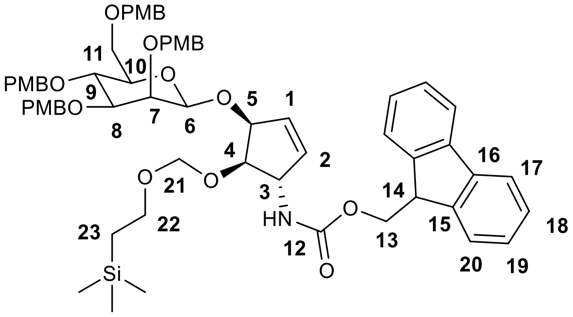

**14** and **16** were dissolved in THF (2 mL) and 3 Å molsieves (20 w% of solvent) were added. The mixture was stirred at 0 °C for 20 min before a solution of TMSOTf (231 µL, 0.1 M in THF) was added. The reaction was stirred at 0 °C for 1.5 h before saturated aqueous $NaHCO_3$-solution (5 mL) was added. DCM (20 mL) was added and the phases were separated. The aqueous phase was re-extracted with DCM (3 x 20 mL). The combined organic phases were dried over $Na_2SO_4$, filtered and the solvent was evaporated *in vacuo*. Column chromatographic purification (EtOAc/*iso*-hexane 1:4 → 1:3) gave an anomeric mixture of the product as colorless solid.

**$R_f$** (EtOAc/*iso*-hexane 1:3) = 0.14

**$^1$H-NMR** (600 MHz, $CDCl_3$): δ = 7.79 (dd, $J_1$ = 7.6 Hz $J_2$ = 3.8 Hz, 2H, C17H), 7.61 (m, 2H, C20H), 7.42 (m, 2H, C18H), 7.33 (t, $J$ = 7.6 Hz, 1H, C19H), 7.28 (m with solvent peak, arom. CH), 7.07 (m, 2H, arom. CH), 6.85 (m, arom. CH), 6.03 (m, 1H, C1H), 5.99 (m, 1H, C2H), 5.01 (m, 1H), 4.87 (m, 1H, C6H), 4.62 (m, 18H), 4.25 (m, 1H), 3.79 (m, $OCH_3$), 3.54 (m, 1H), 3.43 (m, 1H), 0.95 (t, $J$ = 7.3 Hz, 1H, C23H), -0.01 (s, 3H, $SiCH_3$), -0.02 (s, 6H, $SiCH_3$) ppm.

**NOTE:** Due to the received inseparable anomeric mixture, $^1$H-NMR data refer to the most important features proofing the formation of the glycosylation adduct from the starting materials. The anomeric mixture was only separated after completing the synthesis via HPLC. $^{13}$C-NMR-data are not assigned, but can be found in the spectra-section.

**HRMS** (ESI$^+$) calc. for $C_{64}H_{76}NO_{14}Si$: 1110.5030 [M + H]$^+$; found: 1110.5033.

**β-allyl-Mannosyl-Queuosine 10**

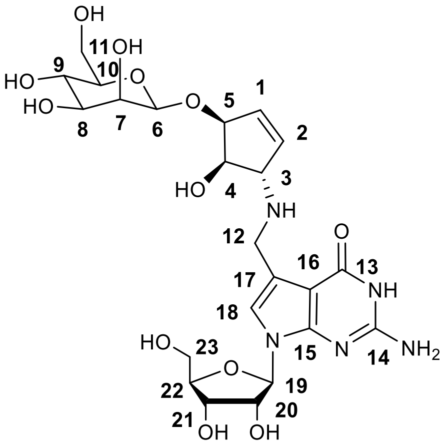

**17** (40 mg, 42.4 µmol) was dissolved in MeCN (1.5 mL) and DBU (224 µL) was added. The reaction was stirred for 1 h at room temperature and then neutralized with AcOH. The solvent was evaporated in vacuo and the crude residue purified by column chromatography (MeOH/DCM 1:20). The resulting product (21 mg) was identified by LC-MS and subsequently used for the next step. For that, it was dissolved in MeOH (0.25 mL) and **8** (21 mg, 1.00 eq) was added. AcOH (1.00 µL) was added and the reaction was stirred for 2 h at room temperature. Next the reaction was cooled to 0 °C and $NaBH_4$ (2.87 mg, 75.9 µmol, 2.60 eq) was added. The reaction was stirred at 0 °C for 15 min before the solvent was evaporated. Product formation was confirmed by LC-MS. The residue was dissolved in DCM (0.9 mL) and TFA (0.1 mL) was added at 0 °C. The reaction was stirred at 0 °C for 20 min before it was neutralized with $NEt_3$. After evaporation of the solvent, the product was detected by LC-MS and then re-dissolved in EtOAc (0.5 mL). HF · pyridine (15 µL) was added and the solution was stirred for 1 d before the reaction was stopped by addition of TMSOMe (46 µL). The solvent was evaporated, the crude product was identified by LC-MS and dissolved in MeOH (1 mL). NaOMe (27.0 mg, 0.5 mmol) was added and the reaction was stirred for 2 d until LC-MS indicated completion. The mixture was neutralized with $NEt_3$ and the solvent was evaporated *in vacuo*. The residue was dissolved in water, filtered and purified by reversed phase HPLC (0-10 % buffer B, 45 min) to afford the product **10** (1.30 mg, 2.33 µmol, 5% over 6 steps) as colorless solid.

**RP-HPLC:** $R_t$ (C18-column, 0-10% buffer B, 45 min) = 21.7 min.

**$^1$H-NMR** (800 MHz, $D_2O$): δ = 7.13 (s, 1H, C18H), 6.25 (m, 1H, C1H), 6.10 (dd, $J_1$ = 6.3 Hz, $J_2$ = 1.7 Hz, 1H, C2H), 5.97 (d, $J$ = 6.5 Hz, 1H, C19H), 4.70 (m, covered by solvent peak, 2H, C5H, C6H (**$^1J_{C-H}$ = 160.2 Hz**)), 4.58 (dd, $J_1$ = 6.5 Hz, $J_2$ = 5.4 Hz, 1H, C20H), 4.41 (m, 3H, C4H, C12H), 4.32 (dd, $J_1$ = 5.4 Hz, $J_2$ = 3.3 Hz, 1H, C21H), 4.27 (m, 1H, C3H), 4.16 (q, $J$ = 3.6 Hz, 1H, C22H), 4.04 (d, $J$ = 3.1 Hz, 1H, C7H), 3.88 (dd, $J_1$ = 12.3 Hz, $J_2$ = 2.3 Hz, 1H, C11Ha), 3.82 (dd, $J_1$ = 12.6 Hz, $J_2$ = 3.3 Hz, 1H, C23Ha), 3.77 (dd, $J_1$ = 12.6 Hz, $J_2$ = 4.3 Hz, 1H, C23Hb), 3.70 (dd, $J_1$ = 12.3 Hz, $J_2$ = 6.6 Hz, 1H, C11Hb), 3.62 (dd, $J_1$ = 9.6 Hz, $J_2$ = 3.3 Hz, 1H, C8H), 3.54 (t, $J$ = 9.7 Hz, 1H, C9H), 3.36 (ddd, $J_1$ = 9.3 Hz, $J_2$ = 6.5 Hz, $J_3$ = 2.3 Hz, 1H, C10H) ppm.

**$^{13}$C-NMR** (HSQC, HMBC, $D_2O$): δ = 151.7 (C14/C15), 152.7 (C14/C15), 136.1 (C1), 130.0 (C2), 118.9 (C18), 100.2 (C16/17), 98.8 (C16/17), 99.8 (C6), 86.6 (C19), 84.8 (C22), 80.7 (C5), 76.3

(C10), 73.7 (C4), 73.4 (C20), 72.8 (C11), 70.4 (C21), 70.3 (C7), 66.7 (C9), 66.4 (C3), 61.4 (C23), 60.9 (C11), 42.0 (C12) ppm.

**HRMS** ($ESI^+$) calc. for $C_{23}H_{34}N_5O_{12}$: 572.2198 $[M + H]^+$; found: 572.2197.

**α-homoallyl-Mannosyl-Queuosine 11**

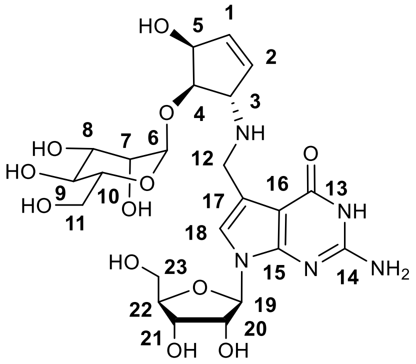

**18** (19.0 mg, 17.2 µmol) was dissolved in MeCN (1 mL) and DBU (149 µL, 1.00 mmol) was added. The reaction was stirred at room temperature for 1 h and then neutralized with AcOH. The solvent was evaporated in vacuo to give the free amine, which was detected by LC-MS. The product was dissolved in MeOH and **8** (12.0 mg, 13.7 µmol) was added together with AcOH (1.00 µL). The reaction was stirred for 5 h before it was cooled to 0 °C and $NaBH_4$ (1.34 mg, 35.6 µmol) was added. The reaction was stirred at 0 °c for 20 min, then $H_2O$ ( 0.5 mL) was added and the reaction was stirred for another 5 min before the solvent was evaporated. The crude product was detected by LC-MS and next dissolved in DCM (0.9 mL) and TFA (0.1 mL) was added at 0 °C. The reaction was stirred for 30 min and then neutralized with $NEt_3$. After evaporation of the solvent, the crude PMB-deprotected product was detected by LC-MS and then dissolved in MeOH (1 mL). NaOMe (27.0 mg, 0.5 mmol) was added and the reaction was stirred at room temperature for 2 d. After neutralization with HCl (2 M), the solvent was evaporated *in vacuo*. The residue was dissolved in water, filtered and purified by reversed phase HPLC (0-10 % buffer B, 45 min) to afford the product **11** (1.51 mg, 2.64 µmol, 15% over 5 steps) as colorless solid.

**RP-HPLC:** $R_t$ (C18-column, 0-10% buffer B, 45 min) = 21.3 min.

**$^1$H-NMR** (600 MHz, $d^3$-MeOD): δ = 7.04 (s, 1H, C18H), 6.11 (ddd, $J_1$ = 6.2 Hz, $J_2$ = 2.4 Hz, $J_3$ = 1.9 Hz, 1H, C1H), 6.05 (dd, $J_1$ = 6.2 Hz, $J_2$ = 1.5 Hz, 1H, C2H), 5.93 (d, $J$ = 6.0 Hz, 1H, C19H), 5.04 (d, $J$ = 1.8 Hz, **$^1J_{C\text{-}H}$ = 170.3 Hz**, 1H, C6H), 4.76 (m, 1H, C5H), 4.43 (t, $J$ = 5.7 Hz, 1H, C20H), 4.26 (t, $J$ = 5.4 Hz, 1H, C4H), 4.23 (dd, $J_1$ = 5.3 Hz, $J_2$ = 3.6 Hz, 1H, C21H), 4.16 (m, 3H, C3H, C12H), 4.02 (q, $J$ = 3.5 Hz, 1H, C22H), 3.93 (dd, $J_1$ = 3.4 Hz, $J_2$ = 1.8 Hz, 1H, C7H), 3.91 (m, 1H, C11Ha), 3.80 (dd, $J_1$ = 12.2 Hz, $J_2$ = 3.2 Hz, C23Ha), 3.78 (dd, $J_1$ = 9.4 Hz, $J_2$ = 3.4 Hz, 1H, C8H), 3.73 (m, 3H, C10H, C11Hb, C233Hb), 3.62 (t, $J$ = 9.4 Hz, 1H, C9H), 1.92 ($AcO^-$) ppm.

**$^{13}$C-NMR** (150 MHz, $d^3$-MeOD): δ = 162.3 (C13), 154.3 (C14), 153.7 (C15), 137.2 (C1), 132.2 (C2), 119.4 (C18), 114.0 (C17), 100.8 (C16), 100.5 (C6), 89.5 (C19), 86.5 (C22), 79.0 (C4), 75.6 (C20), 75.6 (C10), 73.3 (C5), 72.4 (C8), 72.2 (C21), 71.8 (C7), 68.9 (C9), 66.4 (C3), 63.3 (C23), 63.1 (C11), 43.9 (C12), 23.1 ($CO_2CH_3^-$) ppm.

**HRMS** ($ESI^-$) calc. for $C_{23}H_{32}NO_{12}$: 570.2053 $[M - H]^-$; found: 570.2052.

**α-allyl-Mannosyl-Queuosine 4**

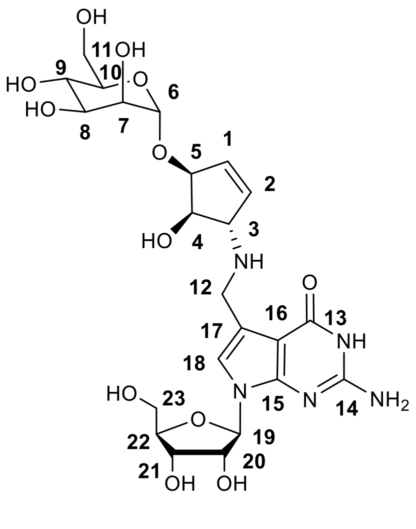

**19** (16.0 mg, 14.4 µmol) was dissolved in 10% $HNMe_2$ in THF (1 mL) and the reaction was stirred for 1 h. The solvent was evaporated in vacuo to give the free amine, which was detected by LC-MS. The amine was dissolved in MeOH (0.5 mL) and **8** (10.2 mg, 14.4 µmol) was added together with AcOH (2.00 µL). The reaction was stirred for 5 h before it was cooled to 0 °C and $NaBH_4$ (1.42 mg, 37.4 µmol) was added. The reaction was stirred at 0 °C for 30 min before $H_2O$ (0.3 mL) was added and the reaction was stirred for another 5 min. The solvent was evaporated *in vacuo* and the crude product was detected by LC-MS. The residue was dissolved in MeOH (2 mL) and NaOMe (24.1 mg, 1.00 mmol) was added. The reaction was stirred at room temperature for 18 h and then neutralized with DOWEX-$H^+$resin. The resin was filtered off and the solvent was evaporated. The residue was dissolved in DCM (1.8 mL) and TFA (0.2 mL) was added at 0 °C. The reaction was stirred at 0 °C for 5 min before the reaction was stopped by addition of a solution of $HNEt_3OAc$ in DCM (1 M, 3 mL). Saturated aqueous $NaHCO_3$-solution was added and the phases were separated. The aqueous phase containing the product was evaporated *in vacuo*. The sample was desalted using a SepPak-C18-cartridge. The sample was loaded with 100% water and the product was eluted from the column with a mixture of MeCN and water (50/50 v/v). The solvent was evaporated by lyophilization. The residue was dissolved in MeCN (1 mL) and HF · pyridine (100 µL) was added. The mixture was stirred for 1 week at room temperature before TMSOMe (1.5 mL) were added. The mixture was stirred for 1 h at room temperature before the solvent was evaporated *in vacuo*. The residue was dissolved in water, filtered and purified by reversed phase HPLC (0-10 % buffer B, 45 min) to afford the product **4** (0.24 mg, 0.42 µmol, 2.9% over 6 steps) as colorless solid.

**RP-HPLC:** $R_t$ (C18-column, 0-10% buffer B, 45 min) = 21.4 min.

**$^1$H-NMR** (800 MHz, $D_2O$): δ = 6.95 (s, 1H, C18H), 6.16 (d, $J$ = 6.3 Hz, 1H, C1H), 6.10 (dd, $J_1$ = 6.3 Hz, $J_2$ = 1.8 Hz, 1H, C2H), 5.96 (d, $J$ = 6.6 Hz, 1H, C19H), 4.96 (d, $J$ = 1.8 Hz, **$^1J_{C\text{-}H}$ = 170.0 Hz**, 1H, C6H), 4.68 (m, 1H, C5H), 4.58 (dd, $J_1$ = 5.4 Hz, $J_2$ = 3.2 Hz, 1H, C20H), 4.32 (dd, $J_1$ = 5.4 Hz, $J_2$ = 3.2 Hz, 1H, C21H), 4.15 (q, $J$ = 3.8 Hz, 1H, C22H), 4.13 (t, $J$ = 5.2 Hz, 1H, C4H), 3.97 (d, $J$ = 13.9 Hz, 1H, C12Ha), 3.94 (d, $J$ = 13.9 Hz, 1H, C12Hb), 3.90 (dd, $J_1$ = 3.4 Hz, $J_2$ = 1.8 Hz, 1H, C7H), 3.84 (m, 3H, C3H, C8H, C11Ha), 3.81 (dd, $J_1$ = 12.5 Hz, $J_2$ = 3.4 Hz, 1H, C23Ha), 3.76 (m, 3H, C10H, C11Hb, C23Hb), 3.65 (t, $J$ = 9.7 Hz, 1H, C9H) ppm.

**$^{13}$C-NMR** (HSQC, HMBC, $D_2O$): δ = 153.2 (C14/C15), 151.7 (C14/C15), 136.1 (C2), 130.8 (C1), 116.8 (C18), 115.5 (C17), 100.8 (C16), 97.9 (C6), 86.5 (C19), 84.6 (C22), 77.7 (C5), 75.7 (C4), 73.1 (C20), 72.9 (C10), 70.5 (C21), 70.4 (C8), 70.2 (C7), 66.8 (C9), 66.5 (C3), 61.5 (C23), 60.8 (C11), ppm.

**HRMS** ($ESI^+$) calc. for $C_{23}H_{34}N_5O_{12}$: 572.2198 $[M + H]^+$; found: 572.2195.

### 4. NMR-Spectra of important compounds

$^{1}$H-NMR (**7**)

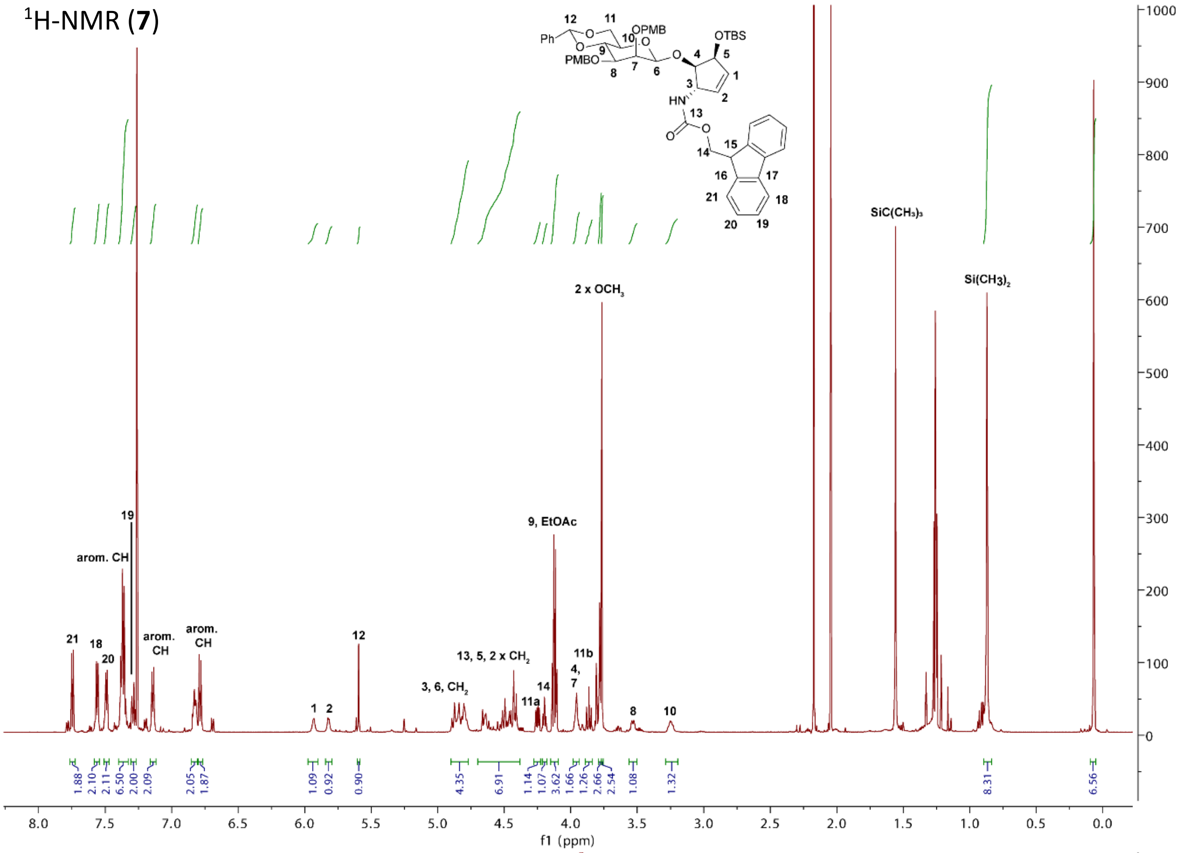

$^{13}$C-NMR (**7**)

$^{1}$H-NMR (**7**)

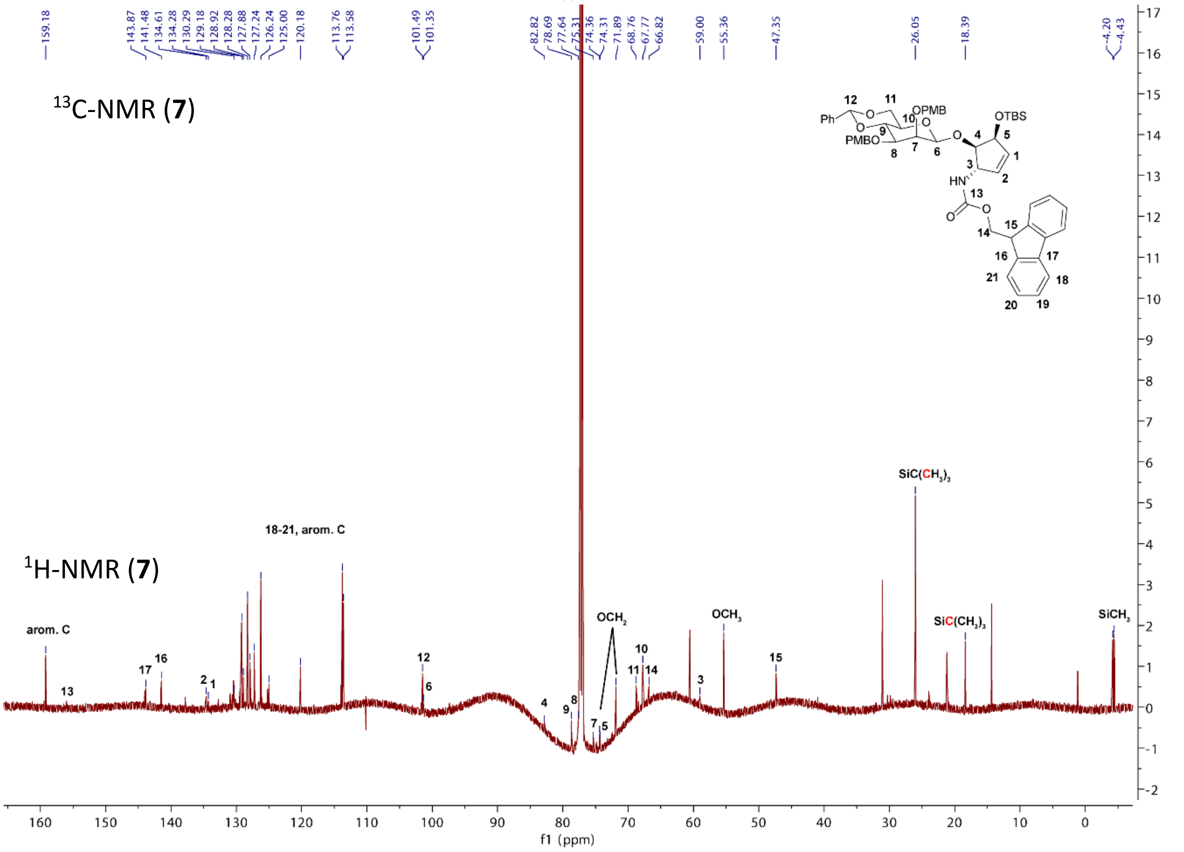

$^{1}$H-NMR (**3**)

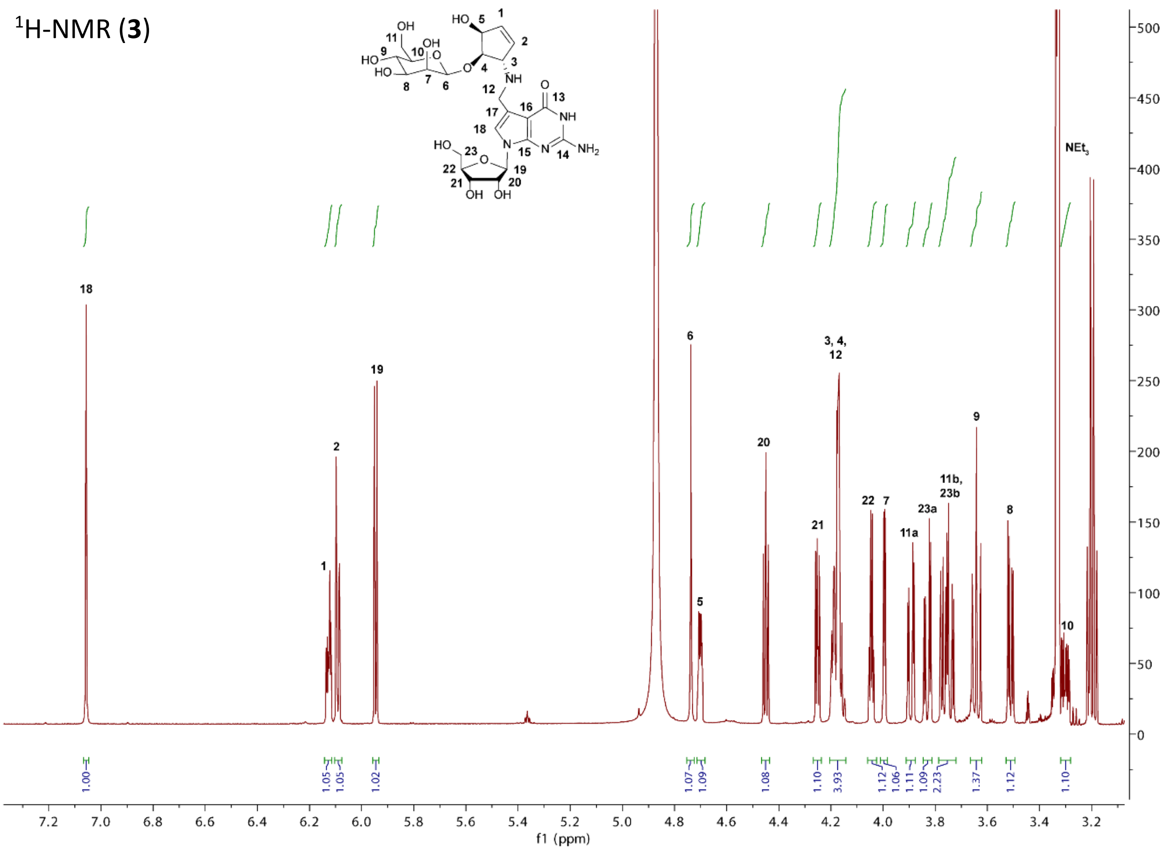

$^{13}$C-NMR (**3**)

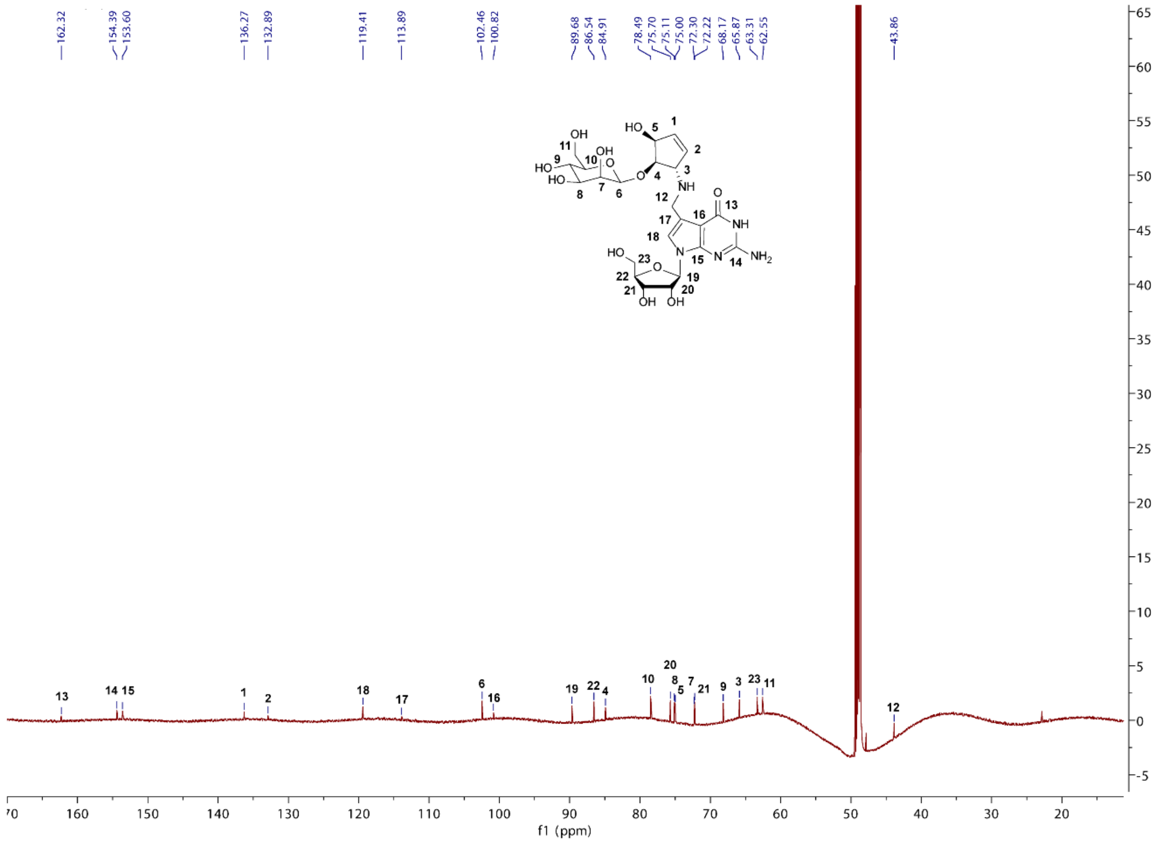

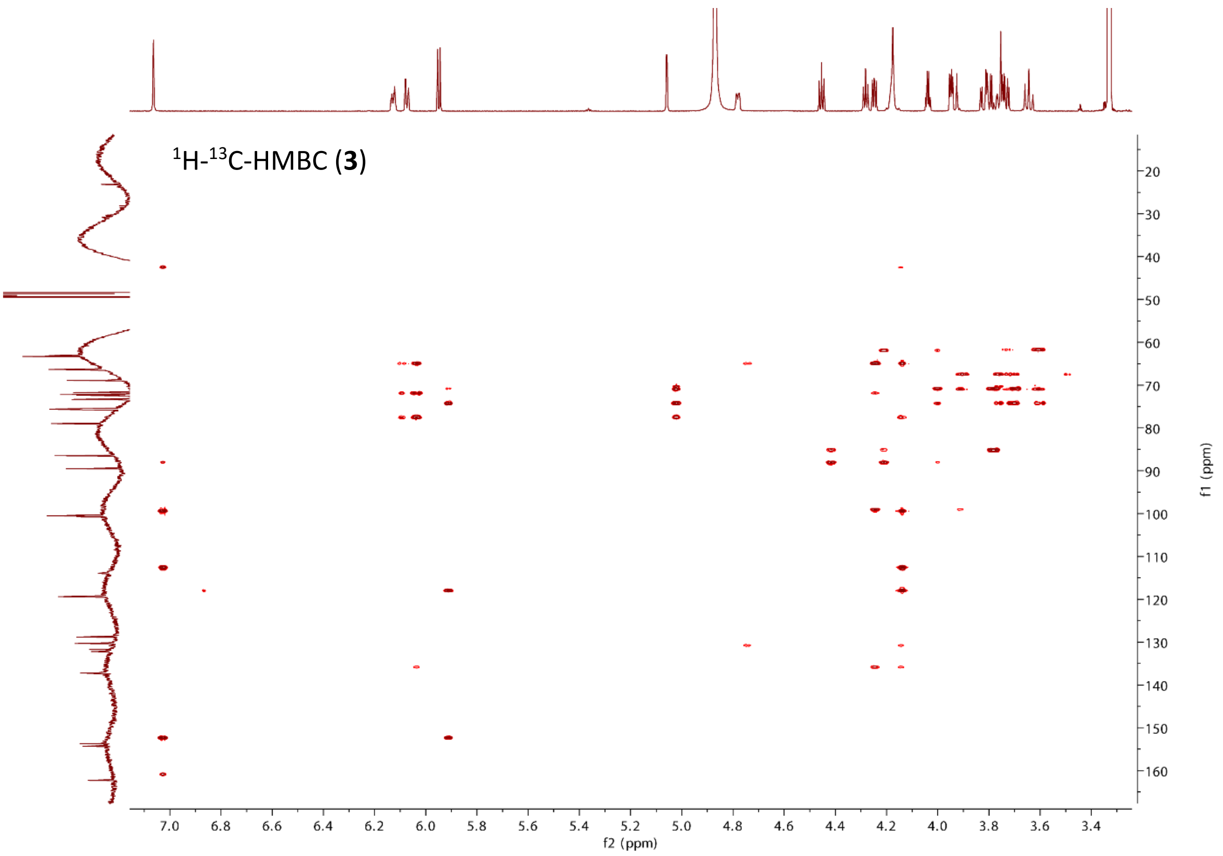
$^{1}$H-$^{13}$C-HMBC (**3**)
f2 (ppm)
f1 (ppm)

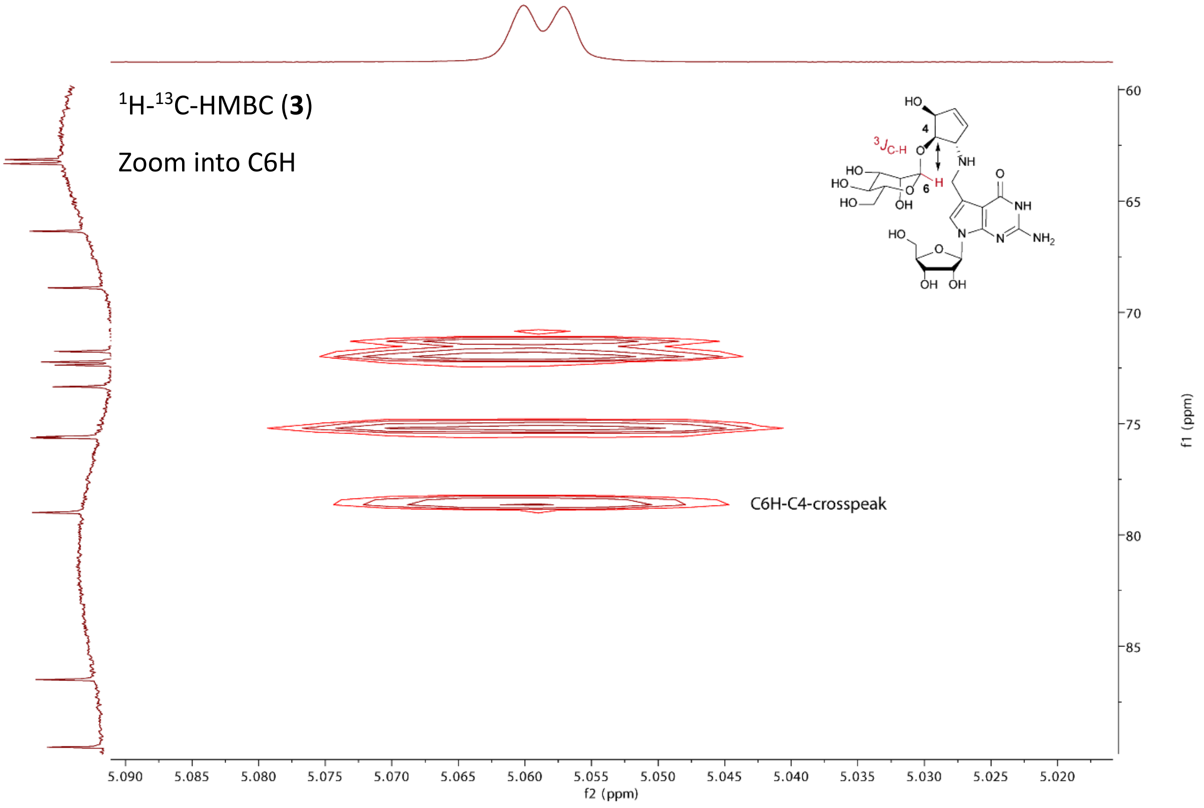
$^{1}$H-$^{13}$C-HMBC (**3**)
Zoom into C6H
C6H-C4-crosspeak
f2 (ppm)
f1 (ppm)

$^1$H-NMR (**29**)

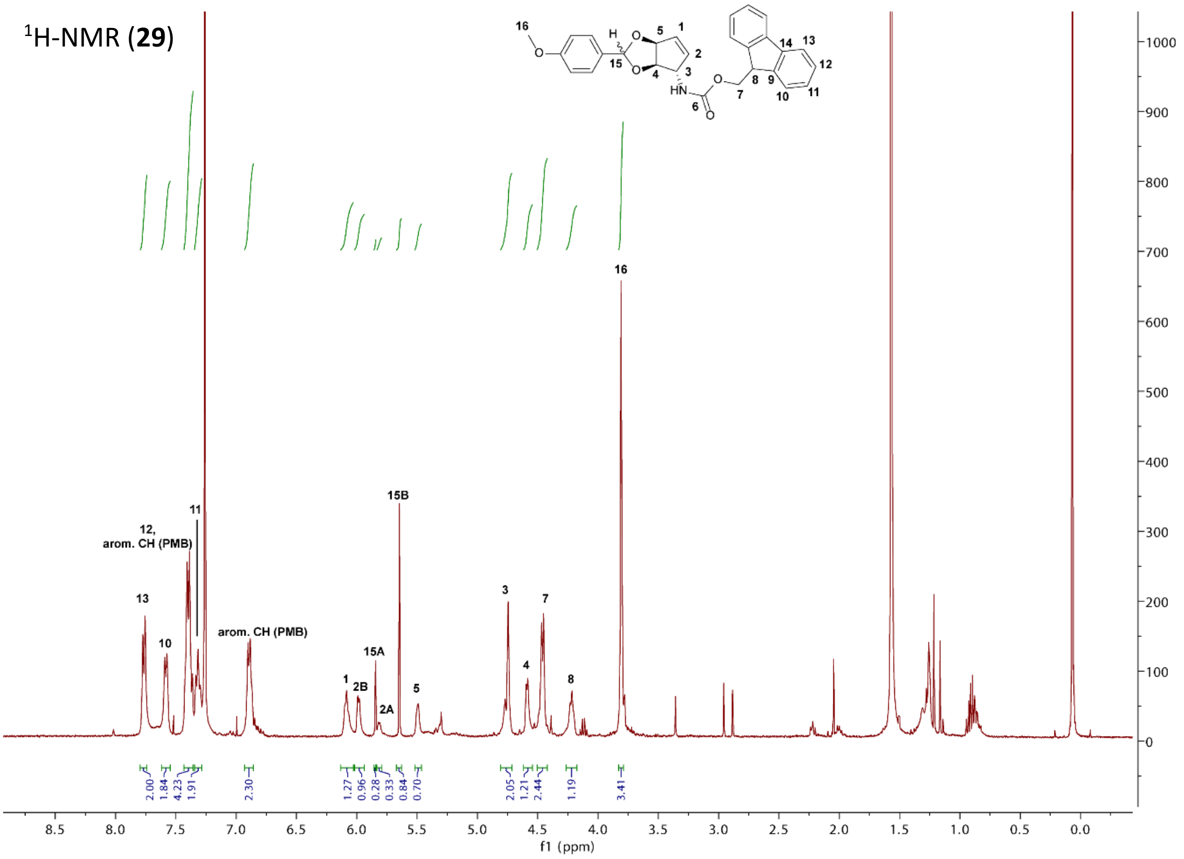

$^{13}$C-NMR (**29**)

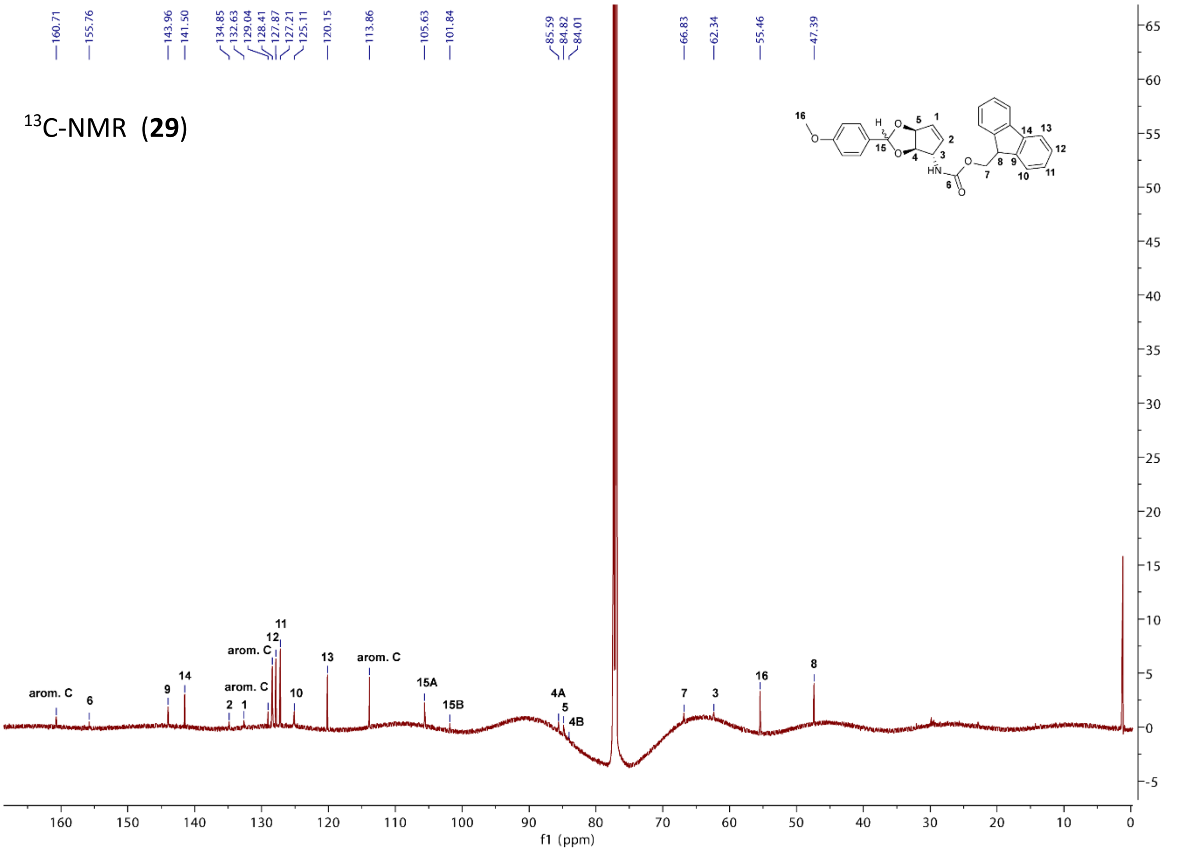

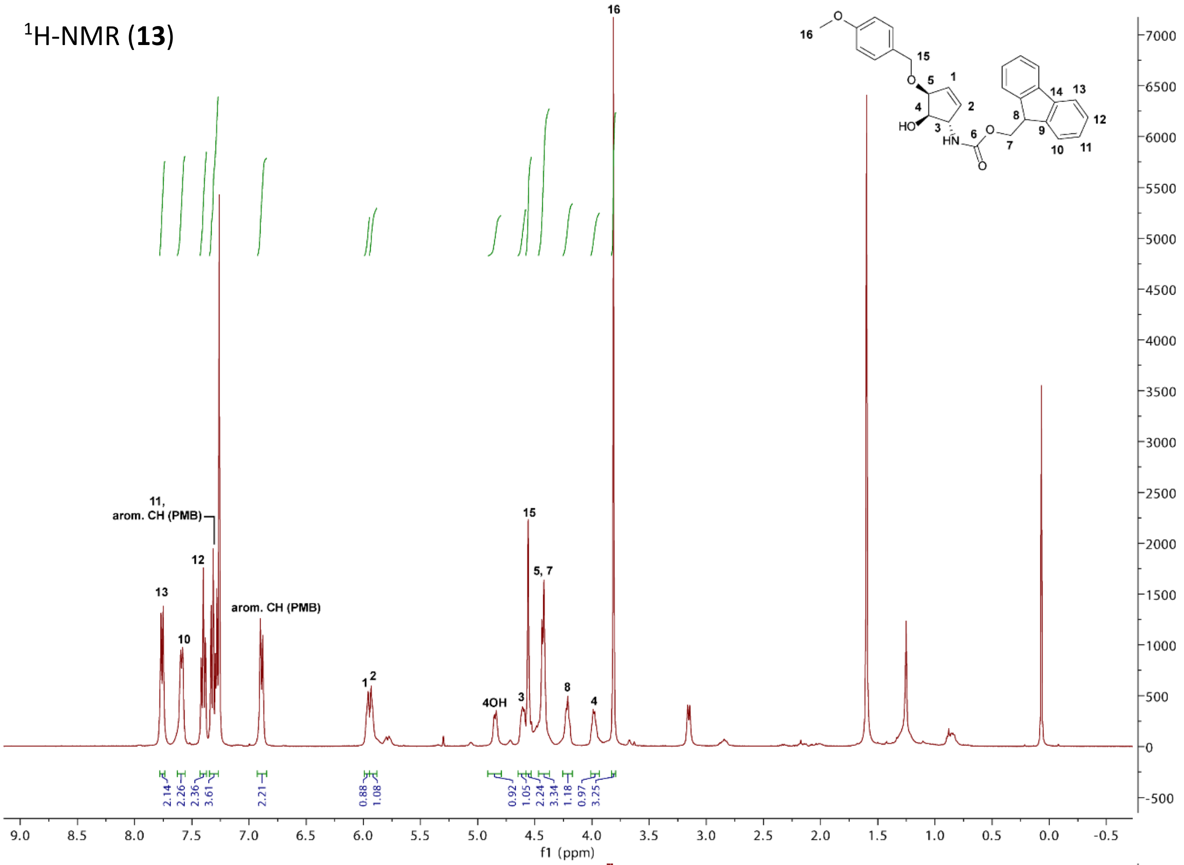

¹H-NMR (13)
16
15
5, 7
13
12
11, arom. CH (PMB)
10
arom. CH (PMB)
1 2
4OH
3
8
4
2.14
2.26
2.36
3.61
2.21
0.88
1.08
0.92
1.05
2.24
3.34
1.18
0.97
3.25
f1 (ppm)

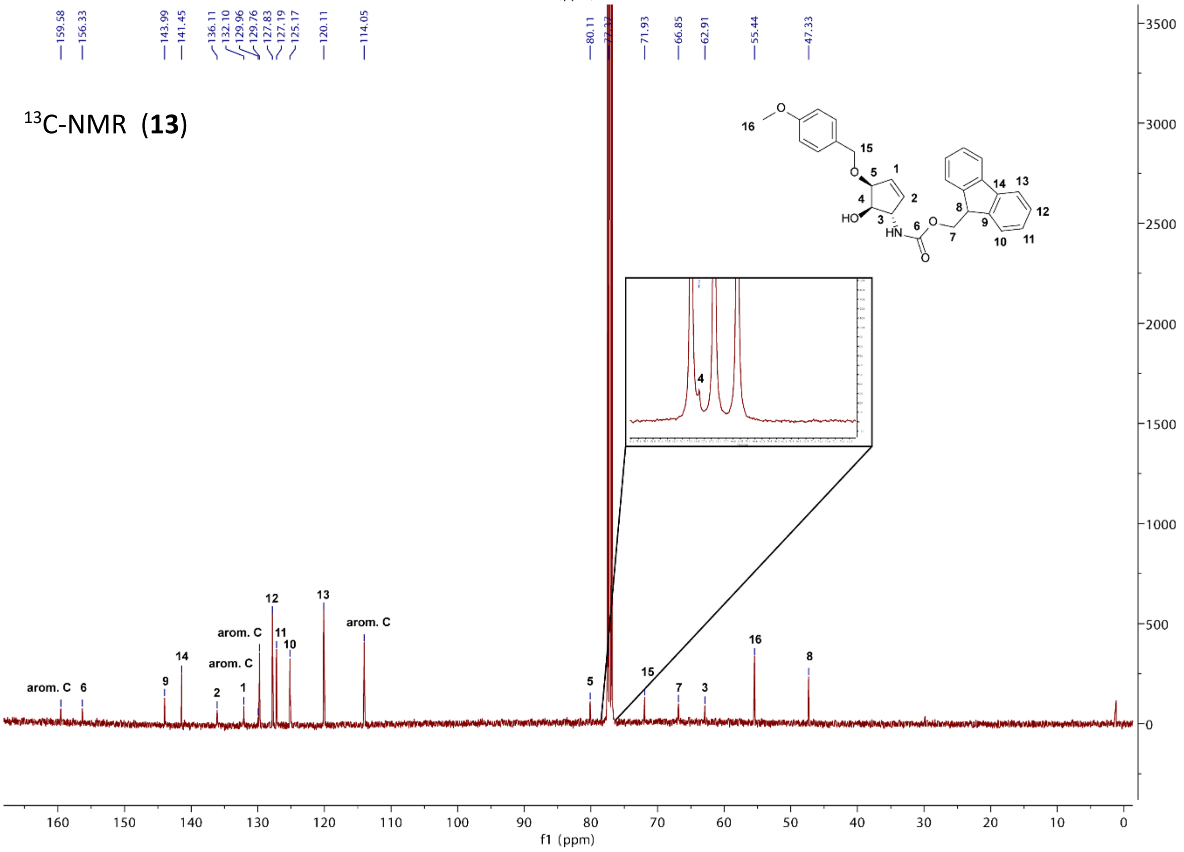

¹³C-NMR (13)
159.58
156.33
143.99
141.45
136.11
132.10
129.96
129.76
127.83
127.19
125.17
120.11
114.05
80.11
71.93
66.85
62.91
55.44
47.33
arom. C
6
9
14
2
1
arom. C
arom. C
12
11
10
13
arom. C
5
15
7
3
16
8
4
f1 (ppm)

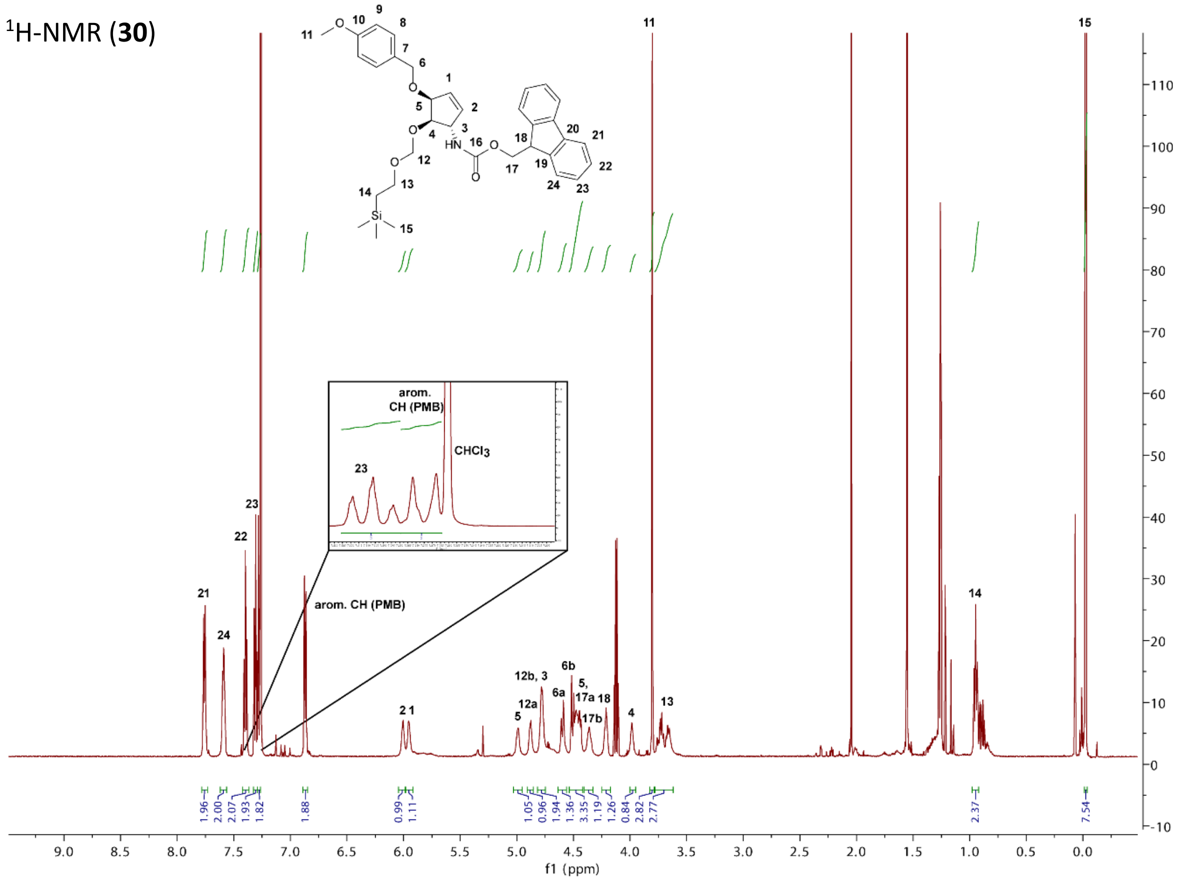

$^1$H-NMR (**30**)
arom. CH (PMB)
CHCl$_3$
12b, 3
12a
6a
6b
5, 17a
17b
f1 (ppm)

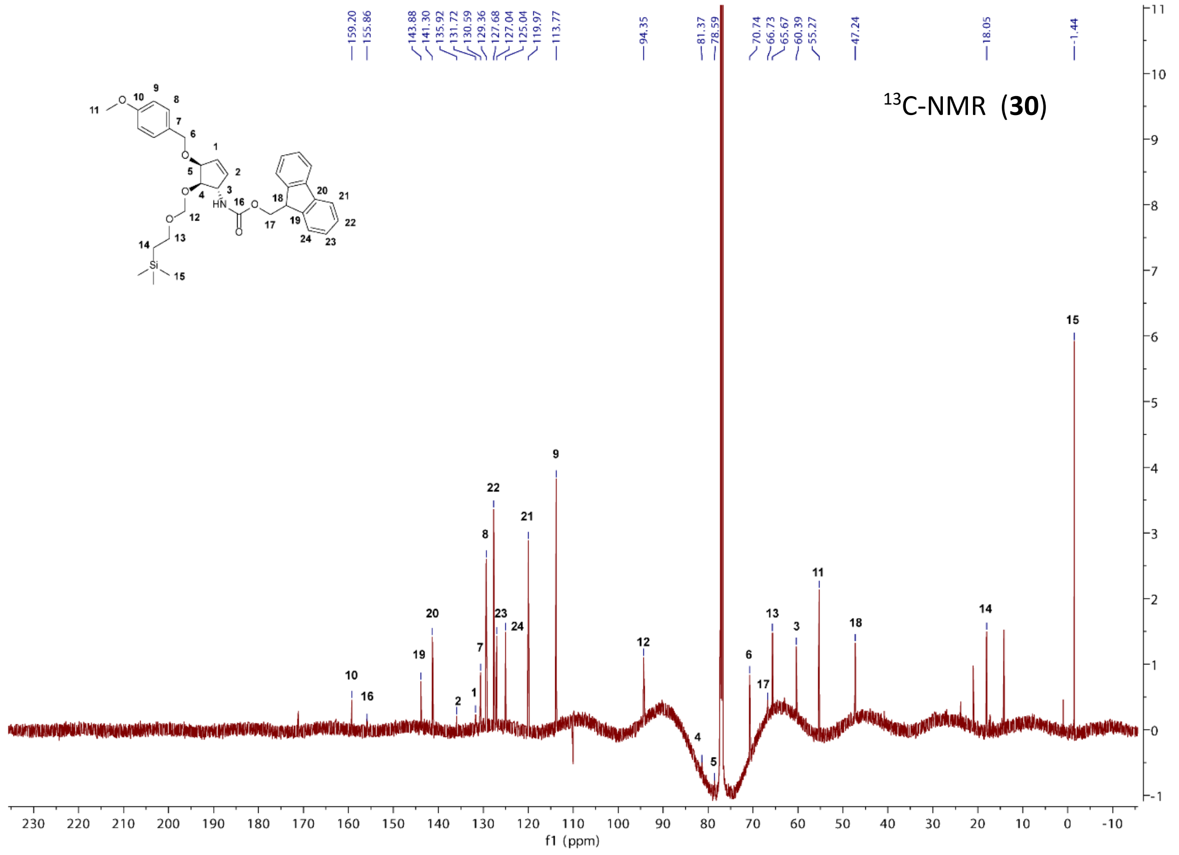

$^{13}$C-NMR (**30**)
159.20
155.86
143.88
141.30
135.92
131.72
130.59
129.36
127.68
127.04
125.04
119.97
113.77
94.35
81.37
78.59
70.74
66.73
65.67
60.39
55.27
47.24
18.05
-1.44
f1 (ppm)

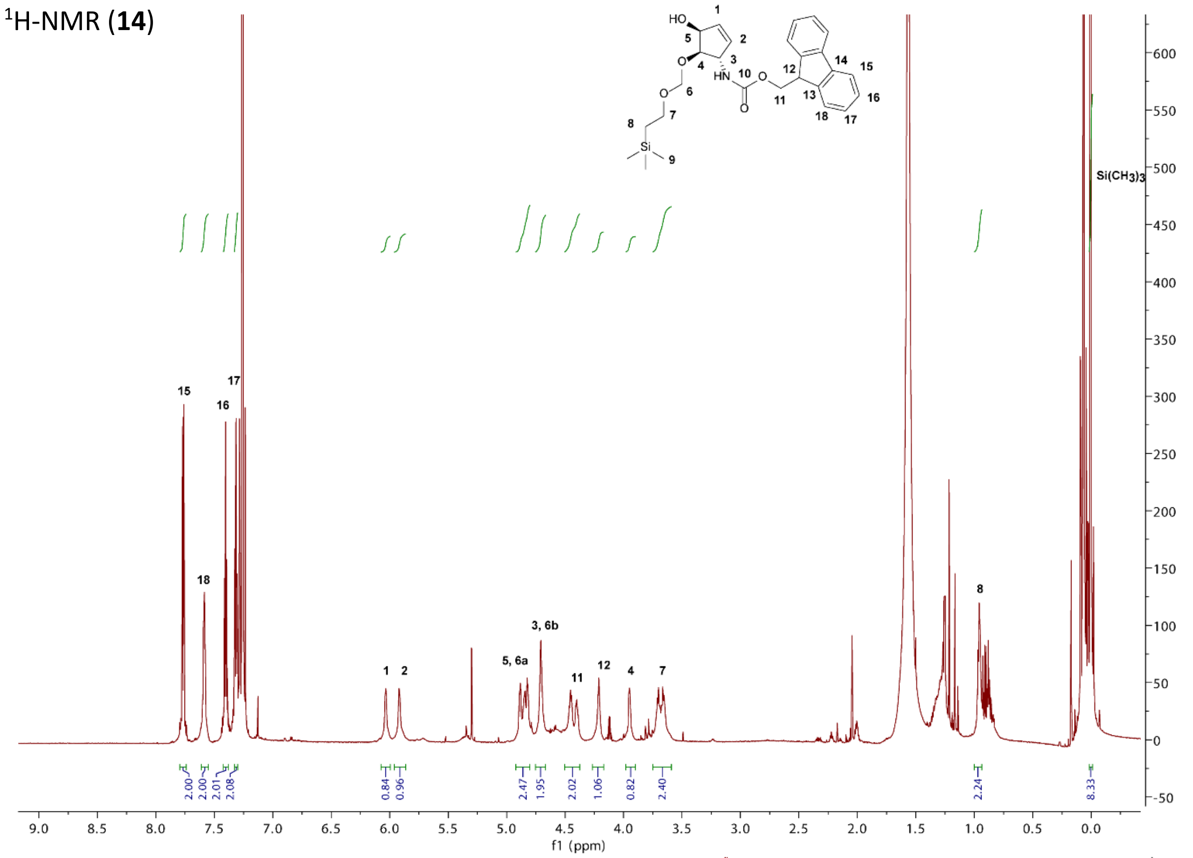
1H-NMR (14)
Si(CH3)3
15
16
17
18
1
2
5, 6a
3, 6b
11
12
4
7
8
2.00
2.00
2.01
2.08
0.84
0.96
2.47
1.95
2.02
1.06
0.82
2.40
2.24
8.33
f1 (ppm)

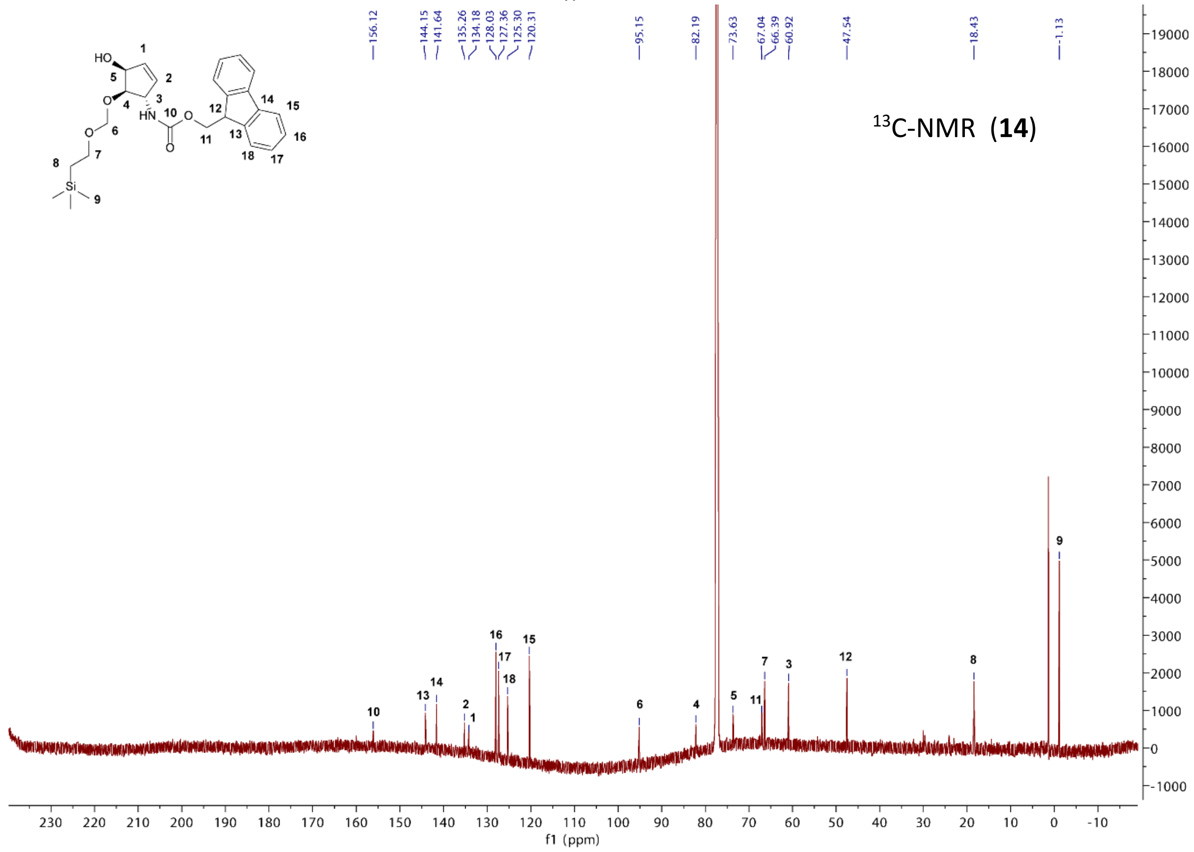
13C-NMR (14)
156.12
144.15
141.64
135.26
134.18
128.03
127.36
125.30
120.31
95.15
82.19
73.63
67.04
66.39
60.92
47.54
18.43
-1.13
f1 (ppm)

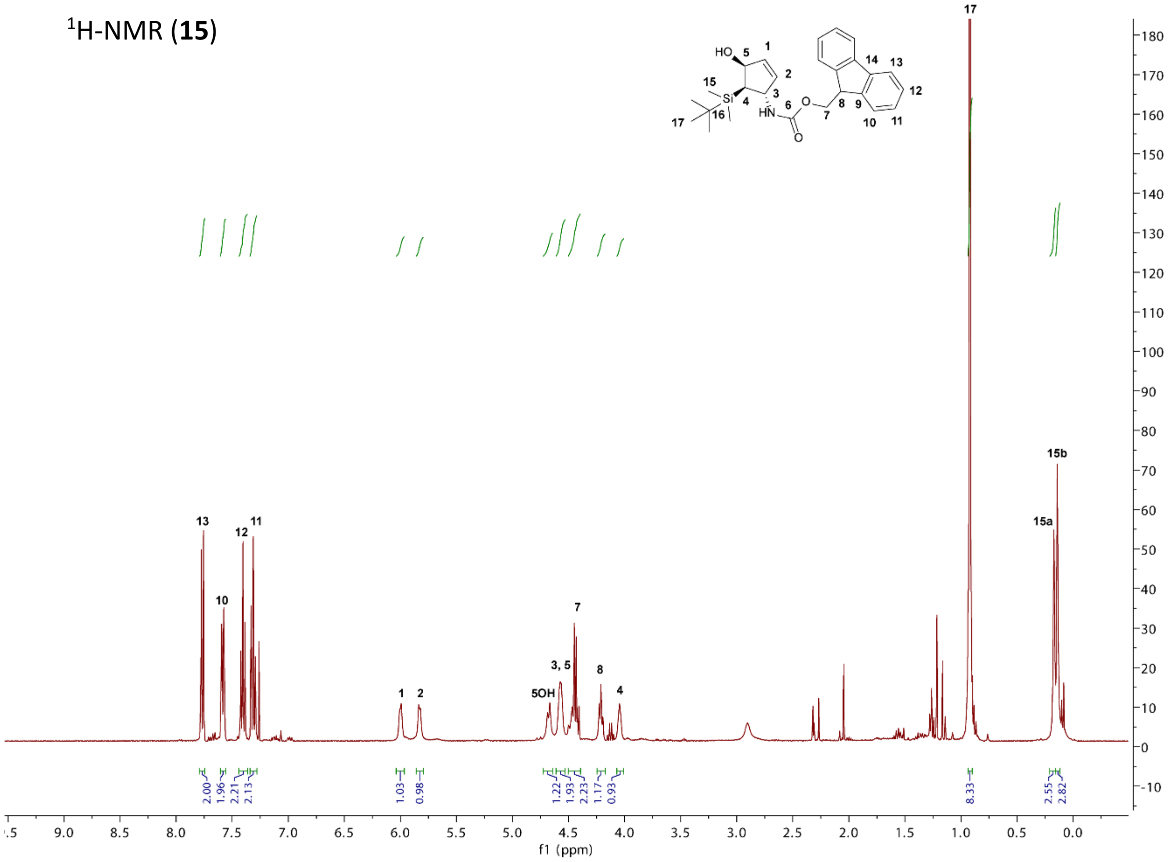

$^{1}$H-NMR (**15**)
13
10
12
11
1
2
5OH
3, 5
7
8
4
17
15a
15b
2.00
1.96
2.21
2.13
1.03
0.98
1.22
1.93
2.23
1.17
0.93
8.33
2.55
2.82
f1 (ppm)

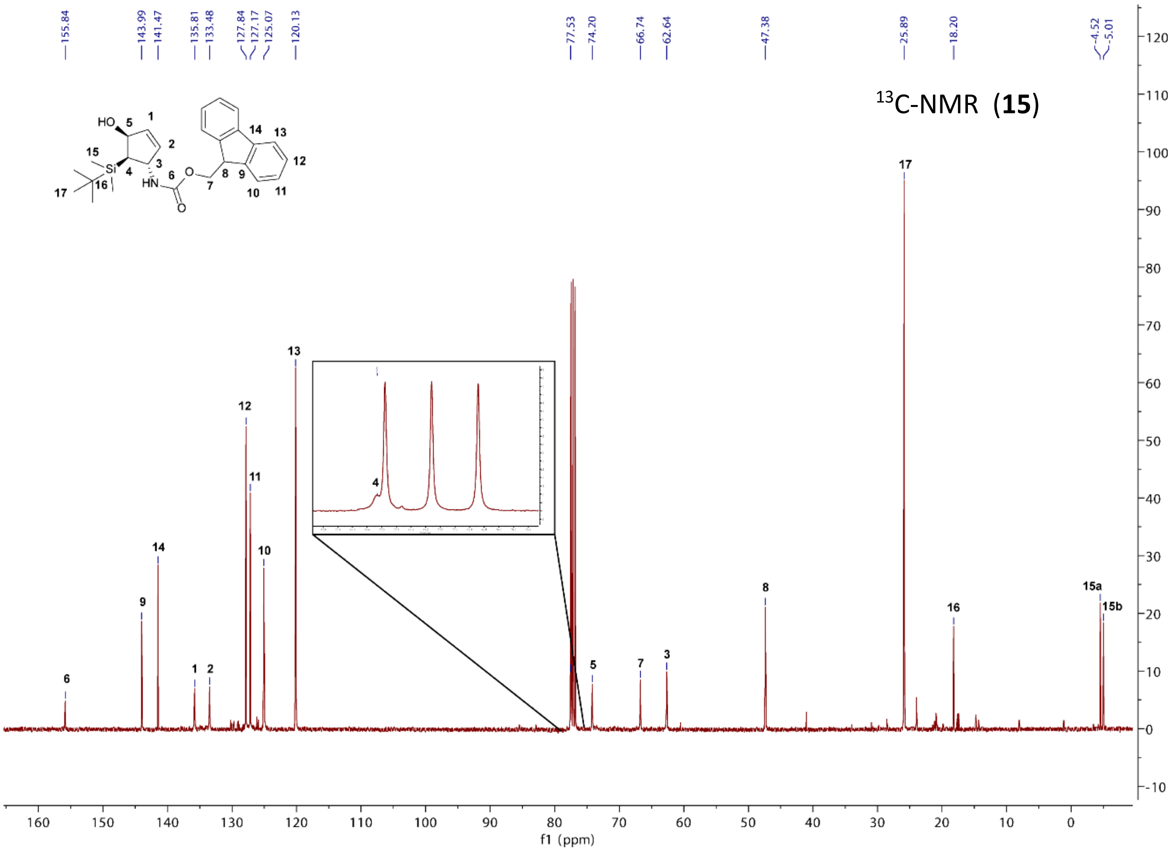

$^{13}$C-NMR (**15**)
155.84
143.99
141.47
135.81
133.48
127.84
127.17
125.07
120.13
77.53
74.20
66.74
62.64
47.38
25.89
18.20
-4.52
-5.01
6
9
14
1
2
12
11
10
13
4
5
7
3
8
17
16
15a
15b
f1 (ppm)

$^{29}$Si-NMR (**15**)

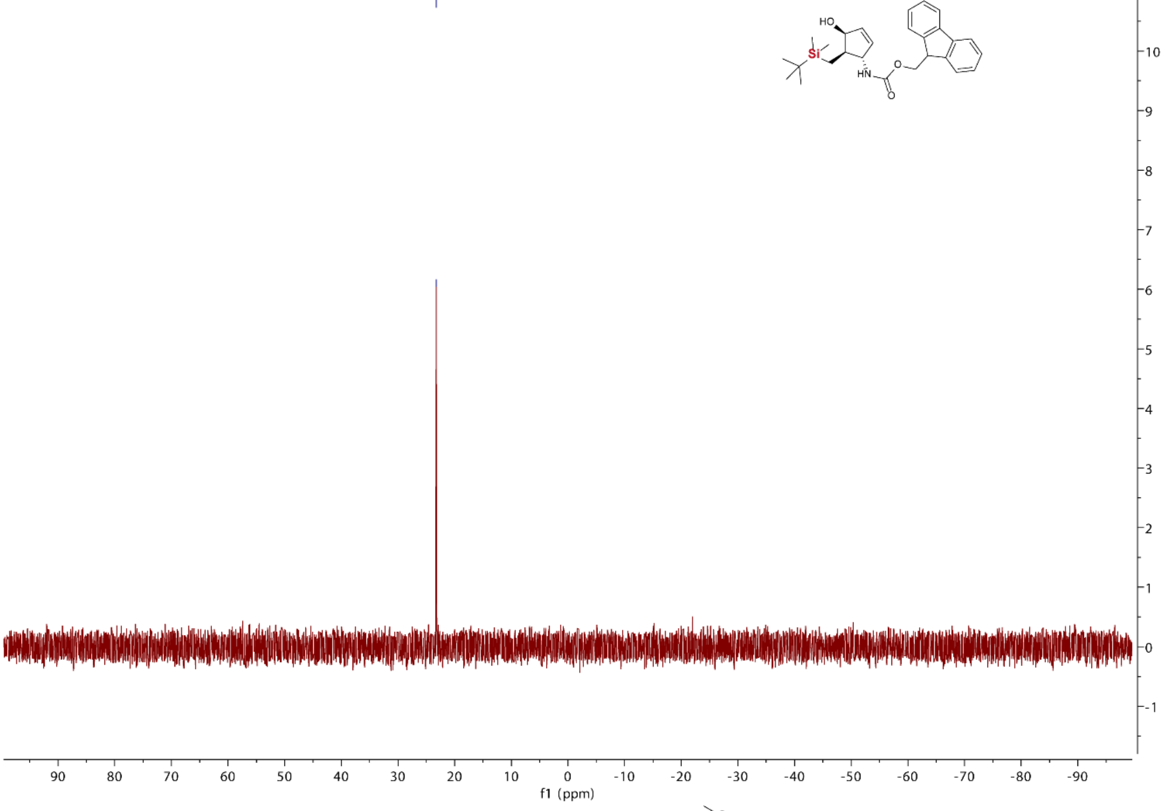

$^{1}$H-NMR (**17**)

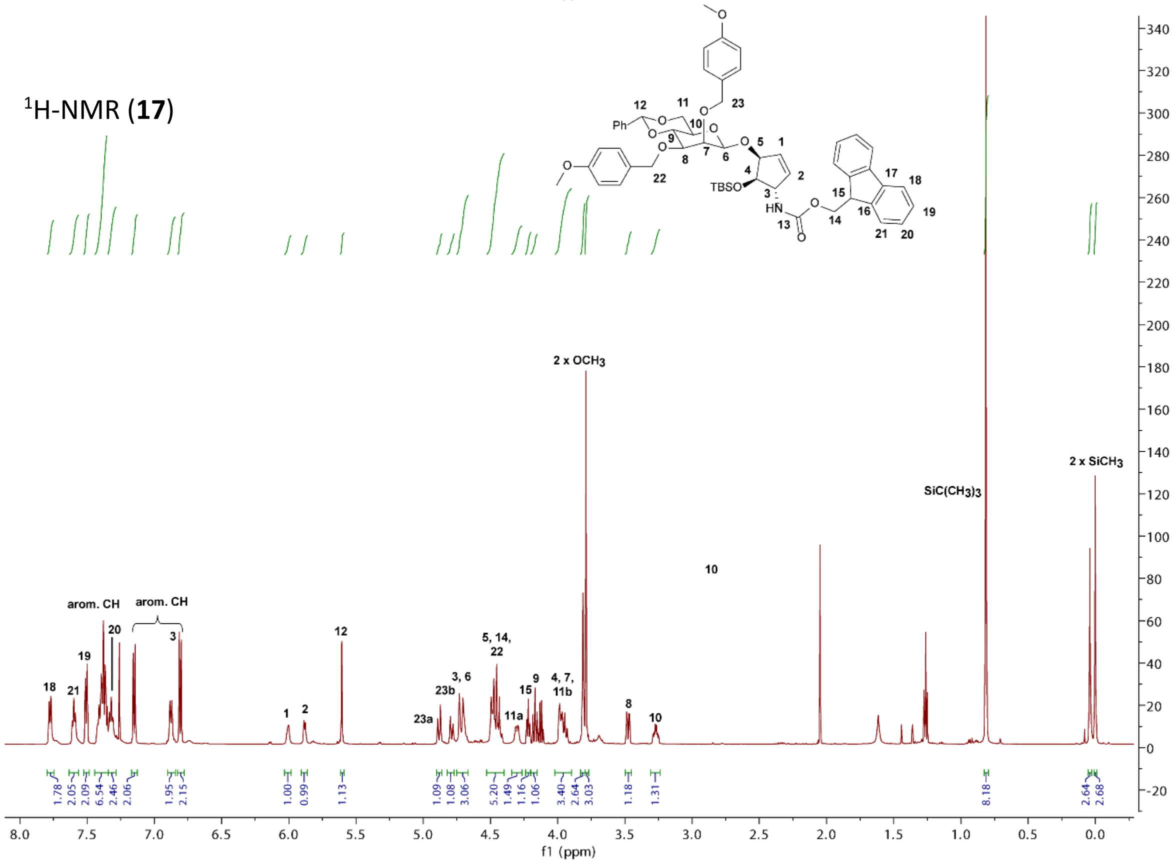

$^{13}$C-NMR (**17**)

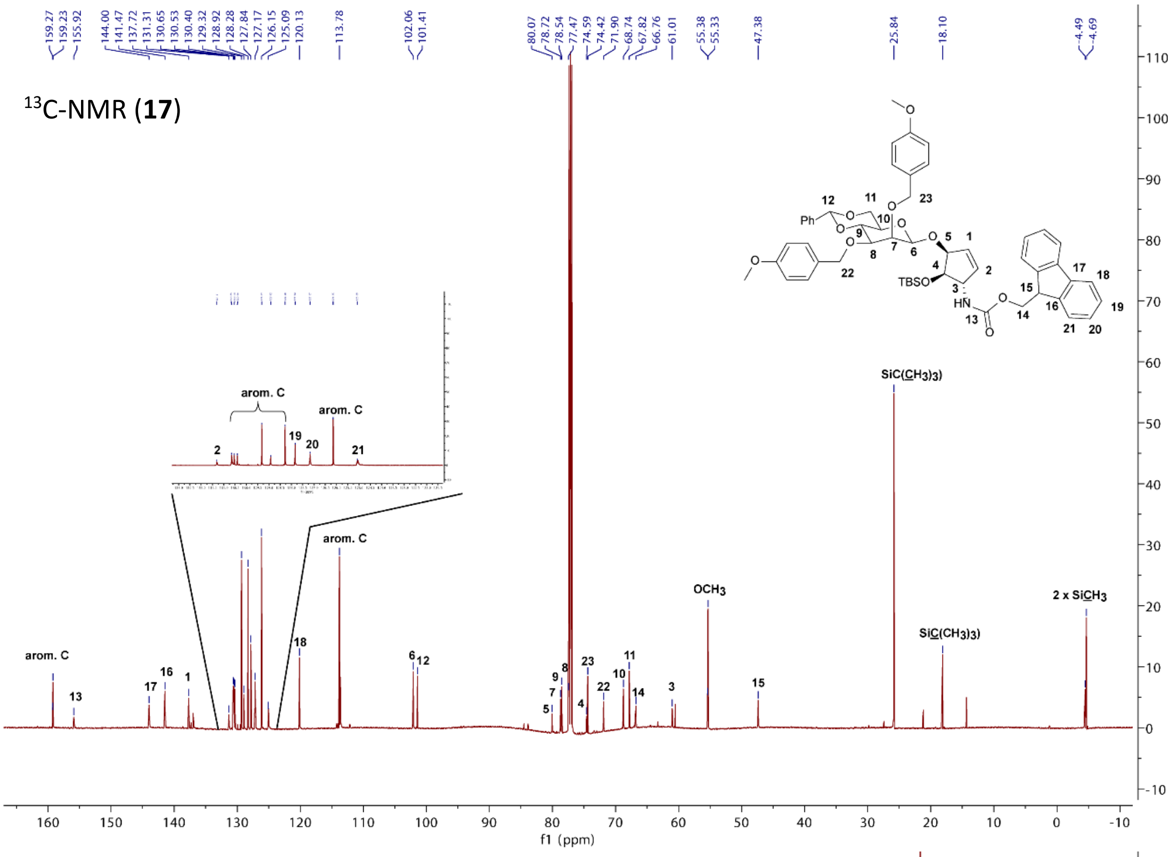

$^{1}$H-NMR (**18**)

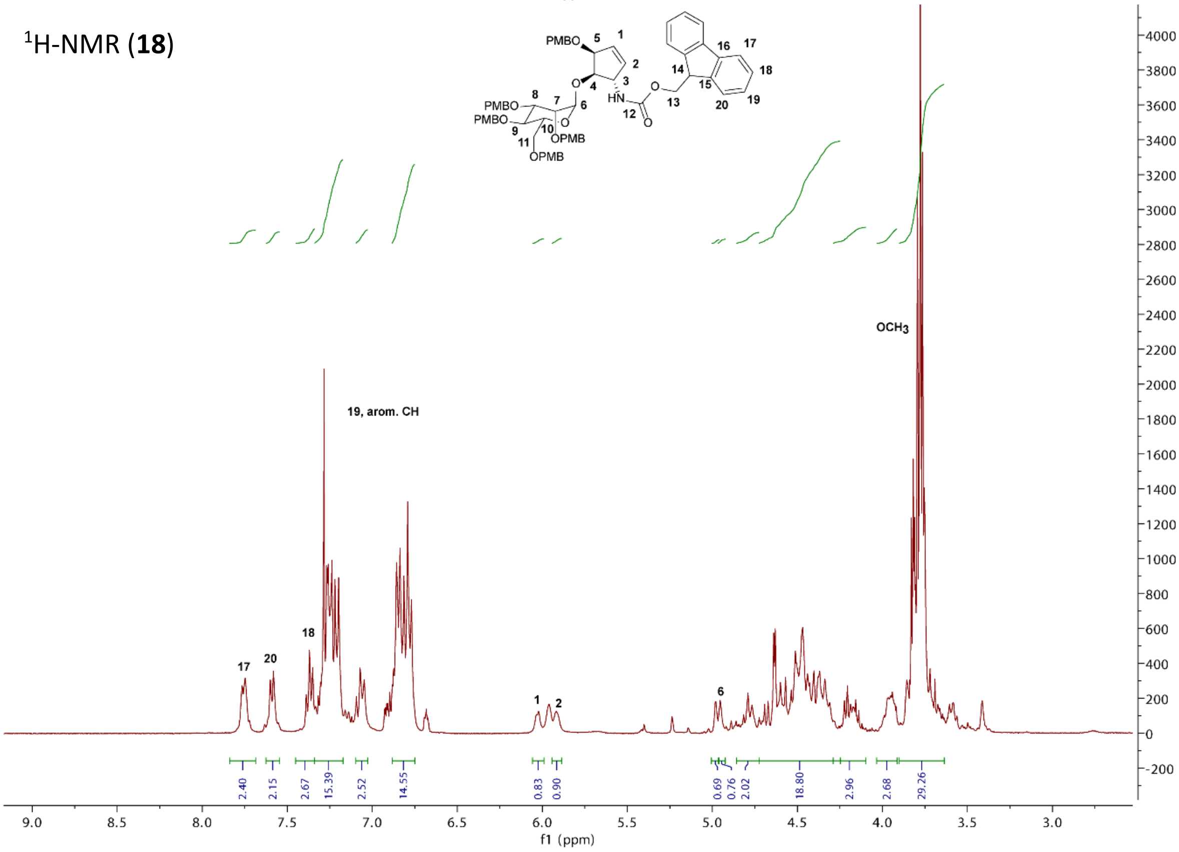

$^{13}$C-NMR (**18**)

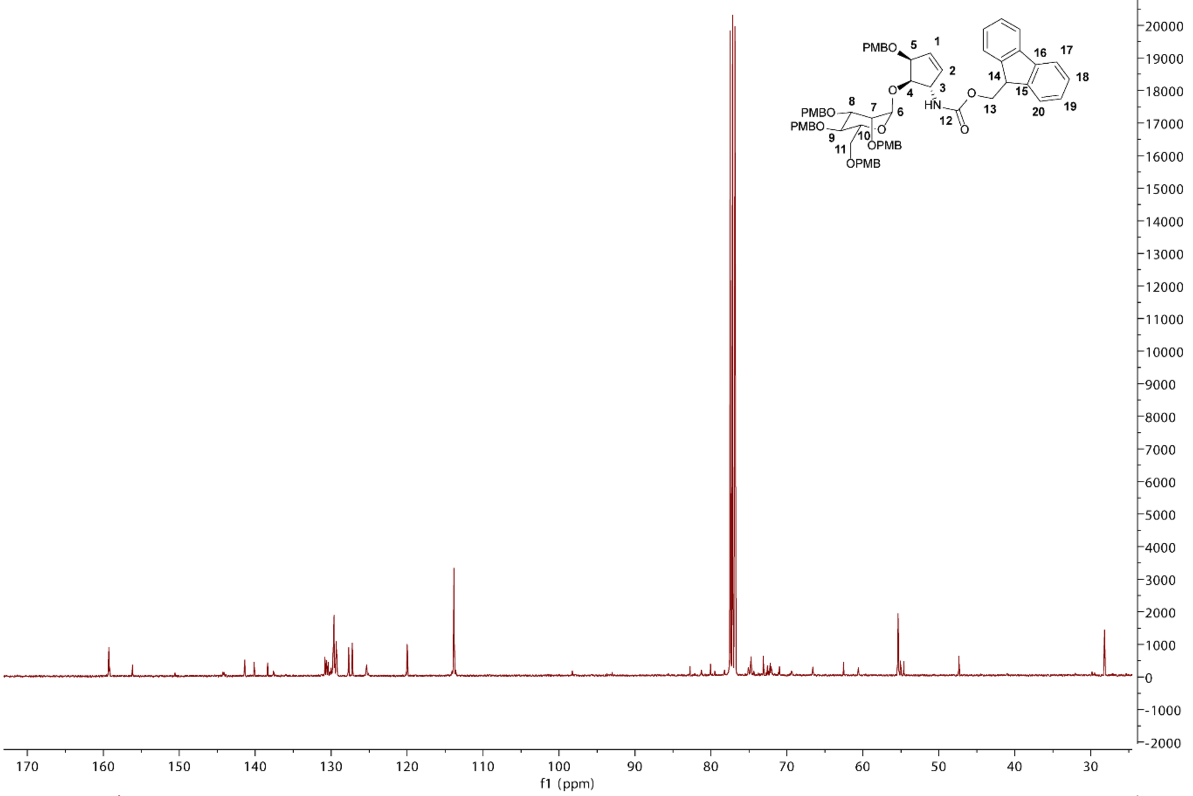

$^{1}$H-NMR (**19**)

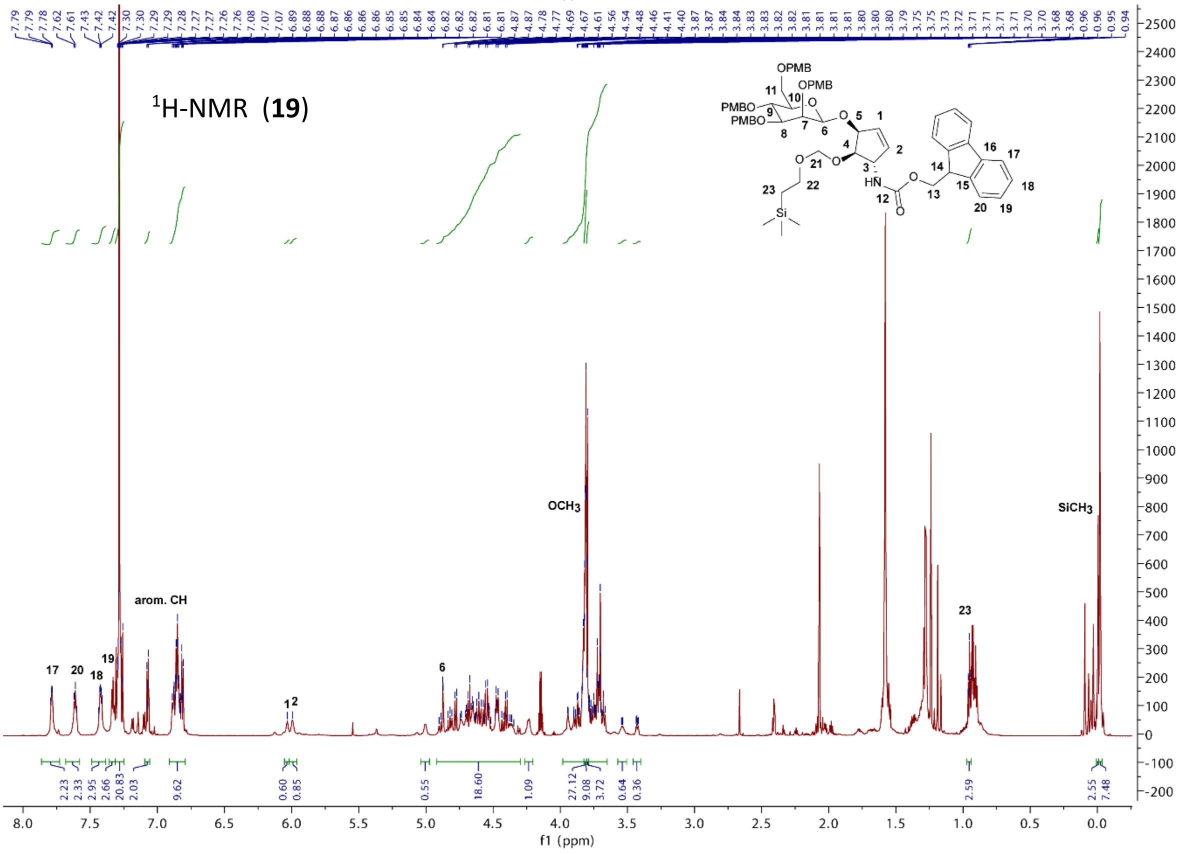

$^{13}$C-NMR (**19**)

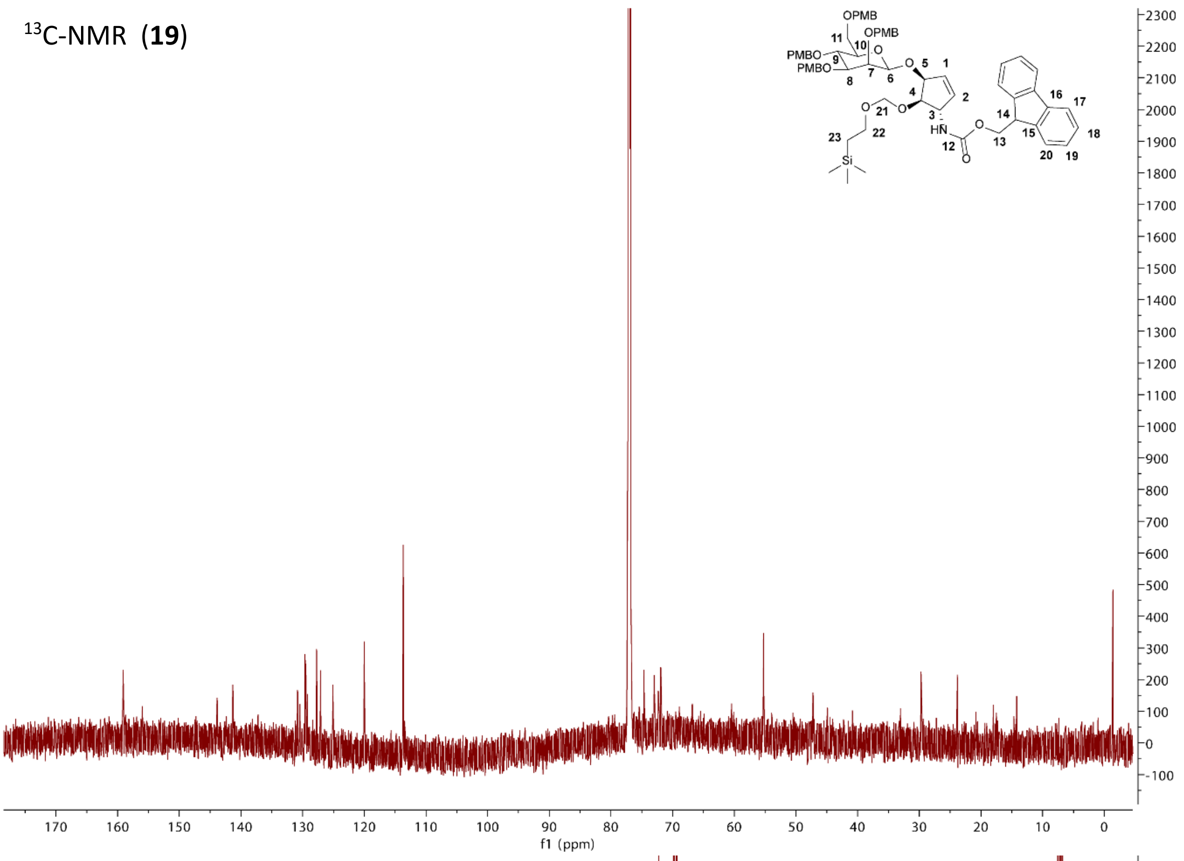

$^{1}$H-NMR (**10**)

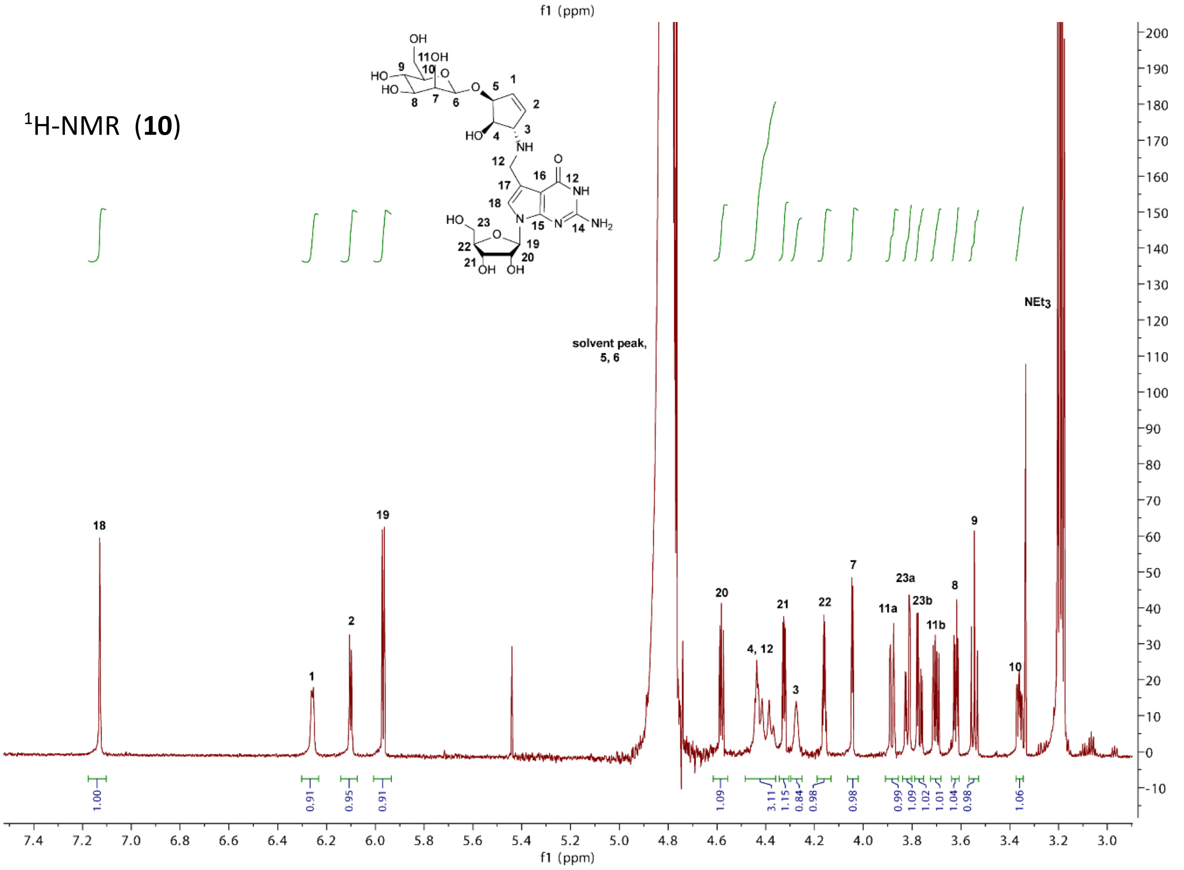

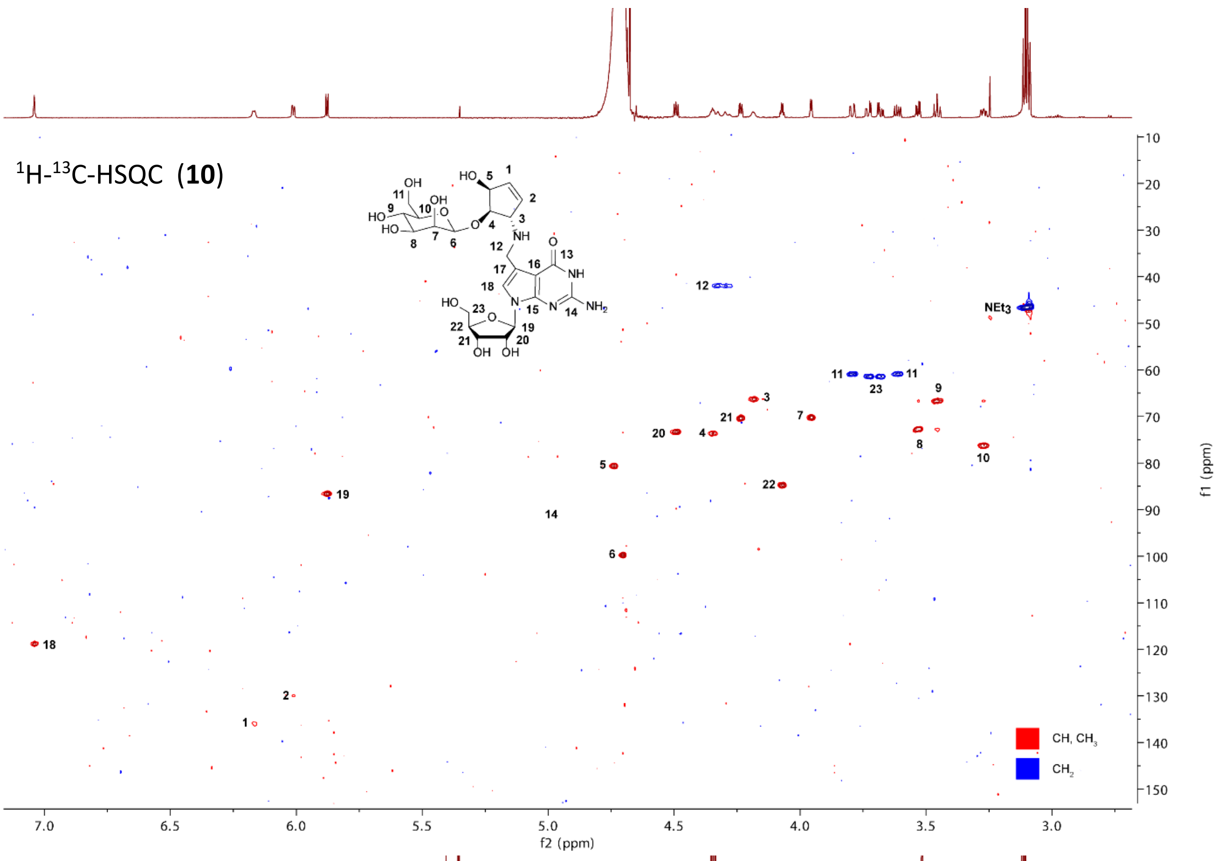

$^1$H-$^{13}$C-HSQC (**10**)
NEt3
CH, CH3
CH2
f2 (ppm)
f1 (ppm)

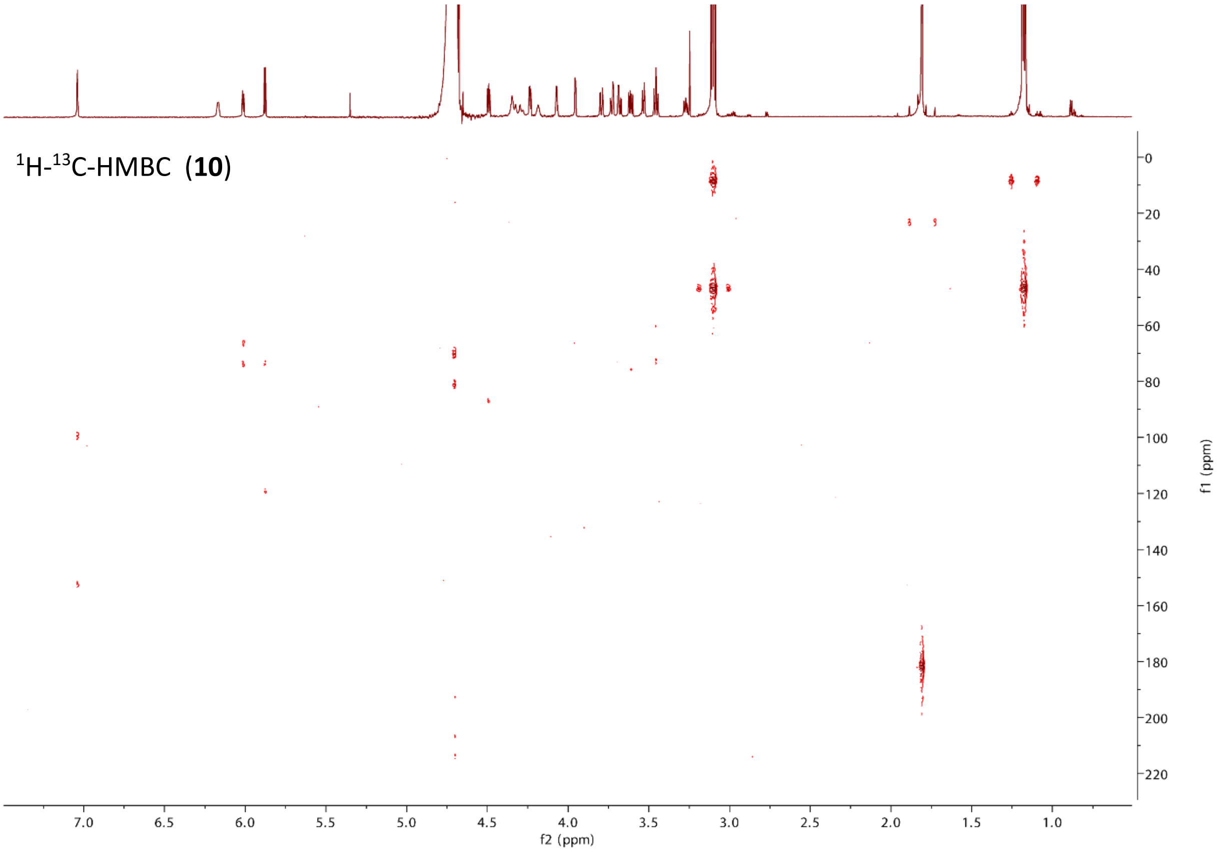

$^1$H-$^{13}$C-HMBC (**10**)
f2 (ppm)
f1 (ppm)

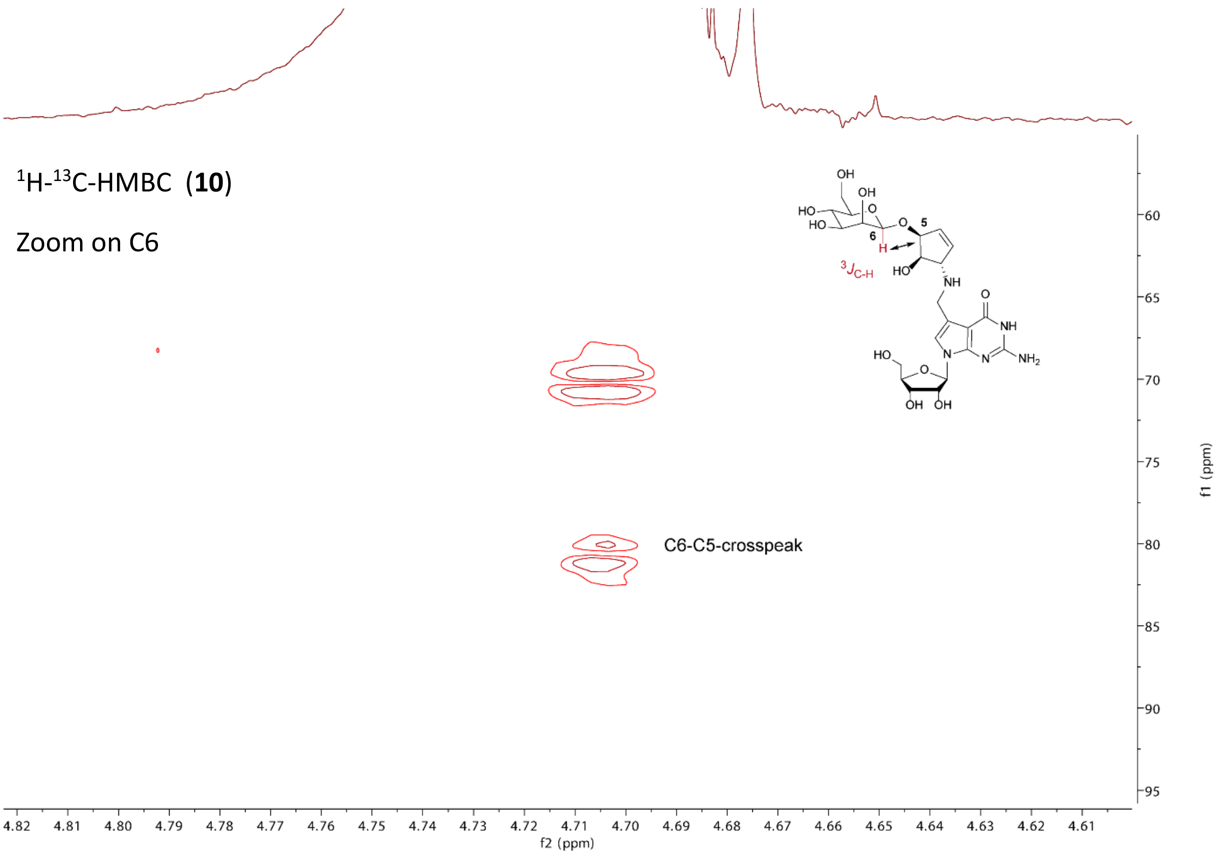

$^{1}$H-$^{13}$C-HMBC (**10**)
Zoom on C6
$^{3}J_{C-H}$
C6-C5-crosspeak
f2 (ppm)
f1 (ppm)

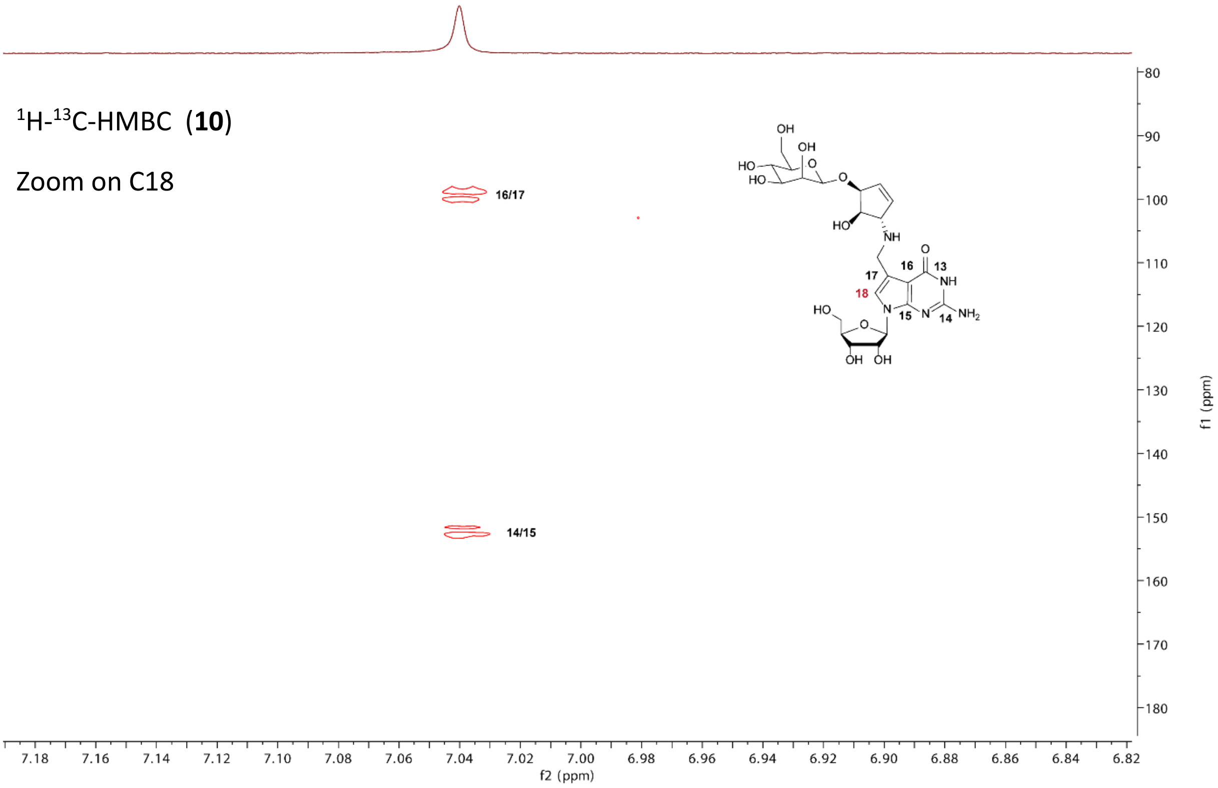

$^{1}$H-$^{13}$C-HMBC (**10**)
Zoom on C18
16/17
14/15
f2 (ppm)
f1 (ppm)

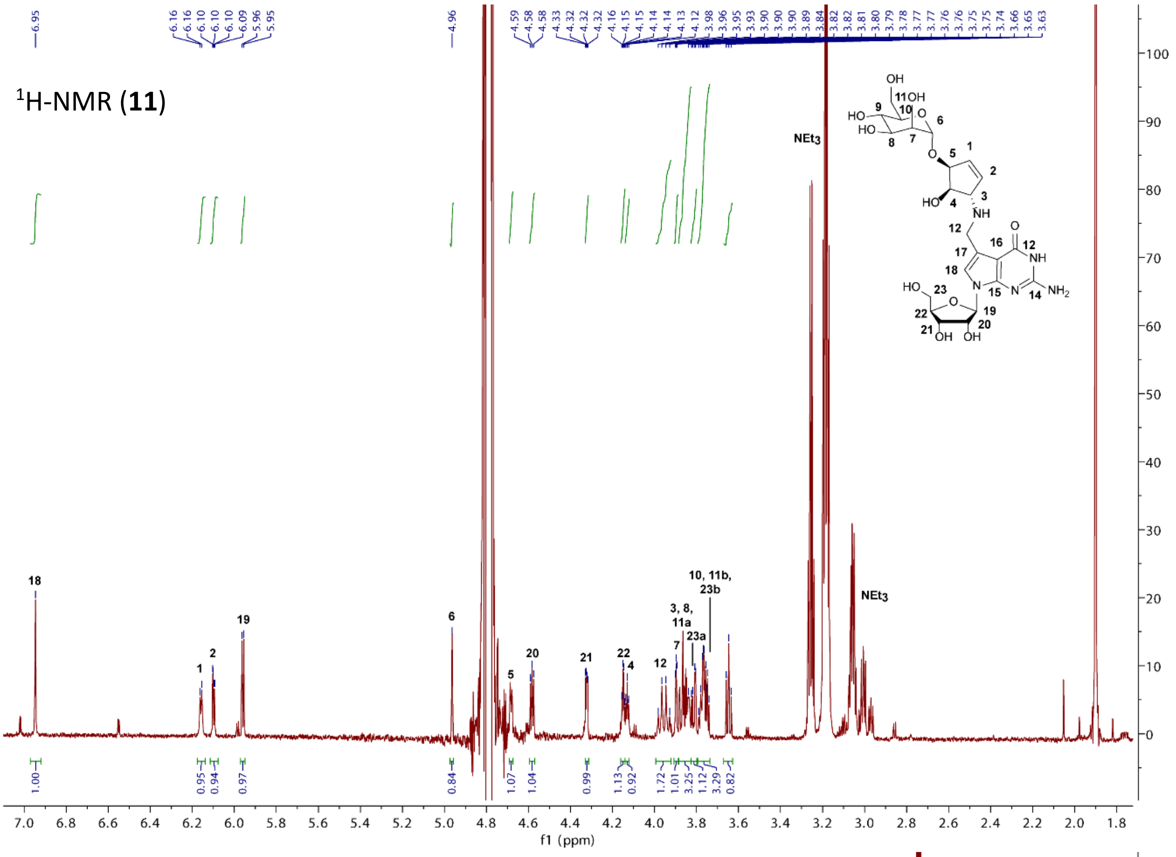

$^{1}$H-NMR (**11**)
NEt3
f1 (ppm)

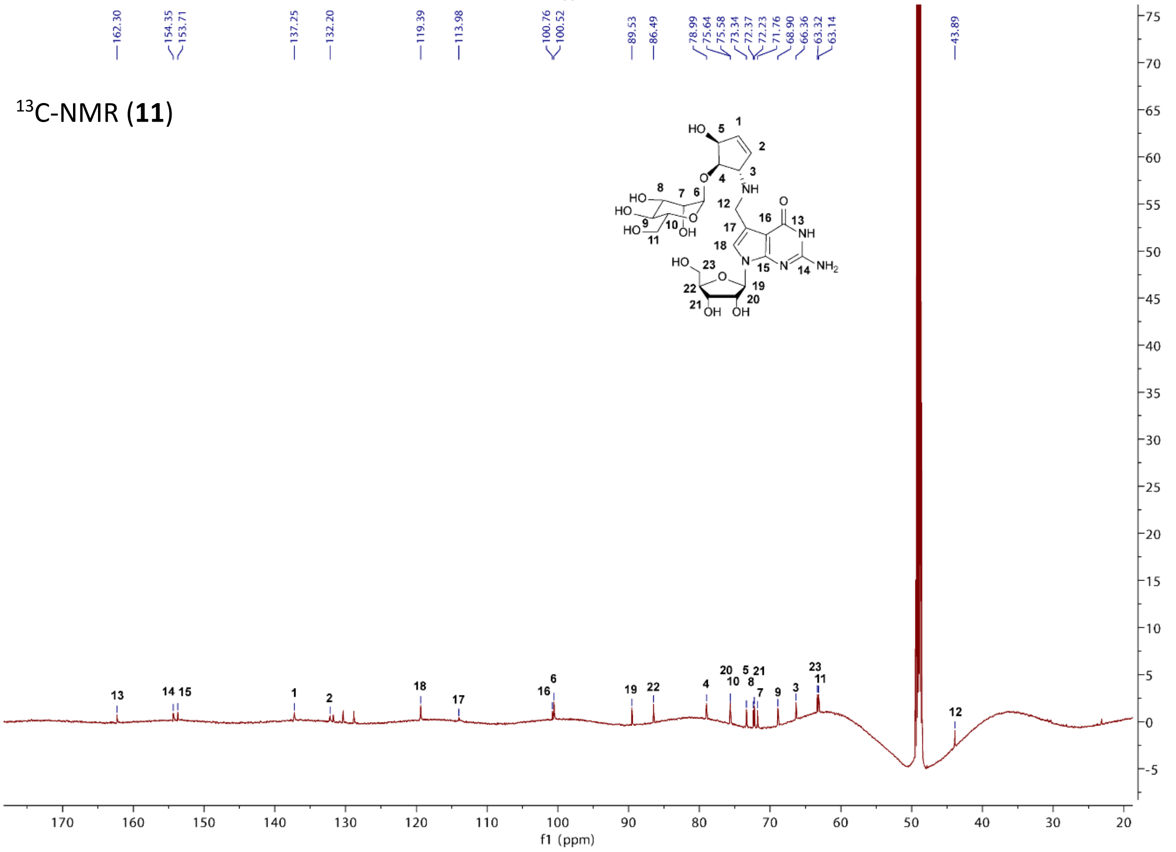

$^{13}$C-NMR (**11**)
f1 (ppm)

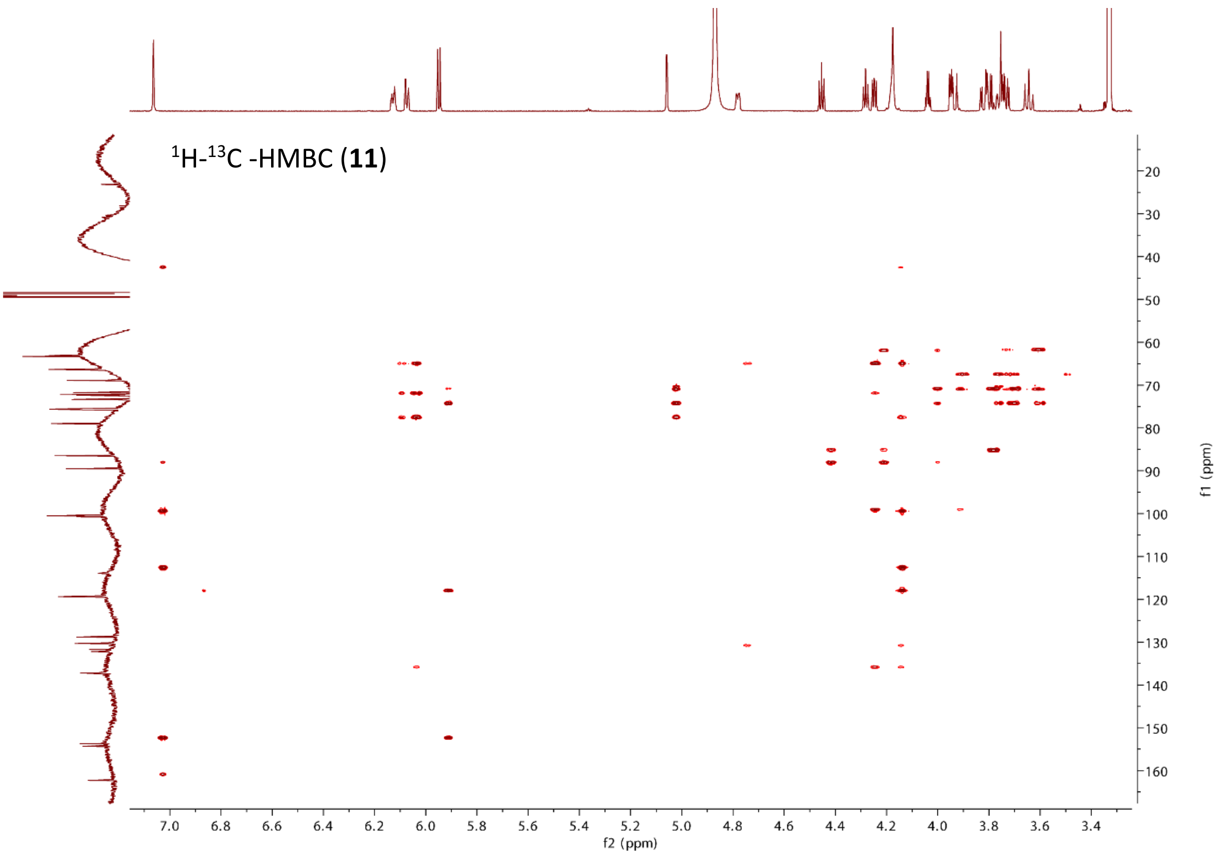

$^{1}$H-$^{13}$C -HMBC (**11**)
f2 (ppm)
f1 (ppm)

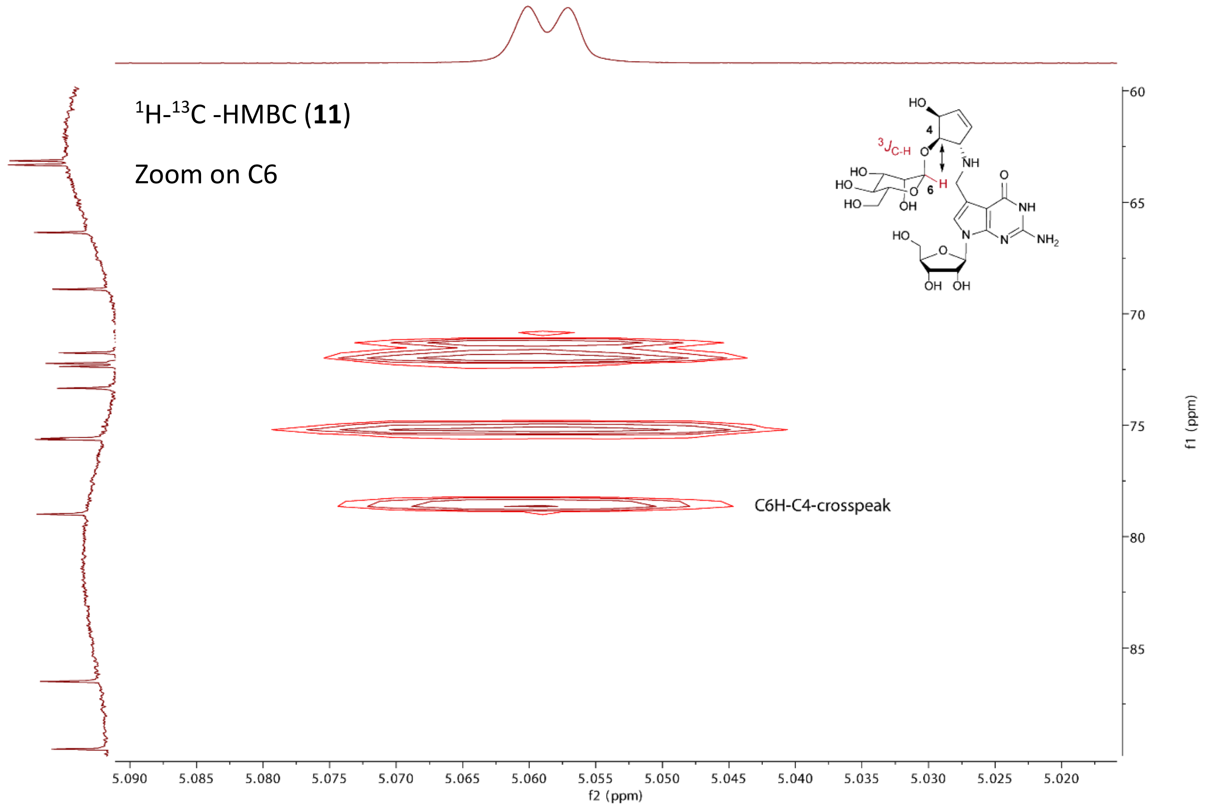

$^{1}$H-$^{13}$C -HMBC (**11**)
Zoom on C6
C6H-C4-crosspeak
f2 (ppm)
f1 (ppm)

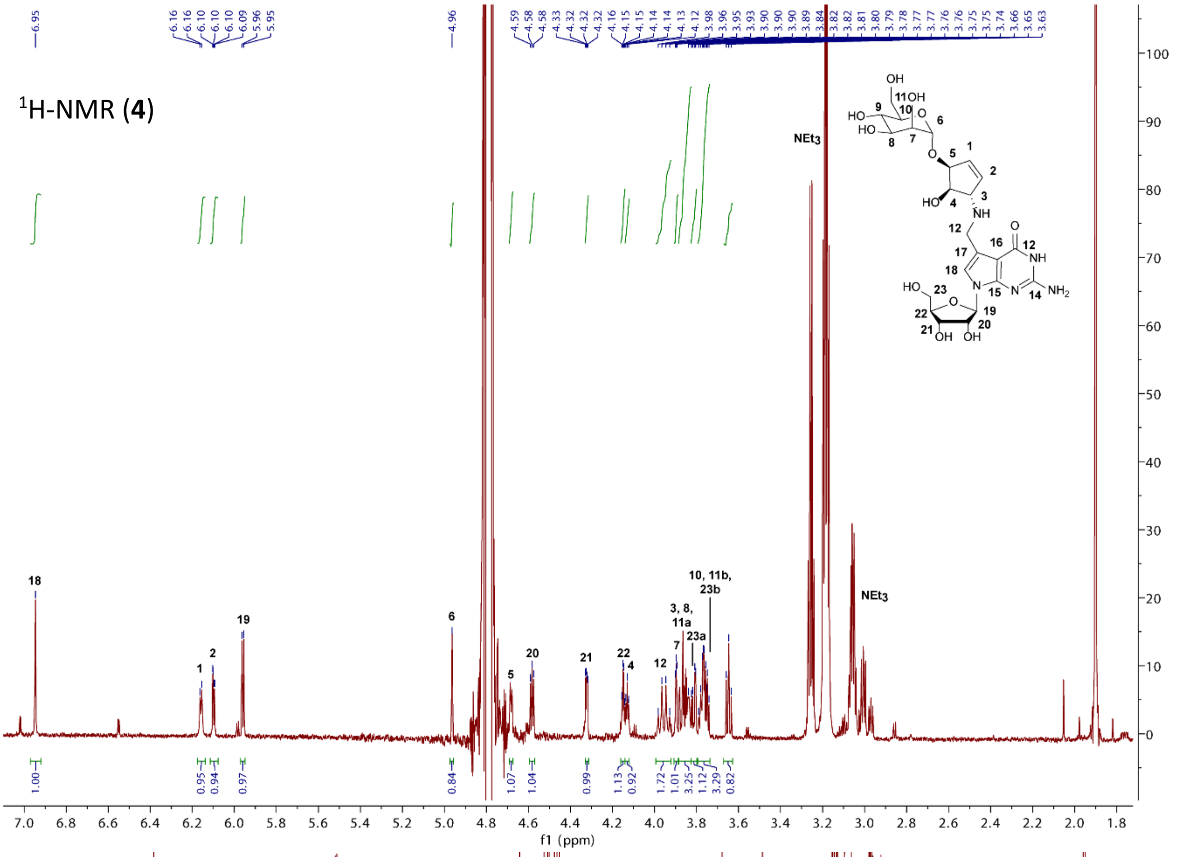

$^{1}$H-NMR (**4**)
NEt3
f1 (ppm)

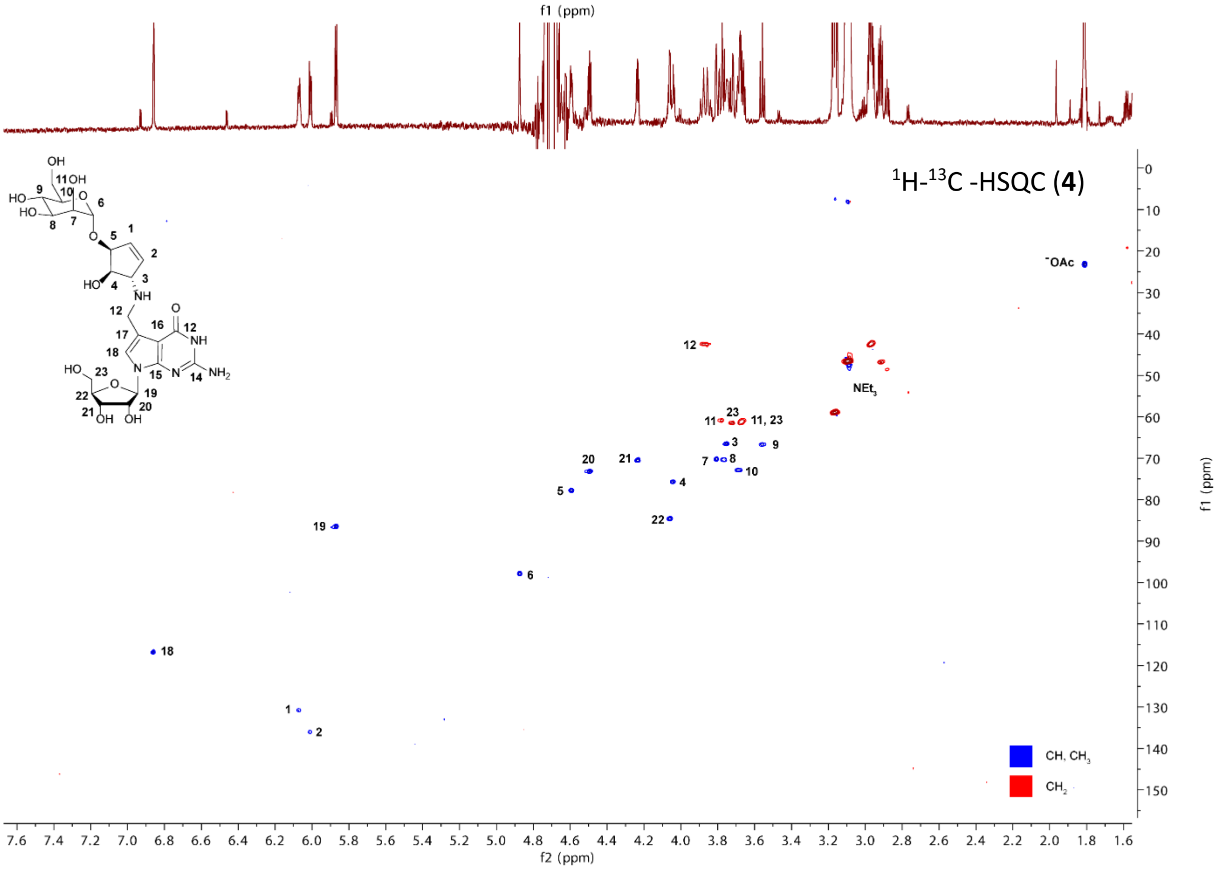

$^{1}$H-$^{13}$C -HSQC (**4**)
-OAc
NEt3
CH, CH3
CH2
f2 (ppm)
f1 (ppm)

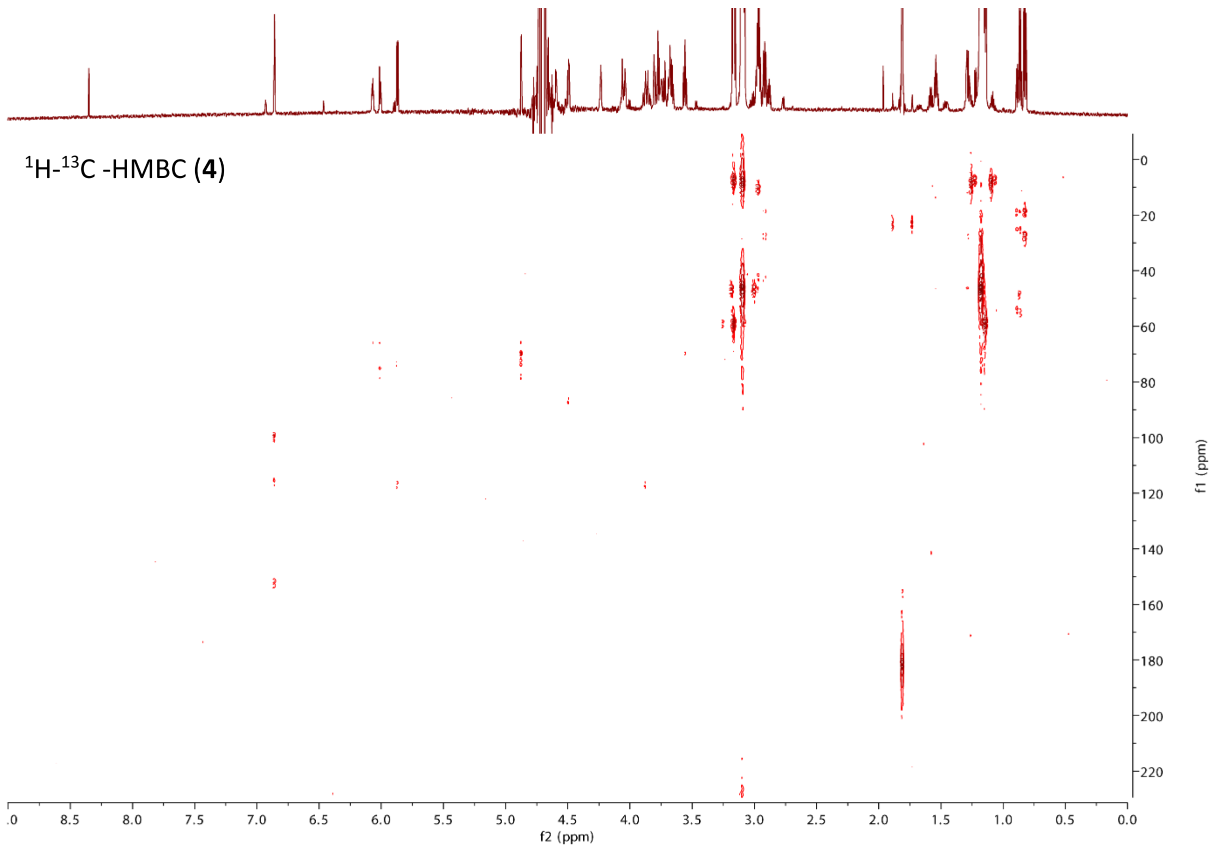

$^1$H-$^{13}$C -HMBC (**4**)
f2 (ppm)
f1 (ppm)

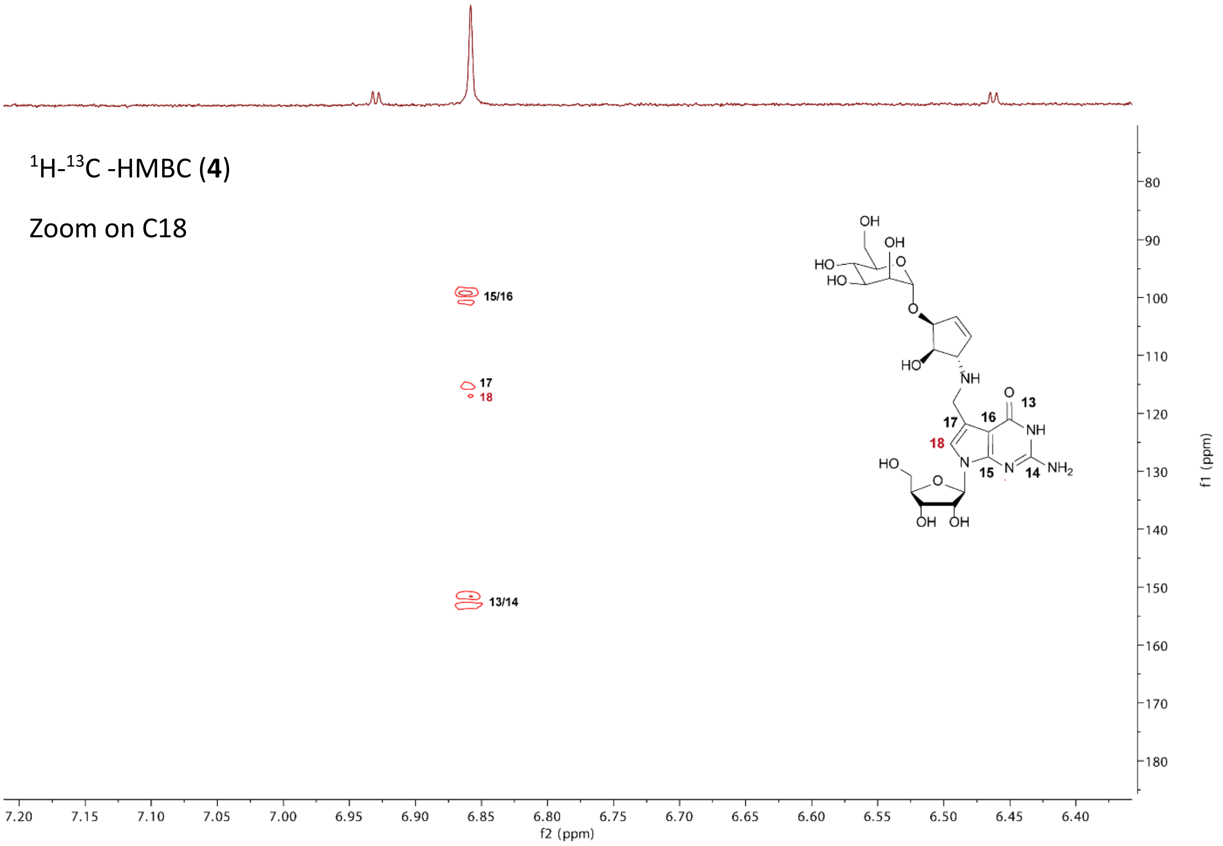

$^1$H-$^{13}$C -HMBC (**4**)
Zoom on C18
15/16
17
18
13/14
f2 (ppm)
f1 (ppm)

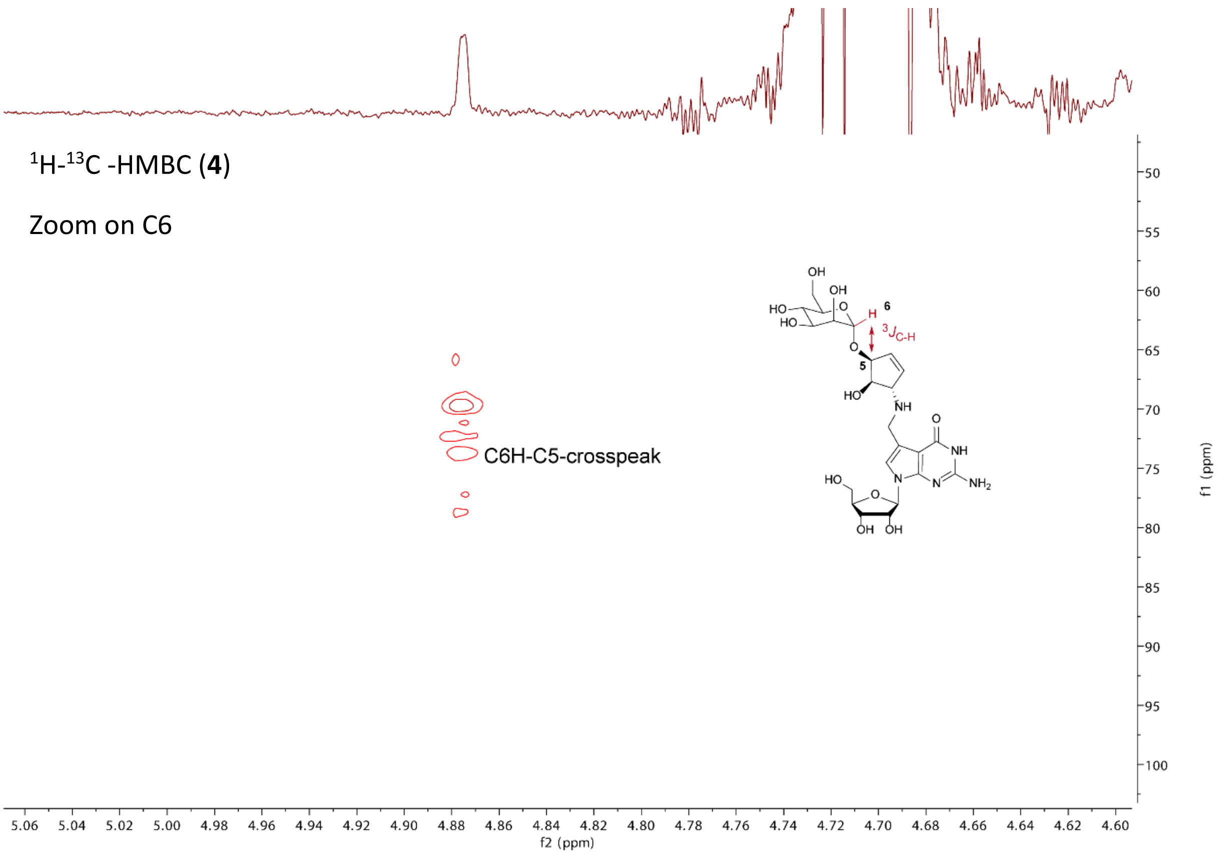

#### 5. References

1 Thumbs, P. *et al.* Synthesis of Galactosyl-Queuosine and Distribution of Hypermodified Q-Nucleosides in Mouse Tissues. *Angew. Chem. Int. Ed.* **59**, 12352-12356, (2020).

2 Kang, B. *et al.* Carbohydrate-Based Nanocarriers Exhibiting Specific Cell Targeting with Minimum Influence from the Protein Corona. *Angew. Chem. Int. Ed.* **54**, 7436-7440, (2015).

3 Krumb, M., Lucas, T. & Opatz, T. Visible Light Enables Aerobic Iodine Catalyzed Glycosylation. *Eur. J. Org. Chem.* **2019**, 4517-4521, (2019).

4 Chevalier, R. *et al.* Synthetic yeast oligomannosides as biological probes: α-d-Manp (1→3) α-d-Manp (1→2) α-d-Manp and α-d-Manp (1→3) α-d-Manp (1→2) α-d-Manp (1→2) α-d-Manp as Crohn's disease markers. *Tetrahedron* **61**, 7669-7677, (2005).

5 Crich, D. & Li, H. Synthesis of the Salmonella Type E1Core Trisaccharide as a Probe for the Generality of 1-(Benzenesulfinyl)piperidine/Triflic Anhydride Combination for Glycosidic Bond Formation from Thioglycosides. *J. Am. Chem. Soc.* **67**, 4640-4646, (2002).

6 Montenegro, J., Reina, J. & Rioboo, A. Glycosyl Aldehydes: New Scaffolds for the Synthesis of Neoglycoconjugates via Bioorthogonal Oxime Bond Formation. *Synthesis* **50**, 831-845, (2018).

7 Burgula, S., Swarts, B. M. & Guo, Z. Total Synthesis of a Glycosylphosphatidylinositol Anchor of the Human Lymphocyte CD52 Antigen. *Chem. Eur. J.* **18**, 1194-1201, (2012).
